# Spectral tuning and signaling of diverse and most red sensitive animal opsins in Mantis Shrimp eyes

**DOI:** 10.1101/2025.11.25.690411

**Authors:** Camille Brouillon, Alina Pushkarev, Marjorie A. Liénard, Songhwan Hwang, Arno Munhoven, Richard McDowell, Robert J. Lucas, Megan Porter, Han Sun, Peter Hegemann

**Author notes:** Corresponding authors: Alina Pushkarev; Marjorie A. Liénard, Peter Hegemann. These authors contributed equally to this work (Camille Brouillon, Alina Pushkarev, Marjorie A. Liénard).

## Abstract

Stomatopod crustaceans (mantis shrimps) possess extraordinary eyes with unmatched visual opsin diversity, but their potential for broad colour detection remains unclear. Here, we report that seventeen opsins, mostly from the midband eye region, are involved in colour sensing and signaling via Gq/Gi-coupling and describe the underlying mechanisms of spectral tuning. Two mid-wavelength opsins preferentially absorb blue light and a majority of long-wavelength opsins absorb in the green range, but three absorb light beyond 600 nm, representing exquisite far-red-sensitive animal opsins. Several blue and red opsins exhibit photoreversibility with opposite spectral shifts explained by retinal binding interactions, revealing unrecognised photochemical flexibility in stomatopod photoreception. Row-specific spectral sensitivity results from combining carotenoid filters with multiple individually-tuned rhodopsins, altogether extending broad vision to the far-red spectrum.

## Main Text

Mantis shrimps are stomatopod crustaceans that possess one of the most unusual and sophisticated visual systems among all known animals. The Caribbean species *Neogonodactylus oerstedii* (Fig. 1A) inhabits shallow-depth marine waters bathed with a broad spectrum of light, and its prominent highly mobile eyes bear at least 12 photoreceptor classes each tuned to specific wavelength ranges between 300 nm to 700 nm, a remarkably high number unmatched thus far in animals, and whose role in colour vision has long intrigued the scientific world (*1*).

**Fig. 1.**
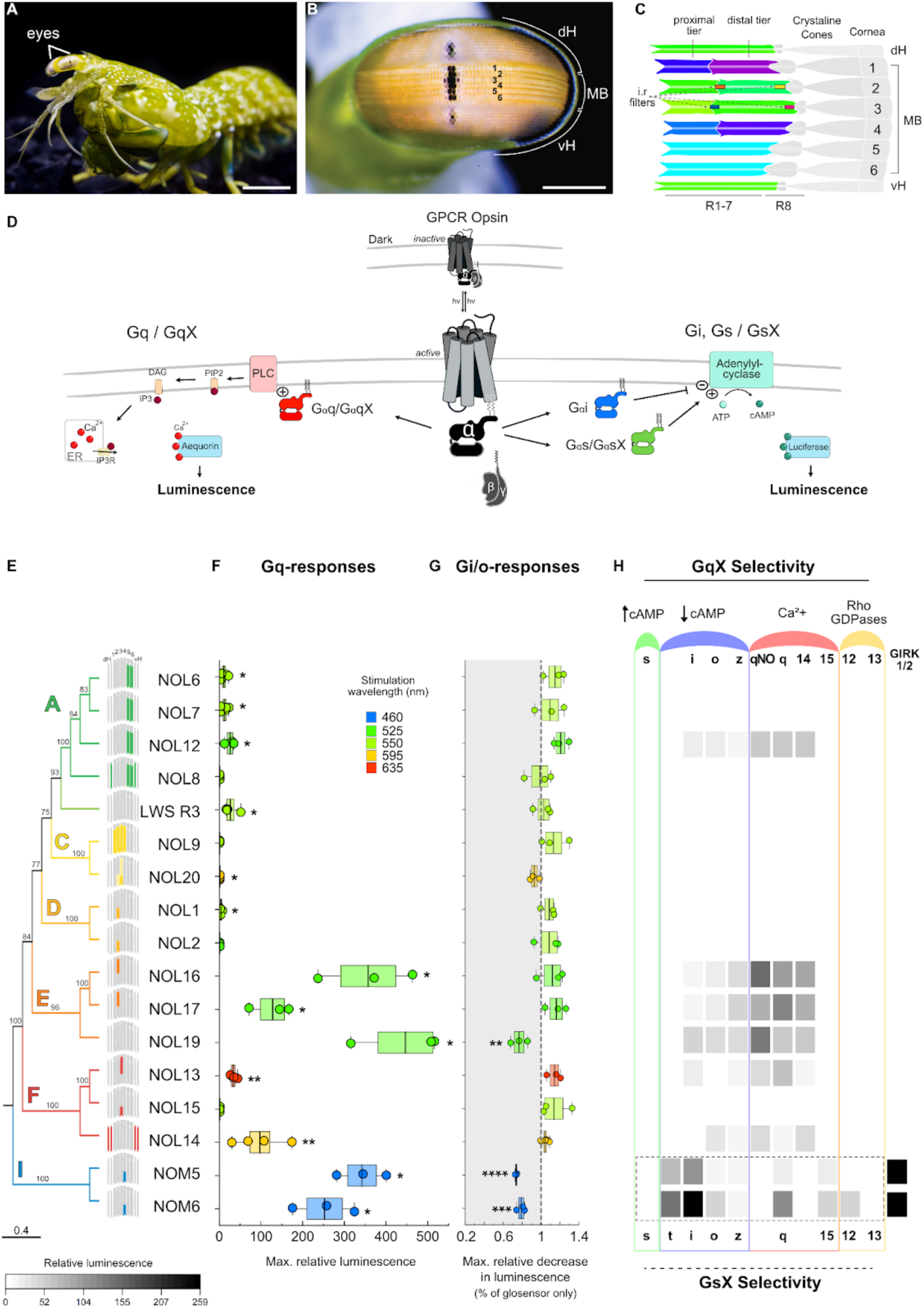
Mantis shrimp visual rhodopsins activate phototransduction through Gi and Gq signaling pathways. (**A**) Photograph of a *N. oerstedii* mantis shrimp (scale bar = 5 mm). (**B**) Close view of the external division of the compound eye composed of 6 <u>m</u>id<u>b</u>and rows (MB), and <u>d</u>orsal (dH) and <u>v</u>entral <u>h</u>emispheres (vH) (scale bar = 1mm) (from Porter et al. (2020) with permission). (**C**) Sagittal cross section diagram of the internal organisation of the retina, including intra-rhabdomal filters (i.r. filters), *i.e.* sitting within the rhabdom itself. Midband photoreceptive units 1-6 are colorized to match maximal photoreceptor absorbance as measured by microspectrophotometry and subdivided by tiers in rows 1-4 (*34*). (**D**) Scheme of bioluminescent second messenger assays with Gɑq signalling via PLC and Gɑs/Gɑi stimulating or inhibiting adenylyl cyclase (See methods for more details). (**E**) Localization patterns (scale bar = 200 μm) and evolutionary relationships from Porter et al. 2020. (**F**) Summary of maximal relative aequorin Gq-responses and (**G**) glosensor Gi/o responses (fig. S3), presented as mean ± SEM, compared to cells transfected only with luminescent reporters. Significant responses are marked with (*). (**H**) GqX- and GsX-protein coupling profiles of NOMs and NOLs.

Photoreceptor diversity arises through three distinct regions, the dorsal and ventral hemispheres devoted to spatial and motion vision, and six central rows of ommatidia (the midband or MB) that run along the equator of the eye (Fig. 1B), with rows 1-4 mediating color discrimination (*2*), whereas rows 5-6 are responsible for polarization vision (*3*). In rows 1-4, three stacked tiers of cells act as successive filtering layers, absorbing a portion of the light spectrum available to cells below. In rows 2 and 3, carotenoid-packed vesicles interspersed between the distal and proximal photoreceptive layers (*4*) considerably tune the light available to their underlying photoreceptors.

In a seminal publication, Cronin and Marshall (*5*) measured bleaching spectra from *N. oerstedii* retinal ommatidial cross-sections for all of the anatomically distinct photoreceptors absorbing within the visible spectrum. These spectra revealed absorbance maxima between 401 and 551 nm, presumed to correspond to one dominant rhodopsin (opsin with retinal chromophore) in each tier, under the assumption that all cells within a given tier expressed only a single one. The absorption spectra of the intra-rhabdomal coloured filter pigments were also measured to estimate the resulting sensitivities of midband receptors (*5*). These filters showed strong spectral overlap with their underlying photoreceptor sensitivity, an overlap that was expected to dramatically reduce photon capture by up to more than three orders of magnitude (>1000-fold) in the proximal tier 3P, thereby proposed to confer red-light selectivity in 3P, but with very low overall sensitivity, an adaptation potentially useful for the animal under bright light conditions (*6*). Wavelength dependent intracellular electrophysiological recordings (E-phys) of the UV-sensitive R8 cells in *N. oerstedii* (*7*), later complemented by measurements spanning 400 and 700 nm for a number of related shallow-and deep-water stomatopods (*1*, *8*), support that together the UV-sensitive R8 cells and the separated R1-7 tiers of ommatidial rows 1-4 in *N. oerstedii*, enable multi-chromatic sampling across the UV-visible spectrum, and spectral processing via colour-opponency and wavelength binning-mechanisms (*9*, *10*).

Early physiological evidence further suggested that the mantis shrimp eye can sense far red light and led to the proposal that red-sensitive cells in row 3 result from strong intra-rhabdomal filtering of a presumably limited opsin set (*3*, *11*, *12*), similarly to most invertebrate organisms (*13*). However, although less sensitive photoreceptors would be expected compared to other midband rows, E-phys sensitivity was remarkably similar between the red-sensitive photoreceptors in row 3 and other photoreceptor types (*3*, *11*), leaving unresolved the molecular mechanisms underlying far red sensitivity.

Most animal visual pigments absorb maximally in the visible range (*λ*_max_ 400-580 nm), and many taxa also have receptors sensitive to ultraviolet wavelengths (<400 nm) (*14*), but efficient red light-absorbing rhodopsins are remarkably rare in animals. So-called red-sensitive opsins generally maximally absorb around 560-580 nm when using an 11-*cis* A1-retinal chromophore and only the far end of the opsin absorbance spectrum reaches into the red part of the light spectrum (*15–20*). Extreme red sensitivity (>600 nm) is typically achieved either by coupling a long-wavelength sensitive (LWS) opsin to a retinylidene chromophore with an extra double bond (11-*cis* A2-retinal) (*21–25*), by addition of a bacteriochlorophyll light harvesting chromophore (*26*), or in combination with filters, such as oil droplets in birds, and carotenoid pigments in insects and crustaceans (*6*, *27*, *28*).

A key missing piece of the puzzle is how these properties build upon the unique opsin diversity recently uncovered in the eye of *N.oerstedii*, with 33 RNA-transcripts grouped into 17 distinct expression patterns and frequent co-expression across phylogenetic spectral clades (*29*). We sought to uncover how multiple specialized rhodopsins and the distribution of carotenoid pigments interact to tune colour discrimination abilities in the Mantis shrimp eye. Specifically, we set up to test whether undiscovered rhodopsins are more efficient in detecting very long-wavelengths of light to extend sensitivity into the far red.

## Three long-wavelength rhodopsins drive absorbance to the far red

To test this hypothesis while eliminating the influence of co-expressed visual pigments and filter components, we set out to fully characterize the remarkable opsin diversity *in vitro* outside the tremendous complexity of the native eye. First, we reanalyzed the genetic diversity and phylogenetic relationships (fig. S1) of long- (NOLs), and middle- (NOMs) wavelength sensitive opsin clades depending on the amino acid signature of the predicted protein retinal binding pocket. We selected and heterologously expressed the 17 opsins mostly found across the eye hemispheres and midband rows 1-4 used for colour discrimination, comprising 15 full-length NOLs, and 2 NOMs of row 4 (figs. S1-2) (*29*, *30*). After reconstitution with the A1 11-*cis*-retinal chromophore used in crustaceans and most other invertebrates (*31–33*), the NOM5 and NOM6 opsins absorbed maximally at 455 nm and 459 nm in line with microspectrophotometric (MSP) data (*34*) and their phylogenetic placement in middle wavelength sensitive clades (figs. S1,2,4). 12 NOL rhodopsins spanning phylogenetic clades A,C,D,E and F (*29*), exhibited absorbance spectra with maxima near 540 nm (fig. S4). Interestingly, NOL13 and NOL14 (clade F) as well as NOL20 (clade C) formed long-wavelength sensitive rhodopsins reliably absorbing maximally near and above 600 nm (fig. S4). These spectra appeared considerably red-shifted compared to predictions from averaged ommatidial bleaching spectra resulting from MSP recordings on *N. oerstedii* eyes (*5*), illustrating unique novel rhodopsin spectral sensitivities, potentially less expressed or whose contributions might have been previously overshadowed in the eye.

## Long-wavelength opsins couple mostly to Gq and middle-wavelengths to Gi and Gq

We next wondered if the *N. oerstedii* functional opsins were actively involved in the phototransduction cascade. We tested G-protein coupling and selective signaling preferences in each opsin using bioluminescent second messenger assays in HEK293T cells. To first test individual rhodopsins coupling to each of the three main families of human G-proteins, we tested the relative activation of Gq, Gi/o and Gs (*35*) (Fig. 1D-E, fig. S4). Five NOL rhodopsins (2, 8, 9, 15 and 20) did not elicit measurable responses to any G-protein type compared with cells transfected only with aequorin or glosensor, despite robust protein expression levels at the plasma membrane (fig. S3), suggesting that they do not couple to human G-proteins. Ten NOL rhodopsins (1, 6, 7, 12, 13, 14, 16, 17, 19, 20), LWSR3, as well as NOM5 and NOM6 coupled significantly to Gq (p<0.04, Fig. 1E, fig. S5, Table S1), although signal intensity between rhodopsins may not be directly comparable, notably owing to different expression levels, membrane targeting, and G-protein coupling efficiencies. We could not detect activation of Gi or Gs for any NOL rhodopsin except for NOL19, however NOM5 and NOM6 reacted readily with Gi (p< 0.001, Fig. 1E, fig. S5, Table S1).

We further investigated the G-protein selectivity profiles by co-expressing *N. oerstedii* opsins in HEK293 cells that lack endogenous G-proteins (*36*), with individual human GsX chimeric proteins where the rhodopsin-interacting helix had been exchanged for helices of other G-subtypes (*37*). Again, NOM5 and NOM6 both exhibited higher G-protein coupling to Gsi than to Gst, whereas NOM5 had lower responses to GsX chimeras of the Gq family compared to NOM6 (Fig. 1F, fig. S5). As most of the signaling NOLs did not couple efficiently to GsX proteins but demonstrated significant Gq-responses, human GqX hybrids (*38*, *39*) were used following the same principle of the GsX assay (Fig. 1F). The Gq signal was quantifiable with enough variability to allow a reliable discrimination between GqX responses from six NOLs (NOL12, 13, 14, 16, 17 and 19) (Fig. 1F). All six characterised NOLs exhibited higher selectivity to GqX proteins of the Gq family, i.e. Gq or Gq14, but not Gq15 (Fig. 1F, fig. S5). In addition, all six NOLs showed weaker coupling to the Gi/o family of GqX proteins, with NOL14 in particular exhibiting comparatively high coupling to Gqo (21.9% of the sum of the GqX-protein selectivity responses, fig. S5). None coupled significantly to GqX components of the G12/13 or Gs families. In particular, NOL13 activated preferentially in presence of Gq (43%) and Gqi (9.6%), and significantly less with Gqz and Gqo (Fig. 1F, fig. S5). We also tested the only so far reported native *N.oerstedii* Gq-protein (GqNO) putatively involved in phototransduction (*30*) and found that all six NOL Gq-rhodopsins also activate Mantis Shrimp GqNO (Fig. 1F) but not with better signaling than for human Gq, except for NOL16 and NOL19 (both 37% to GqNO and 17% to Gq, fig. S5D-E). Due to the strong Gq responses, we measured the wavelength dependence of Gq activation for NOL12, NOL16, NOL17 and NOL19, four green-sensitive opsins expressed across rows 2, 5 and 6. We constructed individual action spectra that closely resembled the *in vivo* rhodopsin absorption profiles (*5*, *40*, *41*) (fig. S4), overall aligning both approaches to confirm these four rhodopsins’ green-sensitivity. Overall, not including that some unresponsive NOLs may couple to other potential native GqNO-proteins, these results identify at least 13 functional opsins actively coupling to G-proteins thereby parsimoniously contributing to phototransduction in the mantis shrimp eye.

## Four photoreversible rhodopsins may modulate fast wavelength sampling

Owing to its shallow water habitat and visual ecology, *N. oerstedii* is subject to fluctuating light and colour distribution, which encouraged us to study the signalling specificity. To examine this question experimentally, we leveraged our Gq assay with the eight most promising, Gq-active rhodopsins (6 NOLs, 2 NOMs), and implemented double pulse experiments to examine their photoreversibility properties, by delivering a first pulse of light for Gq activation, and a second pulse for deactivation, and quantifying rhodopsin conversion back to their original D-state (Fig. 2A) (*42*, *43*).

**Fig. 2.**
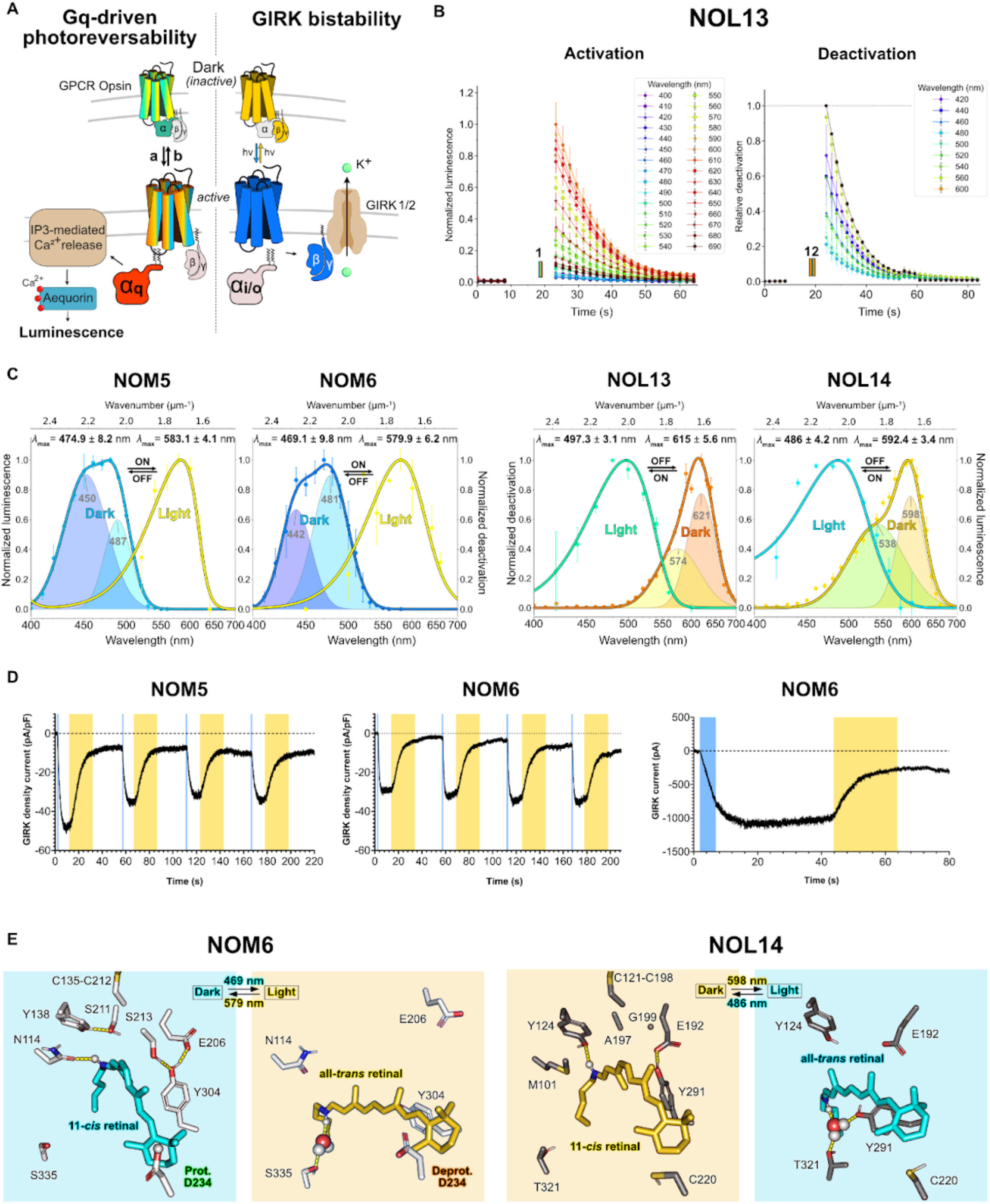
Photoreversible *N.oerstedii* rhodopsins have opposite spectral properties explained by distinct retinal schiff base microenvironments. (**A**) Activation and deactivation of the Gq pathway (*left*) and the GIRK1/2 channel (*right*) by a Gq- and Gi/o-coupled GPCR opsin, respectively. (**B**) Wavelength-dependent aequorin luminescent activation/deactivation responses of NOL13. See SI for other Gq-activated rhodopsins. (*left*) Activation is induced by a 1s light pulse (colored rectangle 1) of fixed wavelengths (1x10^16^ photons cm^-2^ s^-1^,n=3 experiments). (*right*) Deactivation is assayed by two consecutive 1s light pulses, one with the fixed wavelength leading to maximal activation (orange rectangle 1, 600 nm, 1x10^16^ photons cm^-2^ s^-1^) followed by a second light pulse of varying wavelength (colored rectangle 2, 1x10^16^ photons cm^-²^ s^-1^, n=3). (**C**) Activation and deactivation spectra of NOM5, NOM6, NOL13 and NOL14, derived from peak aequorin responses and fitted with a non-linear least squares optimization of the Govardovskii visual template . Dark spectra are fitted with a two component Gaussian Mixture Model (GMM) (Table S1 and fig. S6) Data is presented as mean values ± SEM. X-axis is a linear energy scale (µm^-1^) corresponding to a log wavelength scale (nm). (**D**) GIRK photocurrents induced by the activation of NOM5 and NOM6 (1s 470 nm light pulse, 1x10^14^ photons cm^-2^ s^-1^, blue line) and deactivated by a subsequent 20s light pulse (595nm, 4x10^15^ photons cm^-2^ s^-1^yellow bar). (**E**) Predicted structural models of the inactive NOM6 and NOL14 with 11-*cis* Retinal Schiff Base (RSB) and active (all-*trans* RSB bound) states of NOM6 and NOL14. Key hydrogen-bonding interactions are shown as yellow dashed lines and water molecules are represented as spheres.

Wavelength-dependent activation and deactivation spectra recorded for aequorin luminescent responses of NOM5, NOM6, NOL13, and NOL14 are presented in Fig. 2B,C. We first activated NOM5 and NOM6 with a 1s-light pulse of fixed wavelength at near-saturating light intensity (450 nm) and tested their Gq responses after a second red-shifted 1s-light pulse of varying wavelengths. We found a maximal reduction of 28.1% of the initial response at 583 nm for NOM5 and 21.58% at 580 nm for NOM6 (Fig. 2C). Plotting the inverse of this reduction against wavelength reveals the active Meta (L-) state spectra of the NOM photopigments which are photoreverted to the dark (D-) state with the second light pulse (Fig. 2C). The dark activation spectra presented a dual spectral composition that was best fitted with a two-component Gaussian Model, suggesting a potentially pH-dependent heterogeneity of the protein (Fig. 2C, Table S2, fig. S6).

Similarly, NOL13 and NOL14 activation responses were acquired with a 1s-pulse fixed at 600 nm, and thereafter their Gq-response reduction was quantified after blue-shifting the second light pulse, leading to maximal deactivation (75.1%) at 497 nm for NOL13, and reduced NOL14 response up to 55.7% at 486 nm. Visual fitting characteristics of the dark state also indicated the contribution of two spectral isoforms (Fig. 2C, fig. S6). In contrast, for NOL12, 16, 17 and 19, no second light pulse could change the response to the first, suggesting that these rhodopsins are not photoreversible or have closely overlapping spectra in the dark and photo-activated states.

Finally, bistability and kinetic properties of NOM5 and NOM6 were further investigated electrically using the well established G protein-gated inwardly-rectifying potassium channel GIRK1/2, which canonically activates upon binding of the Gβγ subunit of Gi/o-proteins (*44*) (Fig. 2A). Importantly, the Gi-responsive NOM5 and NOM6 could repetitively activate and deactivate GIRK1/2, with blue and orange light respectively, and with a limited loss in current amplitude, confirming that these opsins are thermally photoreversible and bistable (Fig. 2D). Of note, even after using a 20 second pulse for deactivation at maximal light intensity (20s, 4x10^15^ photons cm^-^² s^-1^), GIRK photocurrents could never completely recover to the pre-illumination baseline, likely due to the still overlapping dark and photoproduct spectra at the absorbance tails and the high light intensity required to revert the opsins to the dark state. Nevertheless, the photoproduct spectra of all four rhodopsins, which are considered as signaling states, were well separated from the dark state spectra (Fig. 2C). GIRK currents indicated kinetics of light-induced activation-deactivation responses and the thermal stability of the signaling state for NOM5 and NOM6 (Fig. 2D).

Thus, NOM5, NOM6, NOL13 and NOL14 are not only stable after photoexcitation without bleaching, they are also capable of returning to their inactive dark state photochemically due to the weak overlap of the D- and L-spectra.

## Inverse spectral tuning of NOM6 and NOL14 arises from specific retinal binding pocket properties

To gain a better understanding of the pronounced spectral differences between the dark states (D) and photoproducts (L), as well as the inverted photochemical shifts observed between NOM6 and NOL14, we generated structural models of the retinal binding pockets while maintaining the retinylidene chromophore protonated (Fig. 2E, fig. S7). The structural transitions from 11-*cis* retinal in the D-state to all-*trans* retinal in the L-state were modeled by combining AlphaFold3 predictions with multiple sequence alignment (MSA) manipulation (*45*, *46*), molecular docking, all-atom molecular dynamics (MD), hybrid quantum mechanics / molecular mechanics (QM/MM) simulations, and vertical excitation energy calculations.

The predicted 7TM helix structure models exhibit high overall confidence, with predicted local distance difference test (pLDDT) values mostly above 80, and high structural stability during equilibrium MD simulations (figs. S7-8-9). The counterions E206 and E192 are positioned unusually distant from the retinal in both proteins, only NOM6 features a unique acidic residue (D234) near the β-ionone, which is replaced by a cysteine (C220) in NOL14.

Comparison of the predicted absorption spectra for different protonation states of D234 in NOM6 revealed that a protonated D-state and a deprotonated L-state showed the closest agreement with the experimental data (fig. S10) in consistency with the original double-point charge model proposed for rhodopsins (*47*, *48*), in which a negative charge near the β-ionone electrostatically stabilises the positive charge that formed in this region upon retinal photoexcitation causing a strong red shift. Next, we sought to rationalise the inverse spectral shifts observed between NOM6 and NOL14 by comparing their QM-optimized structural models. Notably, in NOM6 the residues flanking the disulfide bond are polar (S211 and S213) and S211 forms a hydrogen bond with Y138, while S213 interacts with Y304 and indirectly the counterion E206. The S211-Y138 interaction redirects the N-H of the retinal to N114 increasing polarity at the retinal nitrogen (fig. S11). In contrast, in NOL14, the disulfide neighbors are nonpolar (A197 and G199). And the *11-cis* retinal N-H occasionally binds with Y124, while Y291 interacts with the counterion E192 only. The resulting reduced retinal distortion results in greater π-conjugation and a red shift, compared to the distorted C12-C15 bonds in NOM6 (fig. S12). The all-*trans* retinylidene N-H group in NOL14 forms a water-mediated hydrogen bond with T321, and occasionally to Y291. This configuration electrostatically stabilizes the ground state, increasing the excitation energy and causing a blue shift while the deprotonated D234 near the ring in NOM6 exerts the opposite effect, inducing a red shift. In conclusion, we propose that the inverse spectral shifts observed between NOM6 and NOL14 arise from substantial differences in the polarity and hydrogen-bonding networks within their retinal binding pockets. Such variations modulate the extent of π-conjugation and geometric distortion of the retinal, thereby fine-tuning its absorption characteristics.

## Modeling the tuning of rhodopsin sensitivity with and without filtering pigments partitions photoreceptor spectral specialization

Next, we investigated how the multiple rhodopsins of the eye hemispheres, and midband ommatidial tiers tune photosensitivity without or under the influence of inter-rhabdomal filtering pigments, respectively. In the lateral dorsal and ventral hemispheres (DH and VH) devoid of inter-rhabdomal filtering, NOL14 with *λ*_max_ = 592 nm is expressed in conjunction with NOL8 with *λ*_max_ = 540 nm. Together they confer an unfiltered broad sensitivity (fig. S13), which is at least half maximal between 470 to 660 nm. This offers the dorsal and ventral sides of the mantis shrimp eye an exceptional ability to perceive from blue to very long-wavelength incoming terrestrial red light, the latter possibly persisting at very shallow depths, especially at dusk and dawn (*49*, *11*).

We next calculated the resulting spectral sensitivity of MB tier 2D, where NOL16, NOL17 and NOL9 (*λ*_max_: 521, 537, and 541 nm, respectively) are expressed in conjunction with F1 filters (R2F1) (Fig. 3, Fig. 4A). NOL9 is present in all midband rows 1-4 (*29*), and presumably less likely to contribute to major differences in tier photoreceptor sensitivity. By taking into account the reported absorption spectra of R2F1 (*5*), our results provide evidence that the maximal sensitivities of NOL16 and 17 are reduced by the R2F1 filter to roughly 45% and shifted bathochromically by 16 - 18 nm (Fig. 4B-C, fig. S13). In addition, R2F1 narrows half bandwidth (HBW) from 100 nm to 60 - 80 nm, significantly increasing spectral discrimination in row 2 without great sensitivity loss (fig. S13).

**Fig. 3.**
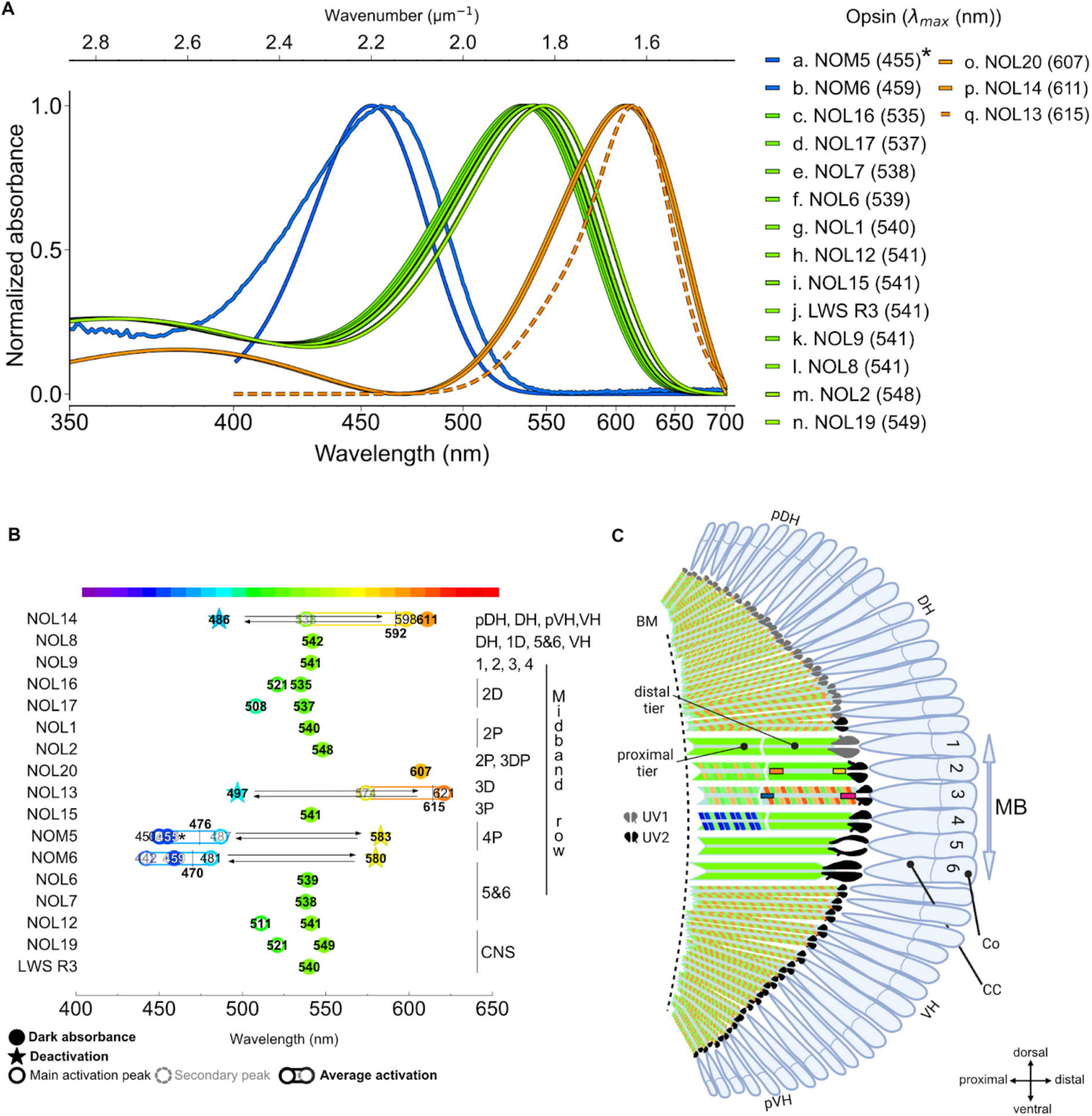
Multiple specialized spectral and photoreversible rhodopsins contribute to sharp, finely-tuned photoreceptor properties in the mantis shrimp retina. (**A**) Spectral fits dark NOL and NOMs absorption with 11-*cis* retinal or 9-*cis* retinal (*) full lines. For NOL13 the action spectrum is plotted in a dashed line (see fig. S4, Fig. 2). (**B**) Absorbance maxima of mid-and long-wavelength rhodopsins characterised in this study, arranged by localization in different regions including eye hemispheres, eye midband rows 1 to 6, or the central nervous system (CNS). DH, dorsal hemisphere; VH, ventral hemisphere; p, peripheral; D, distal tier; P, proximal tier. Numbers indicate best-fit estimates of λ_max_ from dark absorbance (full circles), plus activation (empty circles), and deactivation spectra (stars) when measurable. Maximal activation for NOL13, NOL14, NOM5 and NOM6 is reported as the peak of the Gaussian mixture (main individual peak in empty circle and secondary individual peak in light grey and dashed circles) in bold, circled by an ellipse (see Table S2), Bidirectional arrows indicate that NOM5, NOM6, NOL13 and NOL14 can be reversibly switched bidirectionally between dark and active states. (**C**) Sagittal cross section diagram of the eye. The location and representative absorption range of the multiple rhodopsins expressed in each region are mapped as coloured diagonal cross patterns according to the number of rhodopsins expressed in this study and their absorption. Intra-rhabdomal filters in rows 2 and 3 of the midband row (MB) are depicted by vertical rectangles. Co, cornea; CC, crystalline cone; BM, basal membrane.

**Fig. 4.**
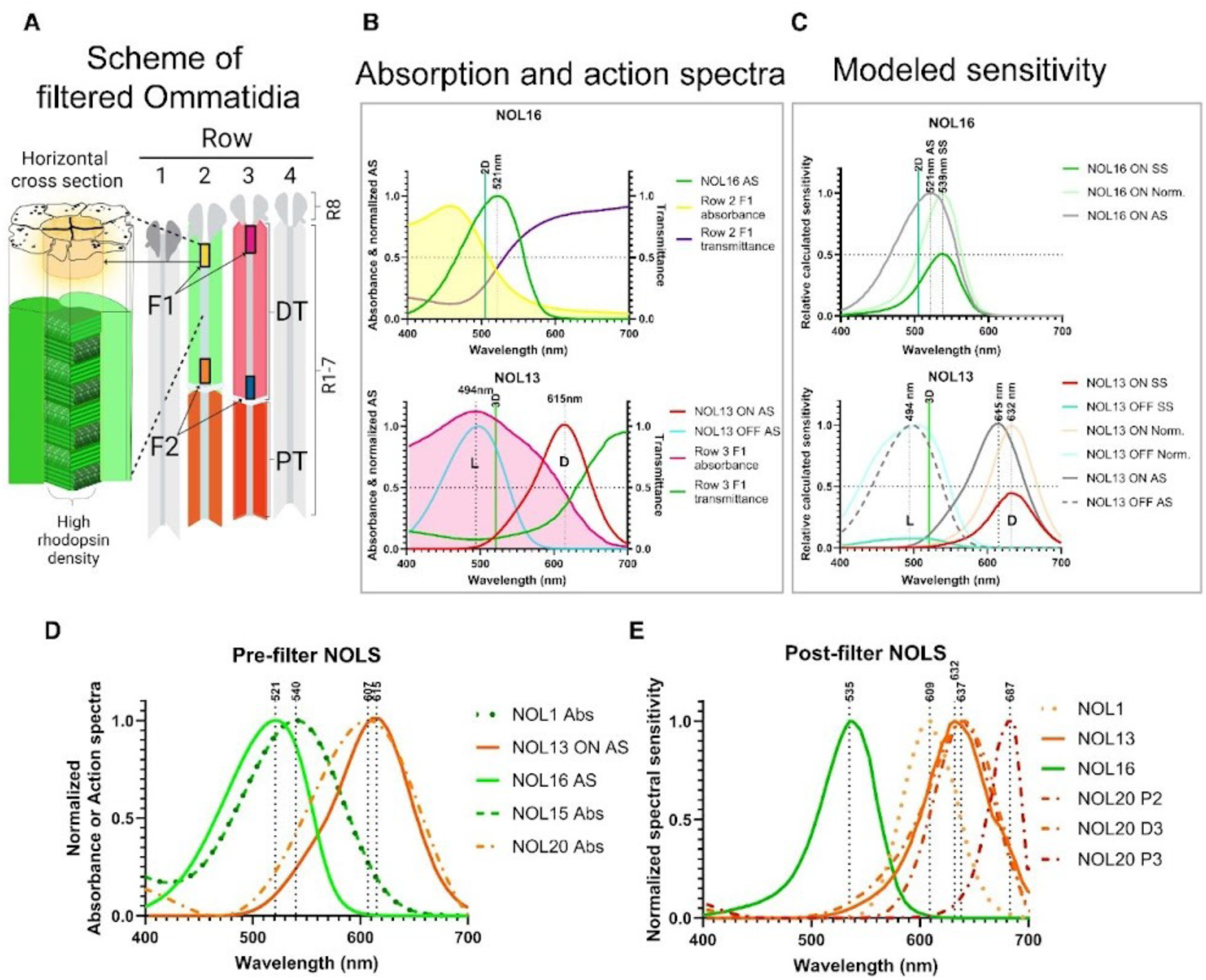
Long-wavelength rhodopsins of midband rows 2 and 3 are fine-tuned by intra-rhabdomal filters, extending sensitivity to the far red. (**A**) Scheme of midband rows 2 and 3 showing the intra-rhabdomal carotenoid filters (F1 and F2) and the microvillar organisation. DT, distal tier; PT, proximal tier. Tiers in rows 2 and 3 are coloured according to the spectral sensitivity obtained with quantitative modeling of rhodopsin-filtering. (**B**) Filter absorbance and transmittance (*5*), and wavelength absorbance of NOL16 (upper panel) and NOL13 (lower panel) of row 2 and 3 are plotted against wavelength. MSP peak absorbance of the distal tiers indicated by the vertical line in turquoise (2D) and green (3D) (*5*). L, light (ON); D, dark (OFF); AS, action spectra. See also fig. S13. (**C**) Calculated spectral sensitivity for NOL16 (upper panel) and NOL13 (lower panel) result from transmittance of the F1 filter multiplied by normalized action spectra of NOL16 and NOL13, respectively. Faint lines show the normalized sensitivities. For NOL13, activation and deactivation spectra are plotted in red and blue, respectively, and the action spectra is plotted in grey. Photoreceptor sensitivities and filtering in the UV are neglected. Half bandwidth is indicated with a bold horizontal dashed line. Norm., normalized; SS, spectral sensitivity. Sensitivities of individual additional rhodopsins are presented in fig. S13. (**D-E**) Normalized spectral sensitivities of tiers D2, P2, D3 and P3. (**D**) Original rhodopsin spectra prior to i.r. filtering. (**E**) Spectral sensitivity post-filtering. Maximal peak sensitivities are marked by dotted vertical lines. Post-filtering sensitivity below 0.1%, was not plotted (e.g. NOL15). AS, Action spectra; Abs, absorbance spectrum. Optical densities are reported in fig. S13.

In MB tier 3D, comprising NOL9, NOL20 and NOL13, the F1 filter (R3F1) absorbs further to the red end of the visible spectrum compared to R2F1 (Fig. 4B, fig. S13). Therefore, in the dark, the spectral sensitivity of NOL13 and NOL20, both red shifted in absorption compared to NOL16, are still screened out by R2F1. Likewise, the resulting NOL13 and NOL20 opsin sensitivities in 3D are shifted bathochromatically by 16 nm, respectively achieving 632 nm for NOL13 and 637 nm for NOL20, on the expense of a final moderate reduction of sensitivity to 45 - 49% (Fig. 4C,E, fig. S13D). Since the absorption maxima of NOL16 and NOL13 are at a wavelength where transmission of the F1 filters changes the most (fig. S13D), it causes compression of the HBW and shifts the sensitivity peaks of the rhodopsins with only moderate sensitivity loss (fig. S13C-D).

Microspectrometric data analysis in *N. oerstedii* predicted 560 nm as peak sensitivity for 3D (*6*), whereas electrical spectral sensitivity measurements for row 3D of two related stomatopods, *Gonodactylus smithii* and *Haptosquilla trispinosa,* placed *λ*_max_ at 610 nm (*8*). Our calculations show that modulation of photoreceptor red light sensitivity in 3D falls above the previously proposed amplitude range for peak sensitivity with 632 nm for NOL13 and 637 nm for NOL20 in 3D (Fig. 4), demonstrating the dual role of R3F1 filtering and the multiple rhodopsin sensitivities. Reversely, we find that rhodopsins with absorption maxima near the F1 maxima are completely shielded with their sensitivity reduced over the whole spectrum accordingly. This is the case of the NOL13 photoproduct (NOL13-L, *λ*_max_= 497 nm) for which sensitivity is reduced at its maximum to 7.6%, proportional to the low R3F1 filter transmission at these wavelengths (Fig. 4B-C).

In MB tiers P2 and P3, sensitivity calculations account for the reduction in light spectrum available after absorption in the distal tiers, and the high optical densities of the proximal filters (A = 7.7 for R2F2, A = 3.6 for R3F2 (*5*)) that further dramatically reduce the light captured by the stacked proximal tier rhodopsins. In doing so, we show that in P2, R2F2 filtering of NOL1 (*λ*_max_ = 540 nm) and NOL20 (*λ*_max_ = 607 nm) reduces their maximal sensitivities to 2.8% and 20%, respectively, and shifts the narrowed maxima to 609 nm and 641 nm (fig. S13D). In P3, R3F2 filtering of NOL15 shields out the rhodopsin sensitivity even further to < 0.1 % due to strongly overlapping absorbance maxima between the rhodopsin and the filter, whereas filtering of NOL20 shifts the tier sensitivity maxima to 687 nm with a reduction to ca 2.5% (fig. S13). This interplay acts as the mechanism to achieve *in vivo* far-red sensitivity in row 3 in the *N. oerstedii eye*.

## Trade-off photoreceptor adaptations between spectral reach and sensitivity allow vision at the far red edge

Our findings show that the Mantis Shrimp *N. oerstedii* uses a subset of long-wavelength opsins (NOL13, 14 and 20) that absorb light maximally near or above 600 nm (Fig. 3A, fig. S13), achieving real far-red absorbing properties by sequence modification while still coupling to the universal A1 11-*cis*-retinal chromophore. In the midband rows 2 and 3 specialized to colour vision, light of the red edge of the rhodopsin absorption spectrum can be effectively used by the employment of selective filters that absorb shorter wavelengths of light through red carotenoid pigments (Fig. 3, fig. S13) (*50*, *5*, *4*, *51*). This mechanism confers *N. oerstedii* with exceptional long-wavelength (far red) vision, but reduces absolute sensitivity. Hence, the magnitude of the resulting shift in effective spectral sensitivity depends on the degree of overlap between the filter’s absorbance spectrum, the rhodopsin absorbance, and on the filter’s optical density. High optical density and asymmetric filter-rhodopsin absorbance overlap produce large shifts (e.g. NOL1, NOL13, NOL20), whereas well-aligned opsin-filter absorption will cause a loss in photon catch and attenuate light sensitivity of the corresponding tier with little or no shift (NOL15) (Fig. 4, fig. S13). In the proximal (P) tiers of colour-vision rows 2 and 3 (P2 and P3), the intra-rhabdomal filters are particularly optically dense, leading to substantial spectral shift reaching up to 80 nm but sacrificing sensitivity down to < 0,1%. Consequently, far red vision is likely restricted to bright daylight, when using P2 and P3 photoreceptors, after the distal D2 and D3 rhodopsins are already inactivated in their P-state. In contrast, at lower light levels, during dusk and dawn, colour vision relies mainly on D2 and D3 that provide a less extreme, but a more sensitive spectral range.

Overall, in *N. oerstedii*, the distal tiers of the midband ommatidia with high sensitivity are designed for low light intensities, whereas the proximal tiers are the main actors at bright light with narrowed but strongly reduced light sensitivities and a far-red maximum near 690 nm.

Mantis shrimps can therefore sample far-red wavelengths more specifically than most other animals and possess the visual capacity to potentially exploit far-red chromatic and achromatic signal reflectance (total luminance), although its contribution in an ecological context is untested behaviourally. For communication, *N. oerstedii* reliably assesses total UV reflectance of competitor’s meral spots (*52*, *53*), which also contribute up to 10% far red light between 700 and 800 nm (*54*, *55*). This suggests that its eye far red sensing photoreceptors may add a signaling channel in specific ecological and visual contexts, alone or combined with UV cues or fluorescent enhancement, or at high sun angles when UV and visible spectral channels are relatively bleached and filtered out (*53*, *56*).

## Spectrally-separated rhodopsin states in the visible spectrum likely contribute to dynamic temporal signaling and colour recognition

The photoreversibility properties of at least four mantis shrimp rhodopsins are unique in the sense that they contrast to the monostable vertebrate cone rhodopsins (*27*), which loose the retinal after photoactivation (*57–59*), to the bistable invertebrate parapinopsins (*60*) or *Platynereis* opsins (PdCO) (*61*) where one state absorbs in the visible and the other in the UV, and to the squid and jumping spider rhodopsins (*62*) where both D-state and L-state absorb in the visible but with strongly overlapping spectra (*42*, *63*, *64*).

Interestingly, in Mantis Shrimp, photoreversibility with large spectral separation of the D- and L-states without chromophore deprotonation seems to be achieved by a unique hydrogen-bonding network that fine-tunes the electronic environment and geometric distortion of the chromophore, along with large modulation upon photoisomerization, without alteration of the RSB pKa. The spectrally-separated D and L states may be an ingenious evolutionary solution to increase rapid spectral sampling between contrasting colours in natural contexts.

Collectively, the results of our current study show that the finely tuned visual system of the mantis shrimp *N. oerstedii* achieves exceptional colour discrimination notably via a diverse functional repertoire of middle and long-wavelength rhodopsins, some of which evolved photoreversibility and far-red absorption properties. Photoreversible middle and long-wavelength rhodopsin states are expected to contribute to rapid contrast perception and spectral tuning, whereas far red rhodopsins act in concert with carotenoid filtering to approach maximal sensitivity near 690 nm. Low density filters of the distal areas shift the spectra and reduce the sensitivity less and should function in dim light whereas the high density filters of the proximal tiers shift more by sacrificing sensitivity which is affordable in bright light, illustrating a series of evolutionary innovations distinct from typical animal rhodopsins, and that may carry highly beneficial adaptations in bright, spectrally broad and complex underwater environments.

## Acknowledgments

The authors would like to thank Antoinette Nguyen for technical assistance, Thomas Cronin and Michael Bok for stimulating discussion and Sonja Kleinlogel for discussion at a very early stage of the project. We thank Lisa Neuhold and Clint Makino (National Eye Institute) for sending us 11-cis retinal, Asuka Inoue for sharing the GqX plasmids, Stefan Herlitze for the HEK-GIRK1/2 cells, and Peter Soba for the HEKΔ7. We thank Dr. Tillmann Utesch for helpful discussions. The authors gratefully acknowledge the computing resources provided by the NHR Center (NHR@ZIB) through the high-performance computer “Lise” as well as by the Erlangen National High Performance Computing Center (NHR@FAU).

## Funding

Hertie Foundation (PH)

Rothschild Fellowship from Yad Hanadiv (AP)

Christiane Nüsslein Volhard Stiftung (AP)

Hector Fellow Academy (AM)

European Molecular Biology Organization Long-Term Fellowship ALTF-168-2021 (AP)

Swedish Research Council VR-2020-0517 (MAL)

Belgian Fonds de la Recherche Scientifique-FNRS F.6002.24 (MAL)

German Research Foundation SPP426019636 (PH)

European Research Council Synergy “SOL” 951644 (PH, RJL).

## Author contributions

Conceptualization: PH, AP, MAL

Methodology: PH, MAL, MP, AP, CB, HS

Investigation: CB, AP, MAL, SH

Visualization: CB, AP, SH, MP

Funding acquisition: PH, RJL, MAL, AP

Project administration: PH, MAL

Supervision: PH, RJL, HS, AP

Writing – original draft: AP, MAL, CB, PH

Writing – review & editing: CB, AP, MAL, SH, AM, RMD, RJL, MP, HS, PH

## Data and materials availability

All data are available in the main text or the supplementary materials.

## Materials and Methods

### Phylogenetic analyses

*N. oerstedii* opsin DNA sequences were obtained from NCBI and reanalysed in Geneious to exclude partial sequences (Supp. Dataset S1). Full-length opsins were selected if they met further criteria: belonging to the proposed long-wave (NOL) or middle-wave (NOM) clades and being expressed either in the colour absorbing midband rows, the lateral hemisphere of the mantis shrimp eye (NOL14), as well as in the eye stalk (LWS R3). Deduced protein sequences for the 17 selected opsins (14 NOLs, 2 NOMs and LWS R3) were aligned in ClustalX (*65*), and the phylogeny was reconstructed as reported earlier (*29*).

### Opsin cloning and expression constructs

Opsin ORF sequences from *N.oersterdii* were codon-optimized for mammalian expression, synthetised by Genscript and subcloned into various expression vector constructs as described in fig. S2a, using PCR amplification followed by restriction digestion and T4 ligation or Gibson assembly (*66*). The coding sequence for the Gαq-protein from *N. oerstedii* was codon-optimized and subcloned into the pCAGGS plasmid vector. Gqx plasmids were obtained from Asuka Inoue (Tohoku University, Japan). All final plasmids were sequence-verified by Sanger Sequencing, and endotoxin free plasmid stocks were prepared using the Zymo Midi-prep procedure (Zymo Research) prior to transfection.

### Cell culture

HEK293T cells (Life technologies) were used for Gq, Gi/o and Gs live bioluminescent second messenger assays and HEK293 lacking GNAS/GNAL/GNAQ/GNA11/GNA12/GNA13/GNAZ (HEK293ΔG7) (*67*) a gift from P. Soba, Friedrich-Alexander-Universität, Erlangen) were used for testing more specifically the Gα signaling selectivity in GsX and GqX assays.

Briefly, cells were grown at 37°C and 5% CO_2_ in Dulbecco’s modified eagle medium (DMEM) supplemented with 10% fetal calf serum (FCS, Biochrome) and 1% penicillin/streptomycin (100µgml−1, Biochrome). Two days before measurements, 5 x 10^5^ cells/well were seeded in 12-well plates (Sigma-Aldrich). Cells were transiently transfected with a 1:1 ratio of Glo22F luciferase (GloSensor, Promega) for Gi/o, Gs and GsX assays, or aequorin (gift from R. J. Lucas) for Gq and GqX assays, together with opsin plasmid (500 ng/well of each plasmid) using FuGENE HD (Promega) according to the manufacturer’s instructions. For GsX and GqX assays, HEK293ΔG7 were also co-transfected with 10ng of individual GsX or GqX chimeras (50:50:1 ratio of GloSensor or Aequorin to opsin to G-protein chimera plasmid).

After transfection, all steps were carried out under red dim light. Cells were incubated for 3-7h at 37°C and subsequently resuspended using Trypsin-EDTA (0.05 %, Gibco Thermo Scientific) and transferred to 96 white well-plates (BRAND) in DMEM with 10µM 11-*cis* retinal (gift from Lisa Neuhold and Clint Markino, National Eye Institute, National Institutes of Health). At least 12 h after resuspension, the medium was exchanged with transparent Leibovitz’s L-15 medium with 10% FCS. For Gq and GqX assays, L-15 medium was supplemented with Coelenterazine-h (10µM, Promega) and cells were incubated for 2h at Room Temperature (RT). For Gs, Gi/o and GsX assays cells were incubated in L-15 medium with beetle luciferin (2 mM in 10 mM HEPES pH 6.9, Promega) for 1h at RT.

### NOL opsin purification for dark absorbance spectra measurements

HEK293T cells were cultured in DMEM with high glucose (GIBCO) supplemented with 10% fetal bovine serum (FBS, Life Technologies) without antibiotics. For each purification, twenty 10-cm dishes were seeded with HEK293T cells at 2M on day 1 and transfected 24 hours later with lipid complexes formed in serum-reduced Opti-MEM (Life Technologies) with 24 µg Endotoxin-free plasmid DNA (pcDNA5-FLAG-T2A-Sirius, fig. S2a) and 72 µL Polyethylenimine (1 mg/mL in pure water; PEI 25,000MW, PolySciences). 11-*cis*-retinal was added to plates at 5µM (in 100% Ethanol) under dim red light illumination 6h post transfection together with fresh culture medium, then plates were wrapped in aluminium and placed back in the incubator. One transfected plate was wrapped separately as control for fluorescence visualisation. Forty eight hours after transfection, the medium was discarded, cells were scraped and collected in a a black tube in a total volume of 50 mL Hepes cold resuspension buffer containing 3 mM MgCl_2_, 50 mM Hepes pH8.5, 140 mM NaCl and EDTA-free protein inhibitors (Sigma-Aldrich), washed, centrifuged for 10 min at 4°C and 1,620 x *g* and resuspended in 10 ml cold resuspension buffer then nutated for 90 min with 40 µM 11-*cis*-retinal and 10 rpm rotation. Following ultracentrifugation at 21,500 rpm for 25 min, the total membrane fraction was extracted in cold solubilization Hepes buffer (3 mM MgCl_2_, 50 mM Hepes pH8.5, 140 mM NaCl) supplemented with 1% DDM and 25% Glycerol vol/vol for 1h at 4°C with 10 rpm rotation cycles. The crude protein extract (supernatant) was collected after ultracentrifugation for 20 min at 21,500 rpm and the opsin-FLAG fraction was bound in a 15-mL black tube to 1000 µL Pierce Anti-DYKDDDDK Affinity Resin (50% slurry, Thermo Scientific) at 4°C overnight under gentle rotation. The next day, the resin-bound fraction was loaded on a 5 mL Pierce column, washed 3x by inversion in 3mL cold wash Hepes buffer (3 mM MgCl_2_, 50 mM Hepes pH8.5, 140 mM NaCl) containing 25% Glycerol vol/vol, 0.1% DDM, without protease inhibitors, followed by centrifugation for 1 min at 1,000 x *g* at 10°C to collect wash fractions.

Active opsin complexes in solution were eluted for 20 min at room temperature and 10 rpm rotation, via competitive binding with 150 µL FLAG peptide diluted in wash Hepes buffer (5mg/mL). After centrifugation for 1 min at 1,000 x *g*, the total eluate fraction was concentrated for 1 hour at 4°C and 4,000 rpm using an Amicon Ultra-2 filter column to a final volume of approx. 200 µL, divided in 40-50 µL aliquots in 1.5mL black tubes and stored on ice in the dark. Under dim-red light, 1.5µL-aliquots were loaded on a Nanodrop1000 (Life technologies) to record UV-Vis spectra taken at 5-nm intervals from 200 to 750 nm.

### Bioluminescent plate reader experiments

Luminescence was measured from a single well in a plate reader (Infinite 200Pro, Tecan) with an integration time of 1s every 0.5s for Gq assays, every 60s for Gs and with an integration time of 300ms every 0.5s for Gi assays. Dark luminescent baseline was measured for 5 cycles until the measurement was paused and the plate ejected for illumination.

For Gq assays, one well was subjected to a 1s illumination using a pE-4000 (coolLED Ltd.) and measured. For Gs assays, luminescence was measured for all wells before ejection. 1s light pulse was applied subsequently to each well before the measurement was resumed.

For Gi assays, 2µM forskolin (Sigma-Aldrich) was added to each well to elevate cAMP levels and recording was resumed for 20-30min and illuminated as before. In the 3 assays, luminescent responses were compared to cells transfected only with the corresponding luminescent rapporter, subjected to the same illumination conditions as the opsin.

### GsX and GqX assays

Luminescence was measured for 1s every 60s. Pre-illumination baseline was recorded for 5 cycles before recording was paused and the plate ejected. Light-dependent GsX responses were acquired using light-titration curves. Each well was exposed once to a single light intensity pulse of the same wavelength, starting from the highest intensity and gradually decreasing in 0.5 log10 steps to capture the peak response, especially small signals from low intensities.

As NOL opsins did not signal efficiently through GsX proteins, GqX chimeric proteins were employed instead, following the same principle. Plasmid DNAs encoding Gαq proteins with the last five amino-acids exchanged with the ones from other Gα subfamilies (gift from A. Inoue, Tohoku University, Japan), as well as a native *N. oerstedii* G-protein alpha subunit (GqNO, Genbank Acc. nr. AUC64085.1) were co-transfected individually with a given rhodopsin, and transfected cells were resuspended in 96-well plates as described previously (*39*). Light-dependent GqX responses were acquired using light-titration curves. Similarly to the Gq assay, each well was exposed once to a 1s light pulse of a fixed wavelength and measured at a time with different light intensities, starting from the lowest and increasing in 0.5 log10 steps to avoid cross-light contamination of the neighboring wells.

### Action spectra

Action spectra were measured using the Aequorin Gq-assay. Light stimulations were delivered using a Polychrome V (TILL Photonics) in combination with neutral density filters to keep an equal photon flux across wavelengths. A 3D-printed holder was built and fixed at the end of the light guide to hold ND filters of varying transmittance (Newport) and maintain a fixed distance between the light source and the sample. A controllable motorized slit was incorporated into the holder to allow precise timing of stimulation and to prevent unwanted stimulation at the resting wavelength of the Polychrome.

To measure activation spectra, a 1s light pulse of a fixed wavelength was delivered to one well from 400 nm to 690 nm in 10 nm steps. Each well was illuminated and measured once before changing the wavelength and continuing to the next well. The activation spectra was reconstituted using the maximal relative increase in luminescence for each wavelength, normalised between the minimal and maximal peak response. Data points were fitted with a non-linear least squares optimization of the Govardovskii visual template (*68*) in python 3.12 using SciPy 1.14 (*69*) and scikit-learn (*70*) with initial parameter guesses based on the experimental peak wavelength. For NOM5, NOM6, NOL14 and NOL13, dark action spectra were fitted with a Gaussian Mixture Model (GMM) with two components. The λ_max_ and parameters values, along with their uncertainties, were determined from the peak of the fitted curves using 1000 bootstrap resampling iterations, and the *R*² score was calculated to assess the quality of the fit.

To measure deactivation spectra, wavelength bandwidths were set to 15 nm to ensure sufficient deactivation. A first 1s nearly saturating light pulse was delivered to activate the opsin at a fixed wavelength close to the λmax, followed by a consecutive 1s light pulse of 1 x 10^16^ photons.cm^-^_2_.s^-1^ varying wavelength. The relative deactivation was calculated from the dual stimulation peak response expressed as a percentage of the single stimulation peak response and fitted similarly to the activation spectra.

It is important to note that the decay kinetics of the aequorin luminescent signal in the Gq-coupled assay does not directly correspond to opsin deactivation because the calcium levels are quickly downregulated by cellular mechanisms, independently of the sustained activation of the opsin. Additionally, the intracellular calcium response triggered by the first light pulse rises and falls too rapidly for the second pulse to precisely probe the opsin’s active state. This means that the reduction in luminescence response observed after the second pulse may reflect an interruption of the ongoing calcium signaling initiated by the first pulse, compounded by the regulation of calcium levels.

We also note that peak sensitivities extracted from action spectra did not strictly coincide with maxima measured from dark absorbances, which may be attributable to the surrounding lipid environment in the cell membrane, as *in vivo* opsin properties such as stability, function and absorption may differ slightly in detergent-based buffer systems (*71*).

### GIRK whole-cell patch-clamp experiments

HEK293 cells stably expressing the GIRK1/2 channel (HEK293-GIRK1/2, gift from S. Herlitze, RUB) were used to measure NOM5 and NOM6 induced photocurrents. Cells were cultured in DMEM supplemented with Geneticin (G418, Brand) and seeded into 12 wells-plates at a density of 350 000 cells per well. Cells were transiently transfected individually with pIRES-CMV-Lucyrho-NoM5-TS-IRES-Turbo635 and pIRES-CMV-Lucyrho-NoM6-TS-IRES-Turbo635 (construct B1, fig. S3) using FuGENE HD (Promega). Cells were incubated in the dark for 3-7h and resuspended using Trypsin-EDTA. They were then transferred to 15mm coverslips coated with poly-D-lysine (Sigma-Aldrich) in 35 mm Petri dishes (TPP) and supplemented with 2µM 11*-cis-*retinal under red dim light. 24h after transfection, cells were recorded at room temperature in whole-cell patch-clamp experiments using patch pipettes of resistance between 2 MΩ and 5 MΩ filled with intracellular solution (140mM KCl, 3mM MgCl_2_, 0.2 mM Na_2_GTP, 3 mM Na_2_ATP, 5mM EGTA, 10mM HEPES, pH 7.4 (KOH) and 290 mOsm).

Cells were bathed in high potassium extracellular solution (60mM KCL, 89mM NaCl, 10mM HEPES, 1mM MgCl_2_, 2mM CaCl_2_, pH 7.4 and 310 mOsm) and GIRK photocurrents were recorded in whole-cell voltage-clamp mode at a holding potential of -75mV. Light stimulation was delivered using a pE-4000 with properties specified in the figure legends, guided through a 60X water-immersed objective (Olympus). An Axopatch 200B amplifier connected to a DigiData 1440A digitizer was used to control and acquire electrophysiological recordings via Clampex 10.4 (all from Molecular Devices). Data was sampled at 5kHz, low-pass filtered using a Bessel filter (60Hz cutoff frequency) and analysed in Clampfit 11.4.

### Confocal Imaging

HEK293T cells were resuspended under red dim light to polymer-bottom dishes (ibidi, Gräfelfing) 3-7h after initial transfection, and incubated in the dark at 37°C. At least 12 after resuspension, cells were washed twice with imaging buffer (25mM HEPES and 1.5g/l of L-glucose in 50 ml of 10X Hanks’ Balanced Salt Solution (HBSS 10X, ThermoFisher)) and maintained at 37°C before acquisition.

Samples were imaged using a confocal laser scanning microscope (FV1000, Olympus, Shinjuku, Tokyo, Japan) with a x60 water immersion objective (UPlanSApo, Olympus). A diode laser set to 559 nm was used to excite mScarlet. The acquired images were analysed using Fiji (*72*).

### Data processing and statistics

All statistical analyses were performed with Graphpad Prism V10.5.0 software for Windows. Statistical significance for maximal Gq, Gi and Gs responses were calculated using unpaired one-tailed Student’s t tests for normally-distributed data, and Mann-Whitney U for non-normally distributed data. Hypotheses of normality were tested by the Shapiro-Wilk normality test and homogeneity of variances were tested with the F-test for t-tests. A Welch-correction was applied when the assumption of homogeneity of variances was rejected. In the figures the different levels of significance are indicated by * for p < 0.05, ** for p < 0.01, *** if p < 0.001, and **** if p < 0.0001. Average values are indicated with ± standard error of the mean. Light titration curves in the Gq, GsX and GqX assays were fitted using a dose–response model with a three-parameter sigmoid function, as described previously (*37*).

### Computational modeling of NOM6 and NOL14 conformational ensembles

We aimed to elucidate the inverse spectral properties of NOM6 and NOL14 by analyzing the structural differences between their inactive and active conformational states, corresponding to the 11-*cis* and all-*trans* retinal Schiff base (RSB) bound forms, respectively. Because experimental structures were unavailable, we employed the SPEACH_AF approach (*46*), which generates multiple conformational states through systematic alanine scanning of the multiple sequence alignment (MSA), followed by structural modeling with ColabFold (*73*) using the modified MSA and subsequent model filtering using MolProbity (*74*). In this study, we replaced ColabFold with AlphaFold3 (*45*), which allows prediction of protein structures in complex with covalently bound ligands. As AlphaFold3 employs MSAs generated by JackHMMER 3.4 (*75*), we used these alignments instead of those from MMseqs2 (*76*), given that both share the same A3M format. To focus on evolutionally conserved positions rather than insertions specific to individual sequences, only conserved residues (uppercase letters in A3M) were retained. For this purpose, the SPEACH_AF notebook code was modified from *if temp[n] != ‘-’:* to *if temp[n] != ‘-’ and temp[n].isupper():*.

To increase the number of conformational samples, we applied a sliding window size of five for alanine substitutions and used a distance cutoff of 4 Å for alanine substitutions of adjacent residues, based on the initial structural model. This process produced 64 and 66 MSAs for NOM6 and NOL14, respectively, and 15 structures per MSA were generated using three random seeds. The resulting conformations were evaluated with MolProbity (*74*), as implemented in Phenix 1.21 (*77*) for structural quality assessment. Mean MolProbity scores were calculated for each set, and those exceeding one standard deviation above the mean were discarded. This procedure yielded 885 and 810 conformations for NOM6 and NOL14, respectively. Because these conformations included alanine mutations introduced during MSA-based sampling, we reverted the mutated residues to the wild-type sequences using MODELLER 10.1 (*78*). To model the 11-*cis* and all-*trans* RSB binding modes more accurately, the predicted retinal moiety was removed and redocked using GOLD software 2024.3 (*79*) with retinal coordinates from PDB entries 2Z73 (*80*) (11-*cis*) or 8A6E (*81*) (all-*trans*). Conformations with negative GOLD fitness scores were excluded, resulting in 843 and 776 conformations for the 11-*cis* retinal bound states of NOM6 and NOL14, respectively, and 823 and 758 for the all-*trans* retinal bound states. To identify models representing the inactive and active states, we selected the 11-*cis* RSB-bound (inactive-like) and the all-*trans* RSB-bound (active-like) conformations based on principal component analysis (PCA) of Cα atoms excluding the flexible N- and C-termini (M1-N57 and M357-A385 in NOM6; M1-D45 and L343-A372 in NOL14) using ProDy 2.5 (*82*), visual inspection of key motif conformations, and GOLD fitness scores (figs. S7-8). These selected models were subsequently subjected to molecular dynamics (MD) simulations to remove nonphysical contacts and equilibrate the rhodopsins within lipid and solvent environments.

### Molecular dynamics and hybrid quantum mechanics / molecular mechanics simulations

Molecular dynamics (MD) simulations of NOM6 and NOL14 were performed using GROMACS 2023 (*83*) with CHARMM36m force field (*84*). To investigate the effect of the protonation states of D234 in NOM6 on spectral shifts, MD simulations were carried out for both the deprotonated and protonated forms of D234 in the 11-*cis* and all-*trans* retinal Schiff bound states. Each rhodopsin model was embedded in a bilayer of 1-palmitoyl-2-oleoyl-sn-glycero-3-phosphocholine (POPC) lipids and solvated with TIP3 water model (*85*) containing 150 mM NaCl, using the CHARMM-GUI webserver (*86*) (Table S3). Force field parameters for the RSB were taken from previously published work (*87–89*). Energy minimization and equilibration were performed with a stepwise release of positional and angular restraints on the RSB, protein, and lipids, as summarized in Table S4. The equilibrated systems were then used for three independent 500-ns production MD simulations at temperature of 303 K. A short-range cutoff of 12 Å was applied for both electrostatic and van der Waals interactions, while long-range electrostatics were treated using the particle mesh Ewald method (*90*). Temperature and pressure were maintained at 303 K and 1 bar, respectively, using velocity rescaling (*91*) for temperature control and either Berendsen (*92*) or Parinello-Rahmann (*93*) coupling for pressure control.

Following the simulations, conformational clustering was performed using backbone atoms of the transmembrane helices (ClusterBB) and residues in the retinal binding pocket within a cutoff of 4.5 Å (ClusterRBP) (Table S5). Clustering was carried out using the GROMOS algorithm (*94*), with root mean square deviation (RMSD) cutoffs of 2 Å and 1.3 Å for ClusterBB and ClusterRBP, respectively. Representative structures from each cluster were selected for subsequent analyses.

To compute electronic properties of RSB in the protein environment, hybrid quantum mechanics / molecular mechanics (QM/MM) simulations were conducted using Amber2021 (*95*) with the second-order density functional tight binding (DFTB2) method (*96*) including dispersion correction (*97*). The RSB was designated as the QM region by severing Cα-Cβ bond and capping it with a hydrogen link atom, while the remaining system was treated with classical force fields. Energy minimization was performed using 2,500 steps of steepest descent followed by up to 2,500 steps of conjugate gradient minimization with positional restraints of 10 kcal/mol/Å^2^ on heavy atoms. This was followed by QM/MM equilibration, consisting of a gradual temperature increase from 0 to 303 K, 50 ps of dynamics at 303 K, and a 1-ns production simulation at 303 K using a Langevin thermostat under constant volume, a 1-fs time step, and a 12 Å cutoff for electrostatic and van der Waals interactions. Spectroscopic properties were computed from the QM/MM trajectories by extracting snapshots every 10 ps (100 snapshots in total) and calculating vertical excitation energies using the second-order algebraic diagrammatic construction [ADC(2)] method (*98*) with the correlation-consistent polarized valence double-zeta (cc-pVDZ) basis set in TURBOMOLE 7.7 and 7.8 (*99*).

### Calculation of spectral sensitivity of individual opsins

Action spectra or absorbance of each expressed rhodopsin was multiplied by the transmittance (T=10^−Abs^) of the relevant filter (for distal tiers) or combined filters (for proximal tiers). Normalization was achieved by dividing all values by the peak spectral sensitivity.

---

**Fig. S1.**
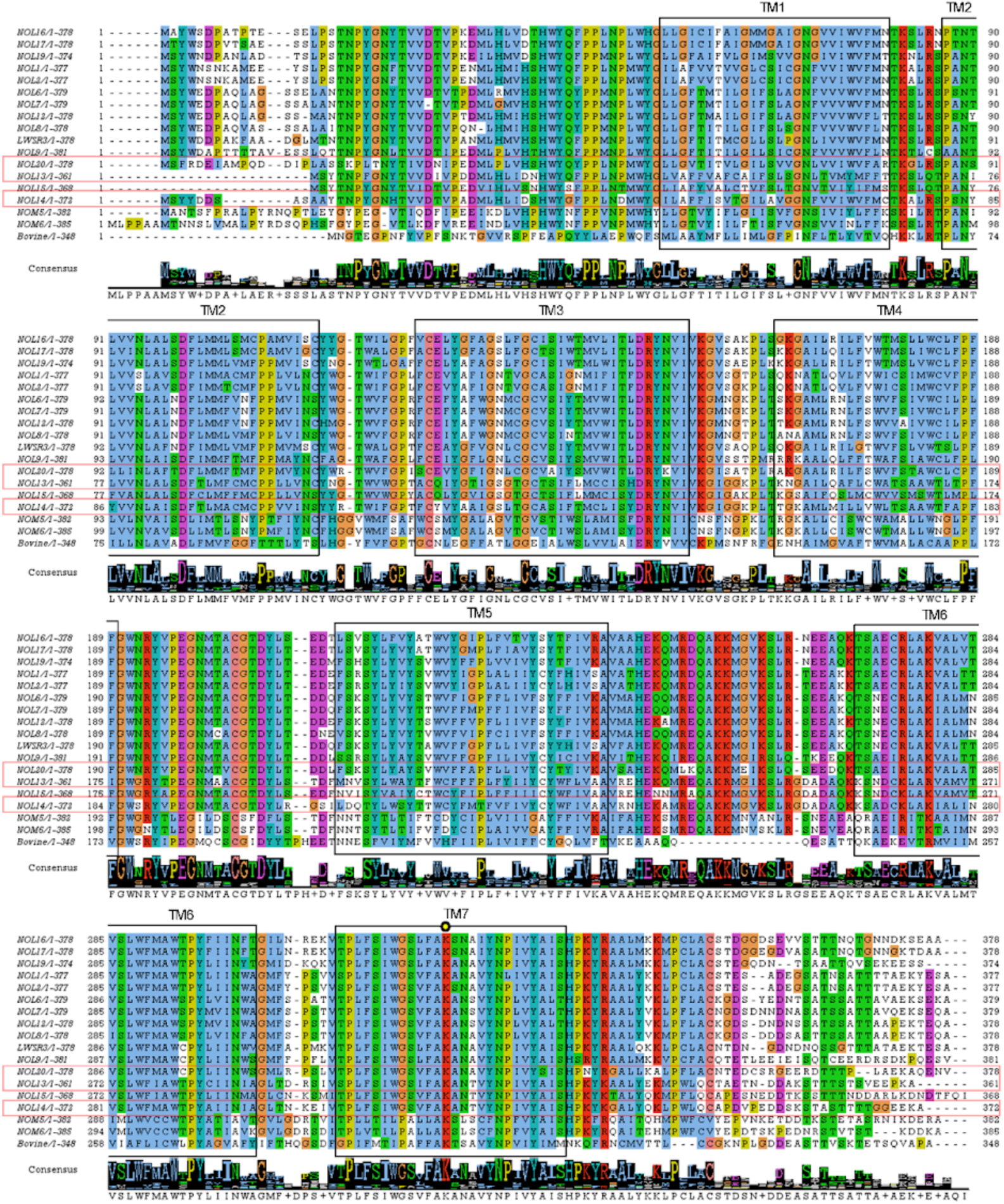
Comparison of the full length NOL and NOM sequences of *N. oerstedii* with bovine rhodopsin as reference. (PDB: 1GZM_A) (*100*). Predicted transmembrane helices are marked with black rectangles based on the bovine rhodopsin model, and the retinal binding lysine is marked with a yellow circle. The most red sensitive rhodopsins are marked with red rectangles.

**Fig. S2.**
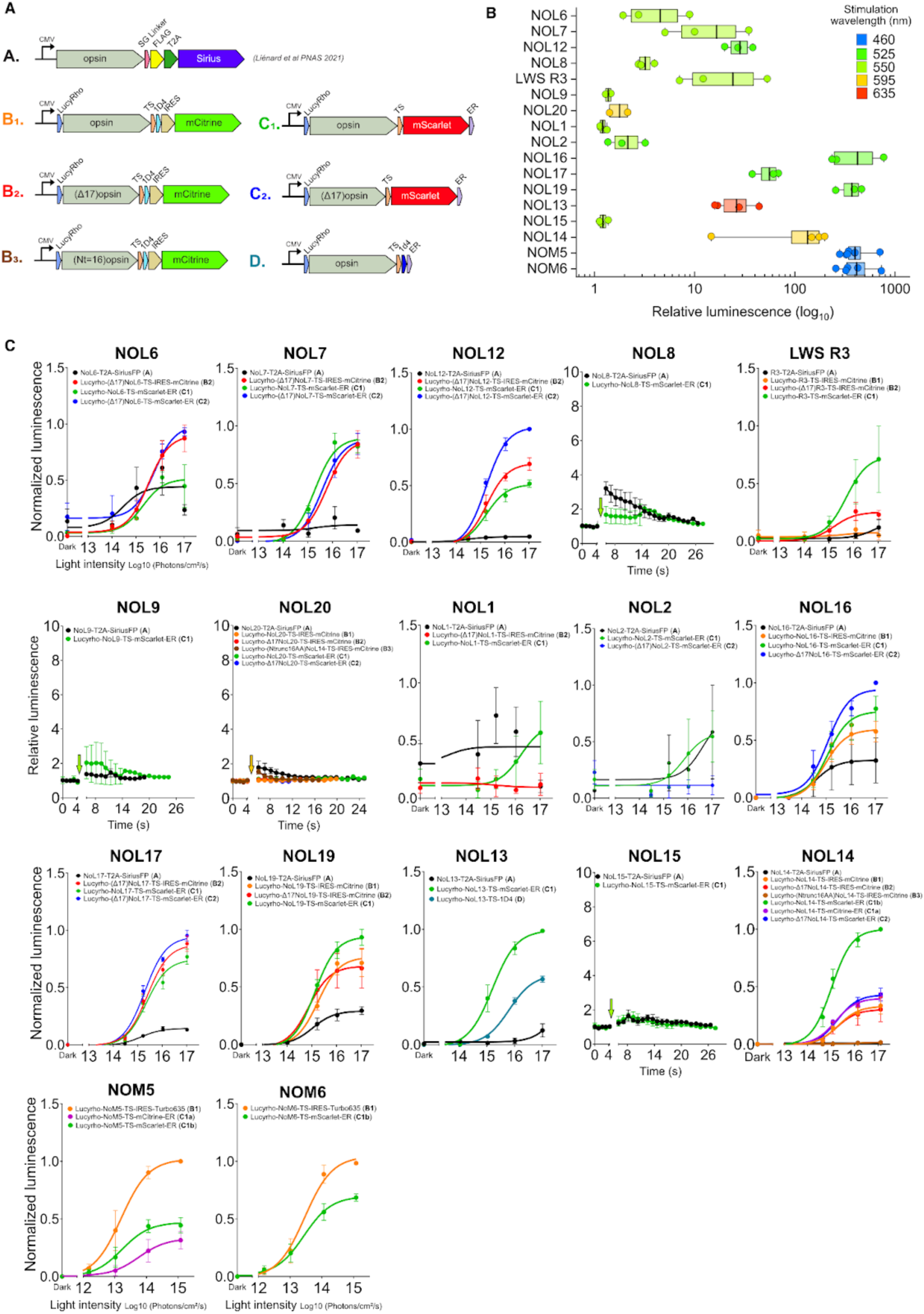
Optimisation of constructs. (**A**) Design of DNA constructs used for optimisation of the luminescent responses (SG linker 2aa, 1D4 affinity tag TETSQVAPA 9aa, FLAG affinity tag DYKDDDDK 8aa, LucyRho self cleavable leucine-rich signal peptide, T2A Self cleaving peptide from *Thosea asigna* virus 2A GSGEGRGSLLTCGDVEENPGP 21aa, TS Golgi export trafficking signal sequence KSRITSEGEYIPLDQIDINV 20aa, ER endoplasmic reticulum export sequence FCYENEV 7aa, IRES internal ribosome entry site, Δ17 deletion of t17aa of an opsin N-terminus, Nt=16 only 16aa remaining upstream of TM1 of the opsin). Construct A. has been previously described (*101*). (**B**) Comparison of relative aequorin luminescent levels of the most effective (highest signal) constructs for each opsin following a 1s activation (wavelength indicated in the legend). (**C**) Light-titration aequorin responses of various DNA constructs used in this study. Unresponsive opsins (NOL8, NOL9, NOL15, NOL20) signals are shown as relative luminescence over time, at the highest light intensity (1x10^17^ photons.cm^-2^.s^-1^).

**Fig. S3.**
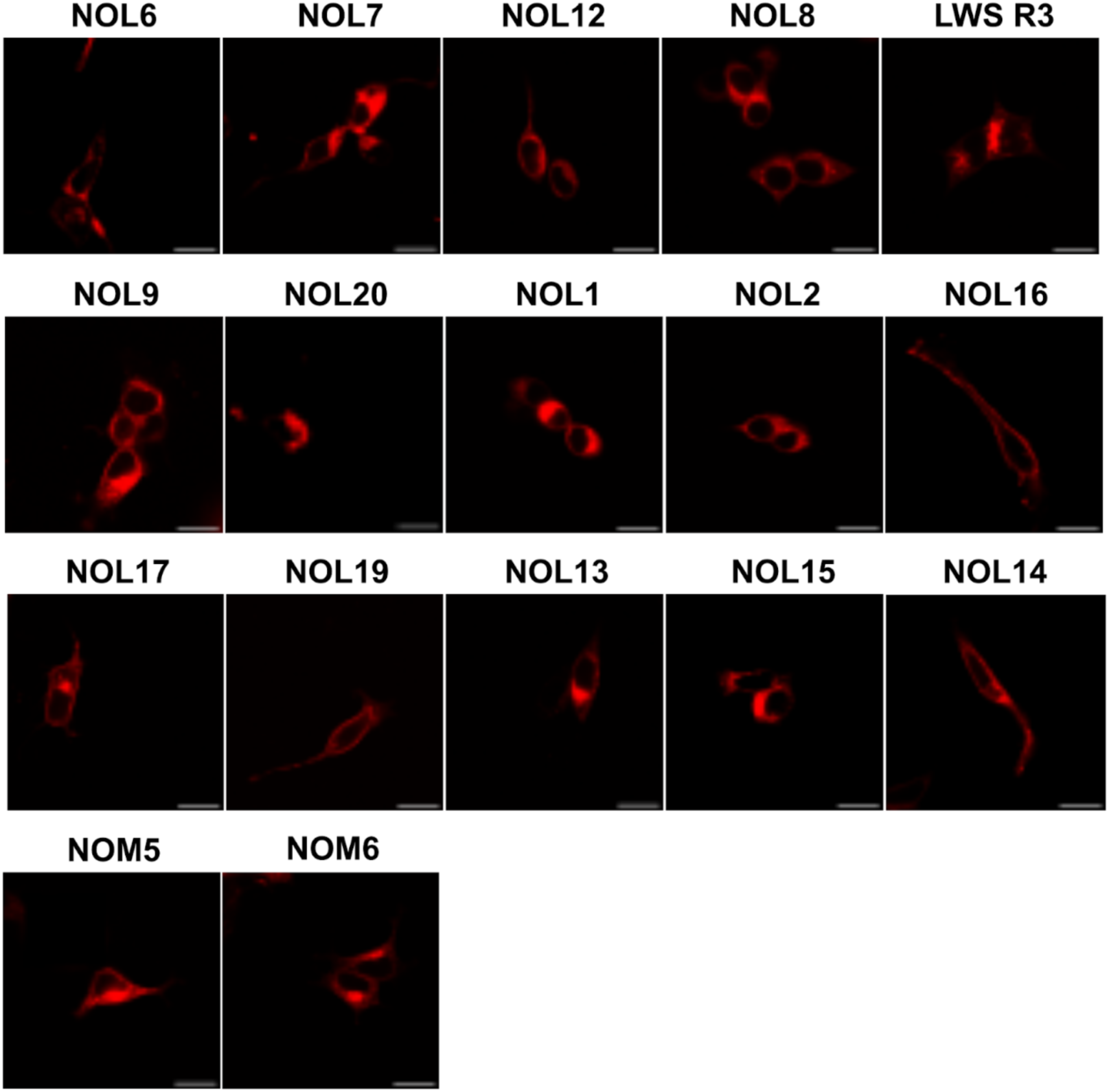
Expression profile of NOM and NOLs opsins. HEK293T cells expressing NOM or NOL opsins fused to mScarlet at the C-termini (construct C1 in fig. S2). Scale bar = 20µm.

**Fig. S4.**
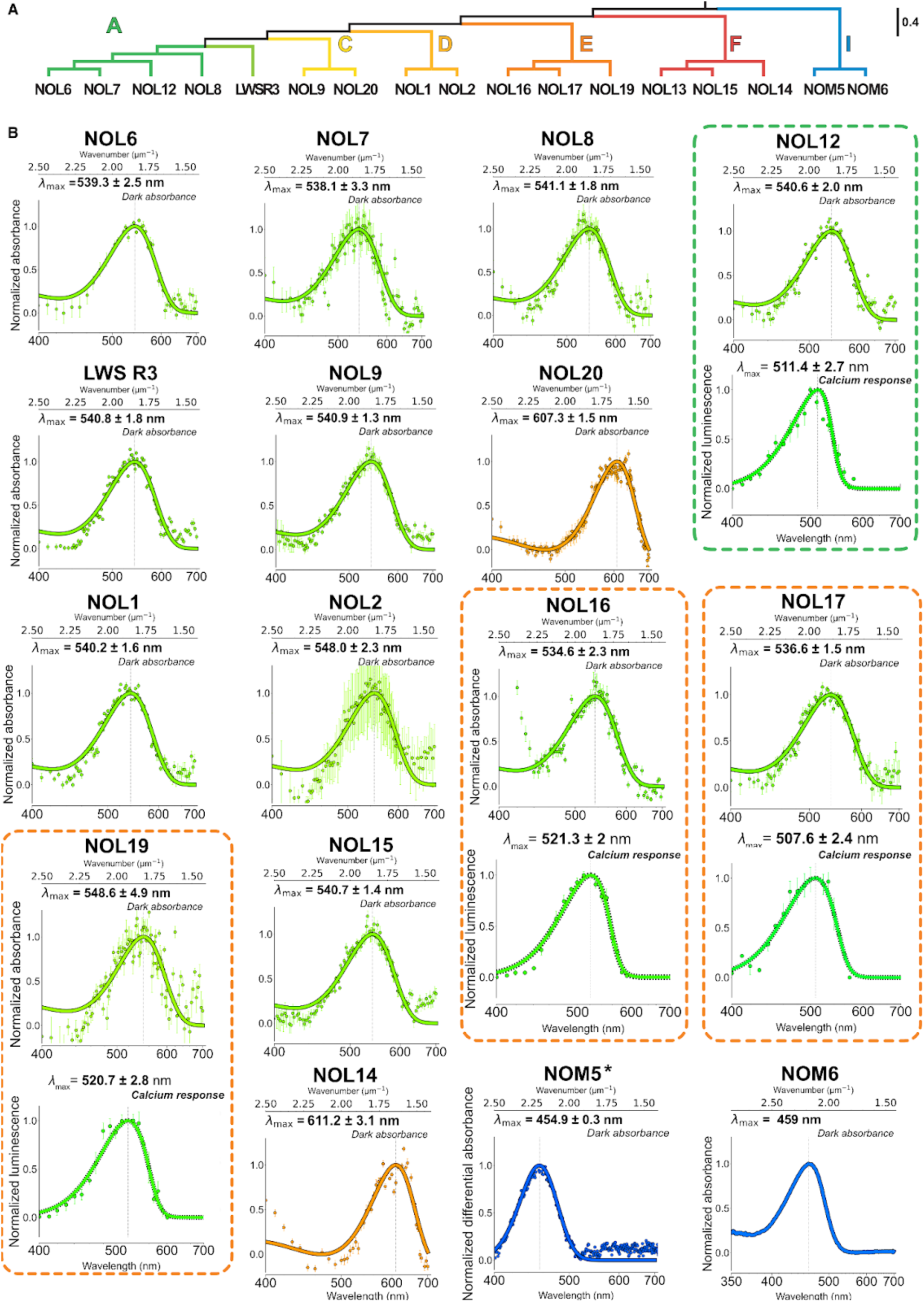
Dark absorption and action spectra of NOL and NOM opsins. (**A**) Phylogenetic relations of NOLs and NOMs from clade A, C, D, E, F and I (adapted from Porter et al. 2020). (**B**) Dark absorbance spectra measured in an UV-vis Nano-spectrometer after reconstitution with *11-cis* retinal, with corresponding action spectra below when possible, established from maximal aequorin responses, as indicated by the dashed frames, color coded according to phylogenetic branch. Light intensity of the 1 s pulse used for the action spectra : NOL12 : 8x10^14^ photons cm^-2^ s^-1^ (n=4); NOL16 (n=3) and NOL17 (n=3) : 4x10^14^ photons cm^-2^ s^-^_1_ ; NOL19 : 8x10^13^ photons cm^-2^ s^-1^ (n=4).Absorbance and action spectra of all NOLs were fitted with a non-linear least squares optimization of the Govardovskii visual template to estimate the best-fit *λ*_max_. Data is presented as mean values and with ± SEM for NOLs. NOL6 and NOL14 spectra were previously reported (*101*). Dark absorption for NOM5 was measured with *9-cis* retinal (*) and plotted as the differential absorbance between the dark absorption and the absorption after a 60s pulse of light (455 nm), fitted with a simple gaussian. For NOM6 a continuous spectra without any fitting but corrected for Mie scattering is shown. Bimodal action spectra for NOL13, NOL14, NOM5 and NOM6 are presented in Fig. 2.

**Fig. S5.**
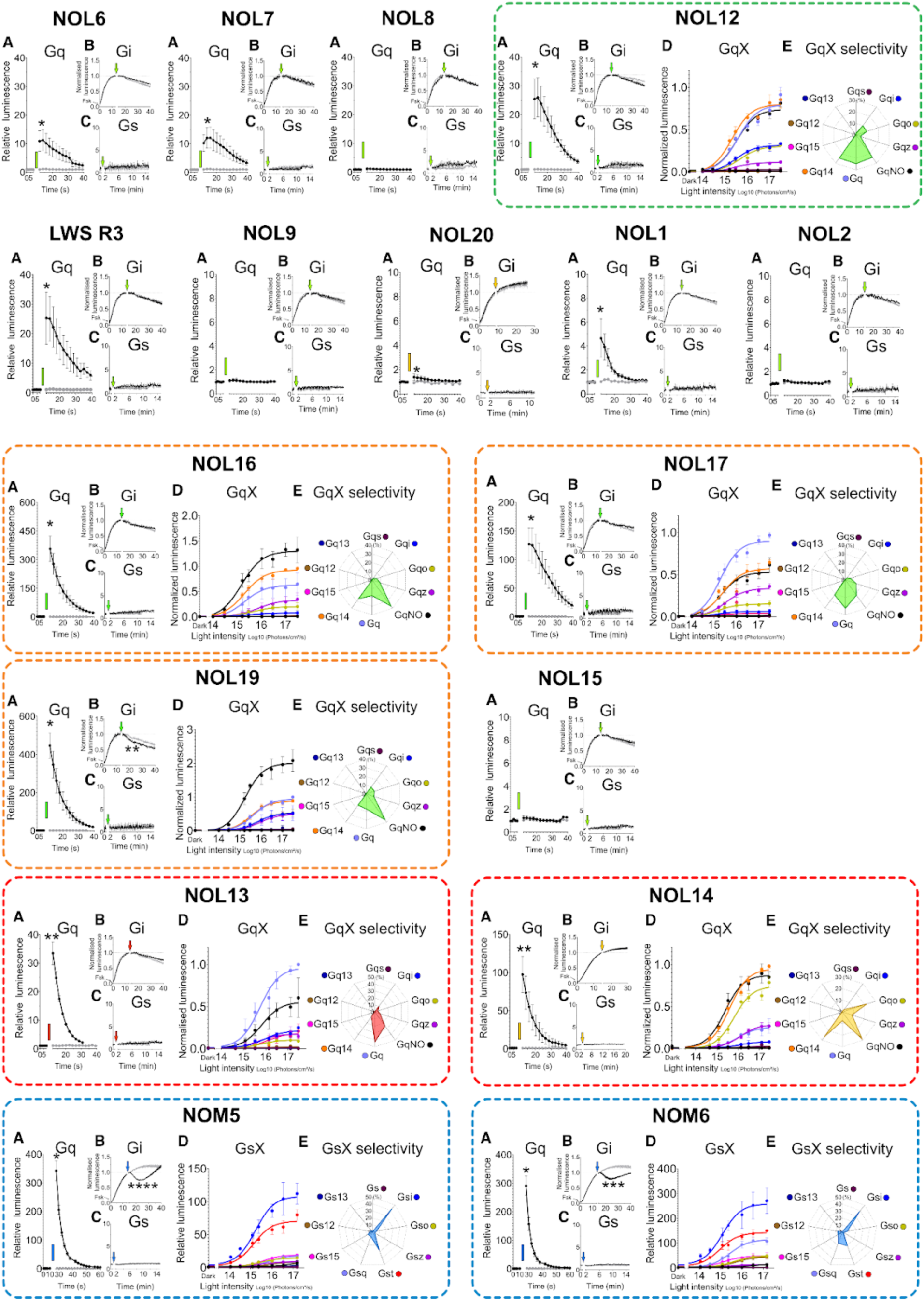
G-protein selectivity profiles of NOL and NOM opsins. (A) Responses to the Gq-protein family were measured by using the aequorin assay (see Table S1 for detailed values). (B-C) Coupling efficiency to the (B) Gi- and (C) Gs-protein families in the glosensor assay. Relative luminescence values are shown as the raw luminescence divided by the averaged pre-illumination baseline. For Gi, cAMP luminescent levels were elevated using 2µM forskolin, 15 min before illumination. (D) Light-dependent G-protein signaling was further investigated for NOMs using the GsX assay and for NOLs using the GqX assay when possible (indicated by dashed frames, colourized based on their phylogenetic clade (see fig. S4). Light-dependent maximal post-stimuli responses are normalized between the no GqX/GsX control and the maximal relative response. (E) Selectivity profiles were established from maximal responses to GsX or GqX chimeras and expressed as percentages of the sum of GsX or GqX protein activity. NOL12, NOL16, NOL17, NOL19 were stimulated with 525 nm light, NOL6, NOL7, NOL8, NOL9, NOL1, NOL2, NOL15, LWS R3 with 550 nm, NOL14, NOL20 with 595 nm, and NOL13 with 635 nm. All responses were measured after a dark pre-illumination baseline followed by a 1s light pulse, indicated by colored arrows and rectangles. All data is presented as mean ± SEM. Different levels of significance at the peak are indicated by * for p < 0.05, ** for p < 0.01, *** for p < 0.001, and **** for p < 0.0001, compared to cells transfected only with luminescent reporters (grey traces).

**Fig. S6.**
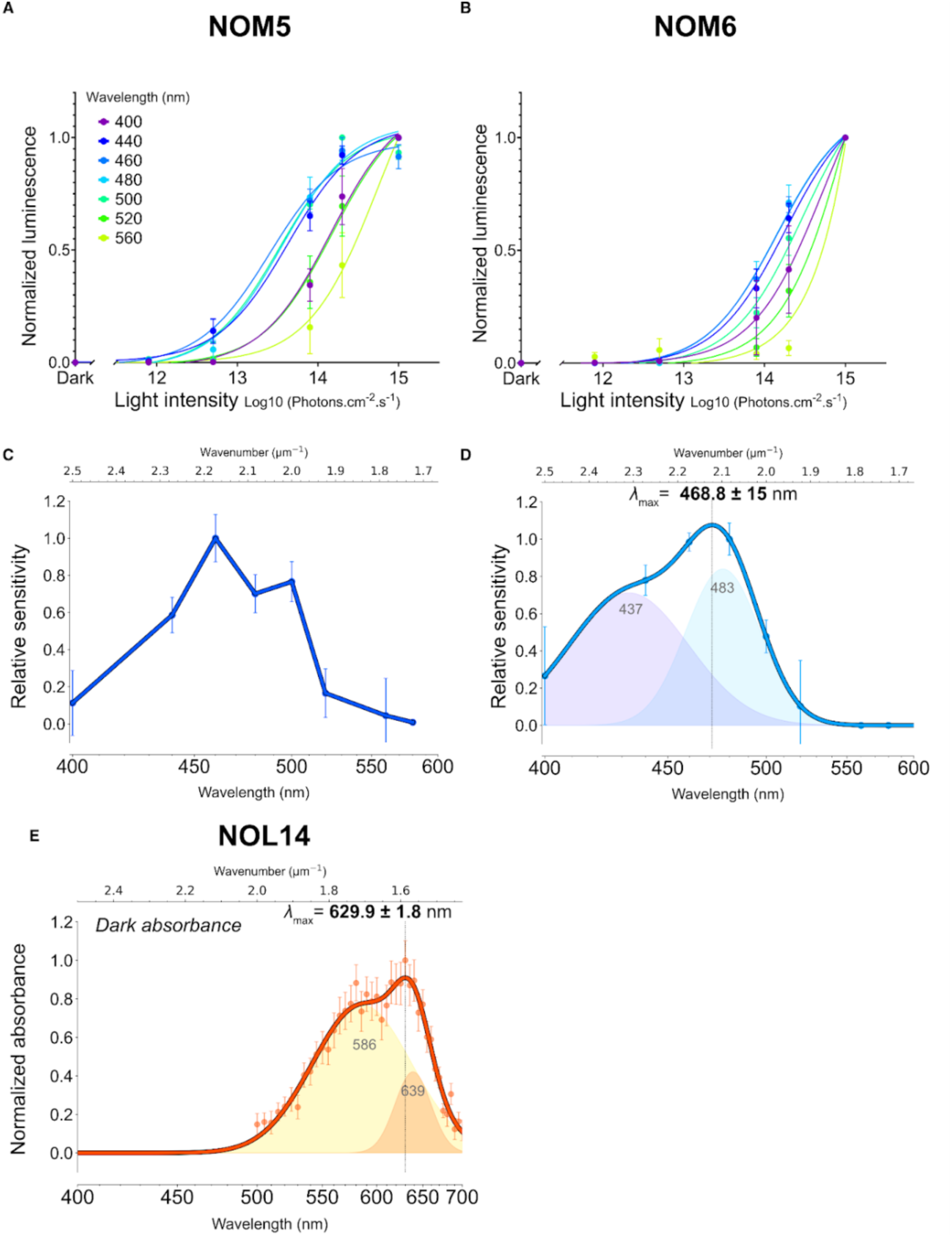
Dark state spectral composition of NOM5, NOM6 and NOL14. Light-dependent aequorin responses following a 1s light pulse of (**A**) NOM5 and (**B**) NOM6 normalized to the maximal luminescent response per wavelength (n=3). Spectral relative sensitivity of (**C**) NOM5 and (**D**) NOM6 calculated from EC50 values extracted from a. and b. at each wavelength. (**E**) Dark absorbance spectrum of NOL14 (data from (*101*)). NOM6 and NOL14 spectra are fitted with a two component Gaussian Mixture Model (GMM) suggesting the existence of two probably pH-dependent isoforms. Data is presented as mean ± SEM.

**Fig. S7.**
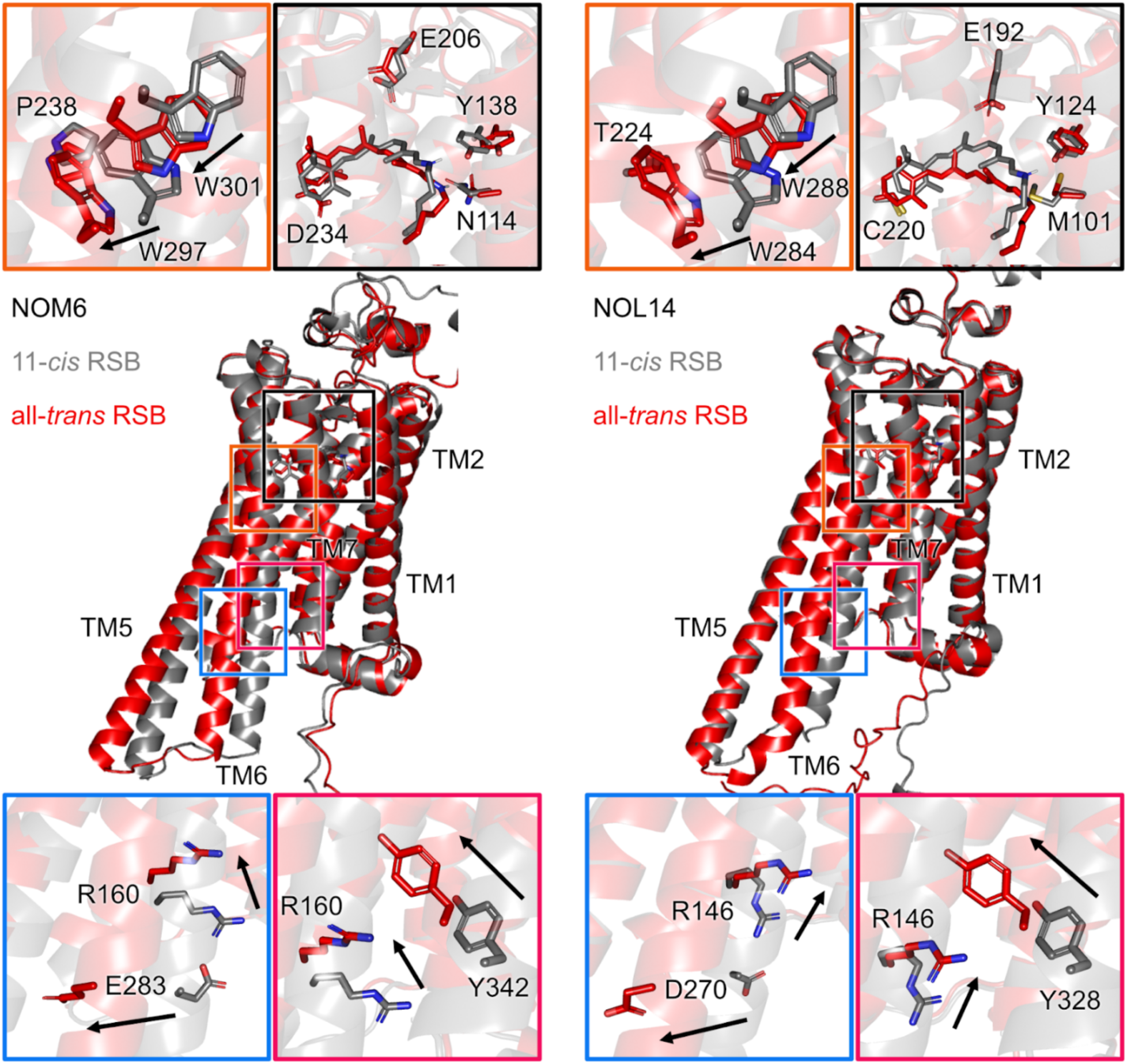
Comparison of computed structural models between NOM6 and NOL14. Inactive (11,*cis*-retinal Schiff base bound) and active (all-*trans* retinal Schiff base bound) states are shown as gray and red cartoons, respectively. In the active states, conserved motifs undergo similar rearrangement that distinguish them from the inactive states (*102*). The PIF motif contains a tryptophan substitution at position 6.44 adjacent to the CWxP tryptophan. Activation rearranges these residues (W297/W301 in NOM6 and W284/W288 in NOL14), reshaping the hydrophobic core, disrupting the DRY ionic lock (R160-E283 and R146-D270), and reorienting the NPxxY tyrosine (Y342 and Y328) toward the intracellular cavity.

**Fig. S8.**
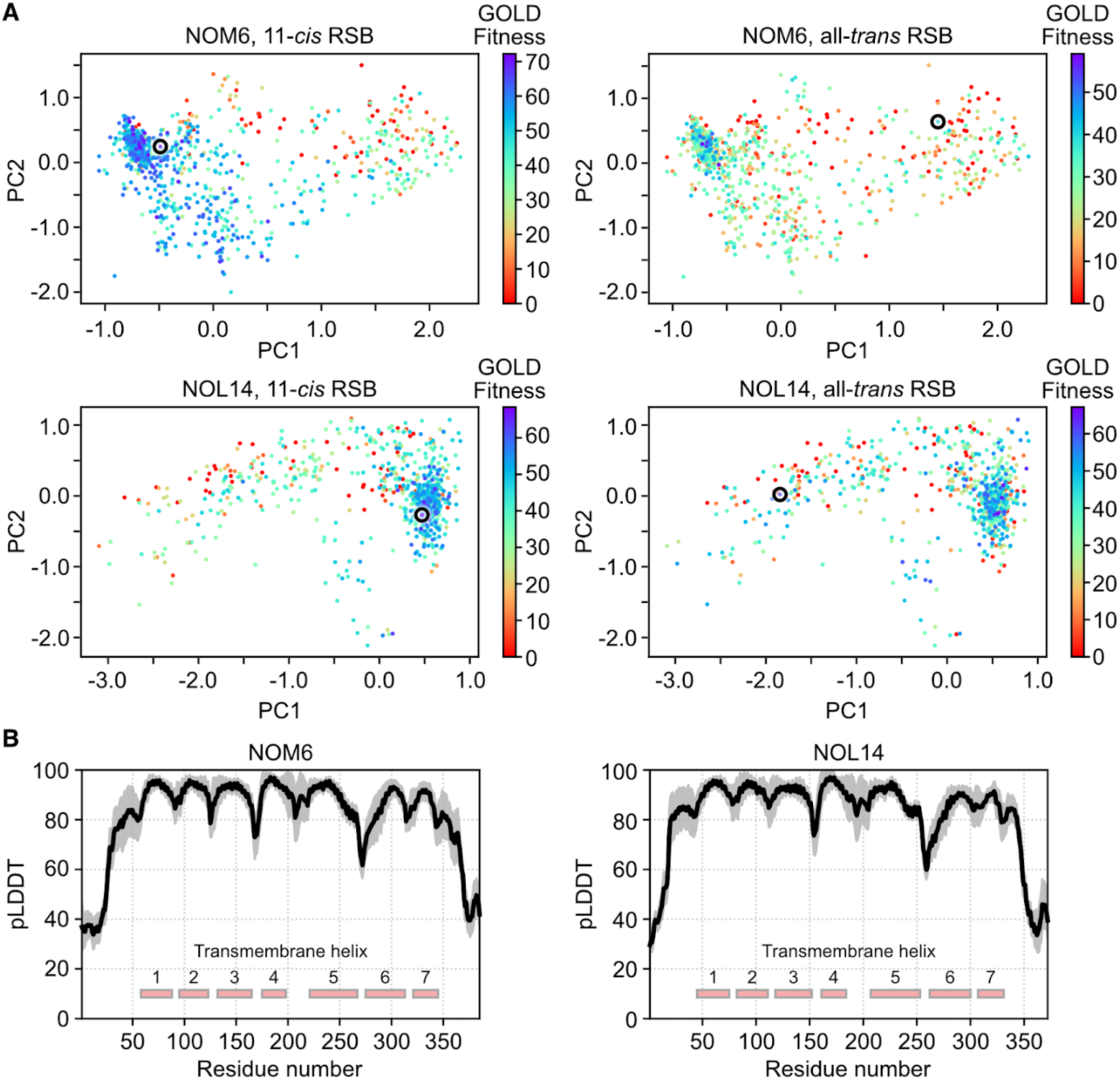
Predicted conformational ensembles and evaluation of model accuracy. (**A**) Principal component analysis (PCA) of NOM6 and NOL14 conformations, excluding the N- and C-terminal regions. Docking scores for the 11-*cis* and all-*trans* retinal Schiff bases are shown by color coding, and the conformations with negative scores were discarded. Selected inactive-like (left) and active-like (right) states are indicated by black circles. (**B**) Predicted local distance difference test (pLDDT) scores of the conformations. Transmembrane helices are shown as red rectangles. The black line represents the mean pLDDT of Cα atoms, and the gray shading indicates the standard deviation. The number of conformations analyzed was 885 for NOM6 and 810 for NOL14.

**Fig. S9.**
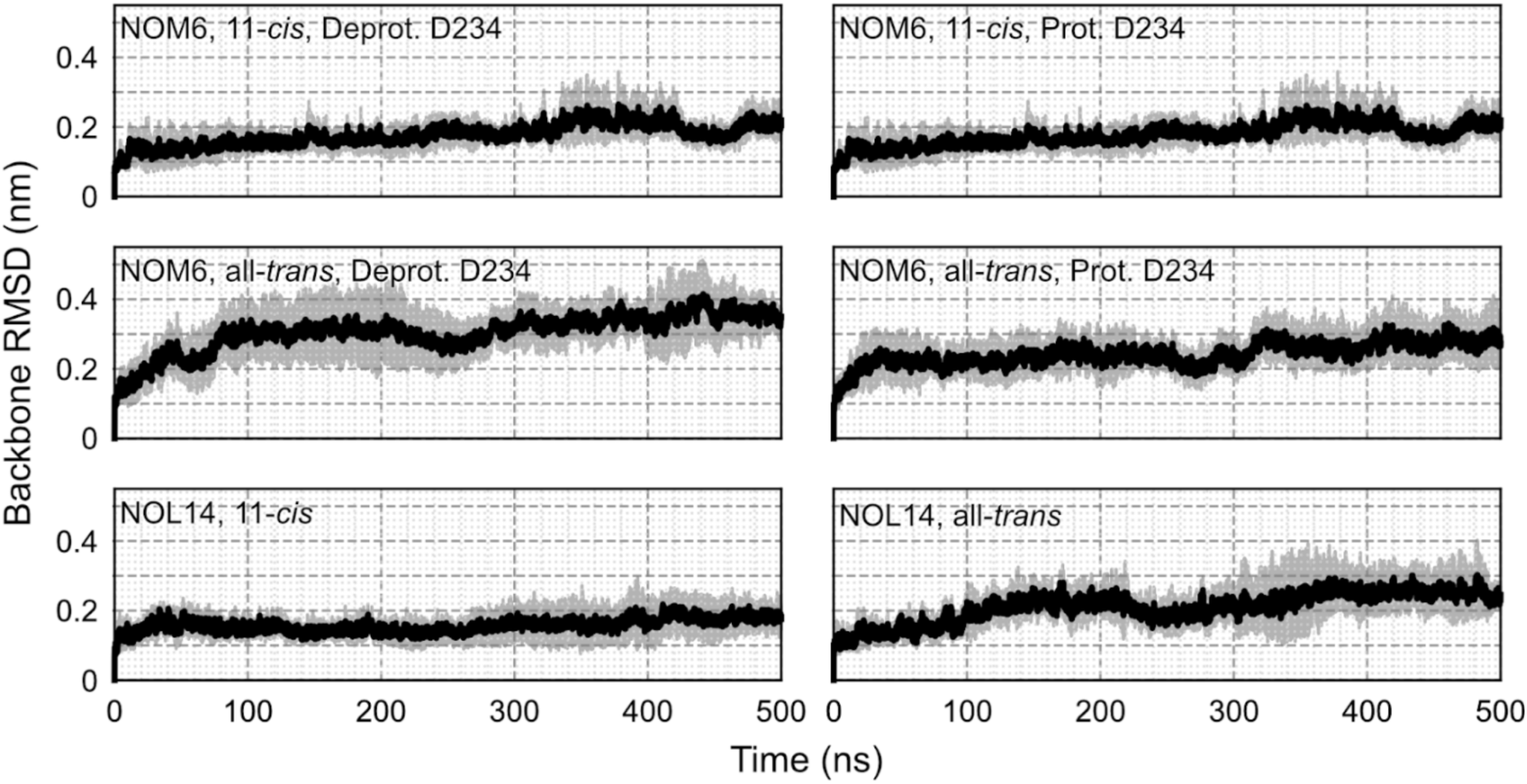
Root mean square deviations (RMSDs) of NOM6 and NOL14. Backbone RMSDs were calculated excluding the N- and C-terminal regions. The solid line represents the mean RMSD, and the shaded area indicates the standard deviation. Molecular dynamics simulations were performed for 500 ns and repeated three times.

**Fig. S10.**
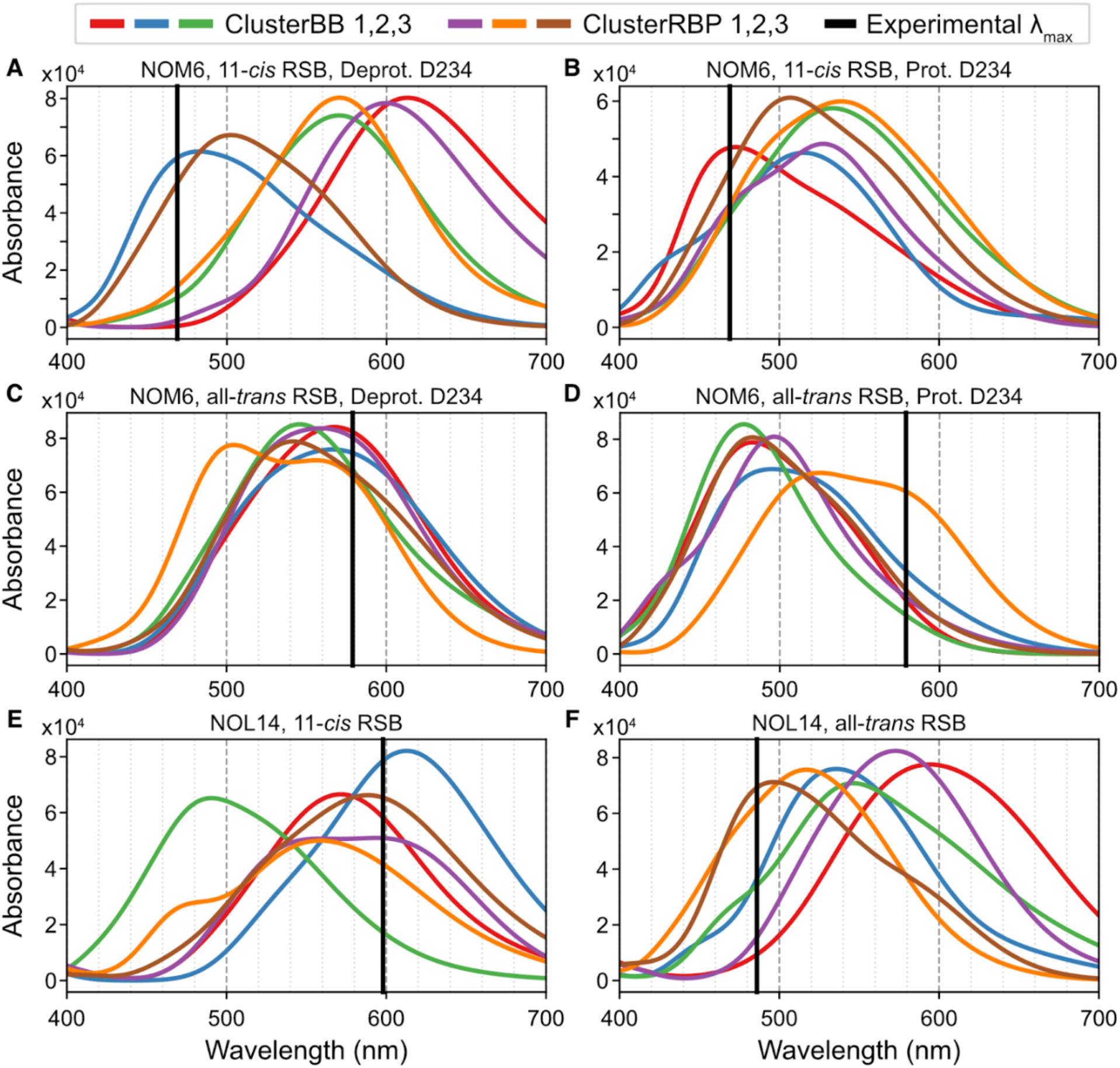
Averaged UV-Vis spectra of NOM6 and NOL14. Spectra calculated at ADC(2) / cc-pVDZ level from snapshots obtained using DFTB2 with dispersion correction. (**A-F**) The experimental mean absorption maximum is shown as a black solid line. Selected clusters are as follows: NOM6, 11-*cis* RSB with protonated D234 (λ_max_ = 473 nm, ClusterBB 1 in (**B**); NOM6, all-*trans* RSB with deprotonated D234 (λ_max_ = 566 nm, ClusterBB 2 in (**C**)); NOL14, 11-*cis* RSB (λ_max_ = 594 nm, ClusterRBP 1 in (**E**)); and NOL14, all-*trans* RSB (λ_max_ = 497 nm, ClusterRBP 3 in (**F**)). Note that NOM6 with 11-*cis* RSB and deprotonated D234 (**A**) and NOM6 with all-*trans* RSB and protonated D234 (**D**) were not selected, as their calculated spectral trends were inconsistent with the experimental observations.

**Fig. S11.**
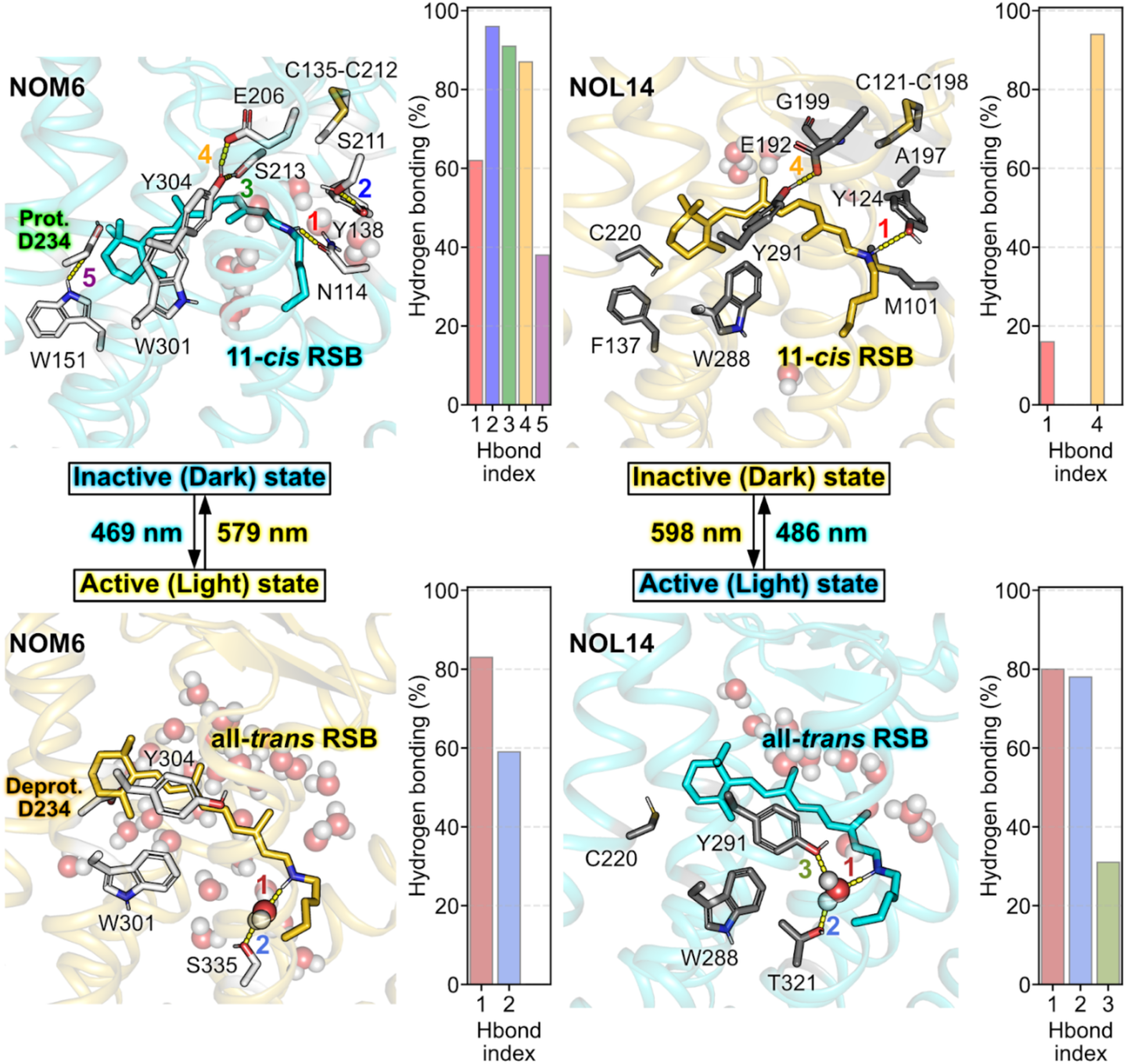
Hydrogen bonding analysis of NOM6 and NOL14. Percentage of hydrogen-bonding interactions were analyzed over 1-ns hybrid quantum mechanics / molecular mechanics (QM/MM) simulations using a distance cutoff of 3.5 Å between donor (D) and acceptor (A) atoms and an angle cutoff of 150° for the D-H-A geometry. Water molecules are shown as spheres.

**Fig. S12.**
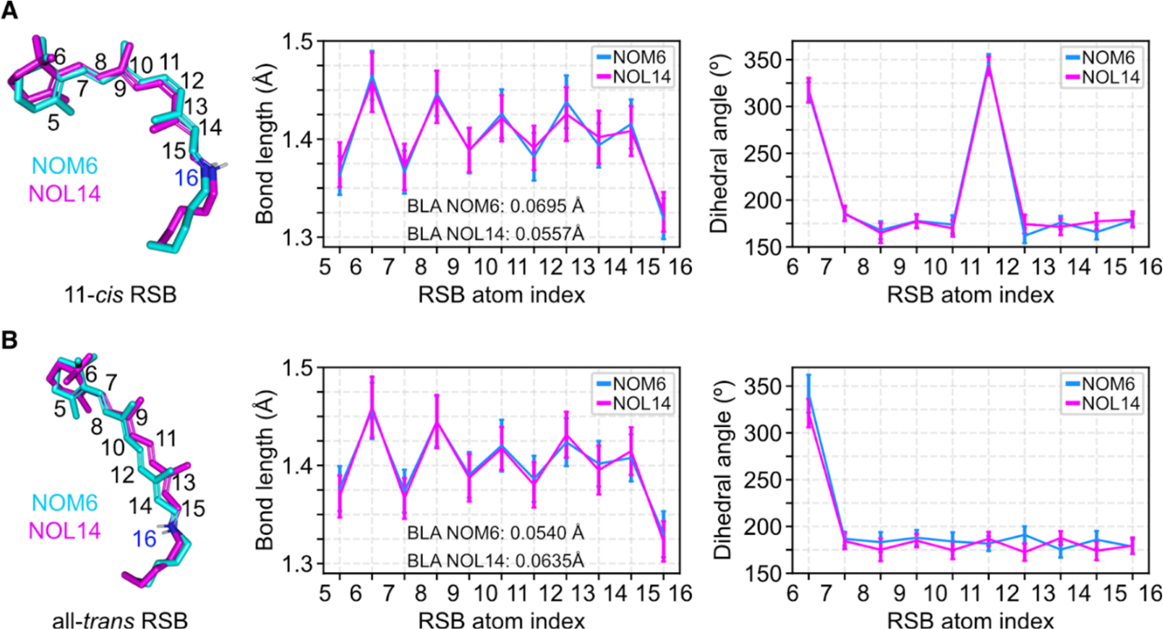
Bond length and dihedral angle analysis. Bond lengths and dihedral angles of (**A**) 11-*cis* and (**B**) all-*trans* retinal Schiff base bound in NOM6 (cyan) and NOL14 (magenta) were obtained from 1-ns hybrid quantum mechanics / molecular mechanics (QM/MM) simulations. The bond length alternation (BLA) was calculated as the difference between the average lengths of single and double bonds.

**Fig. S13.**
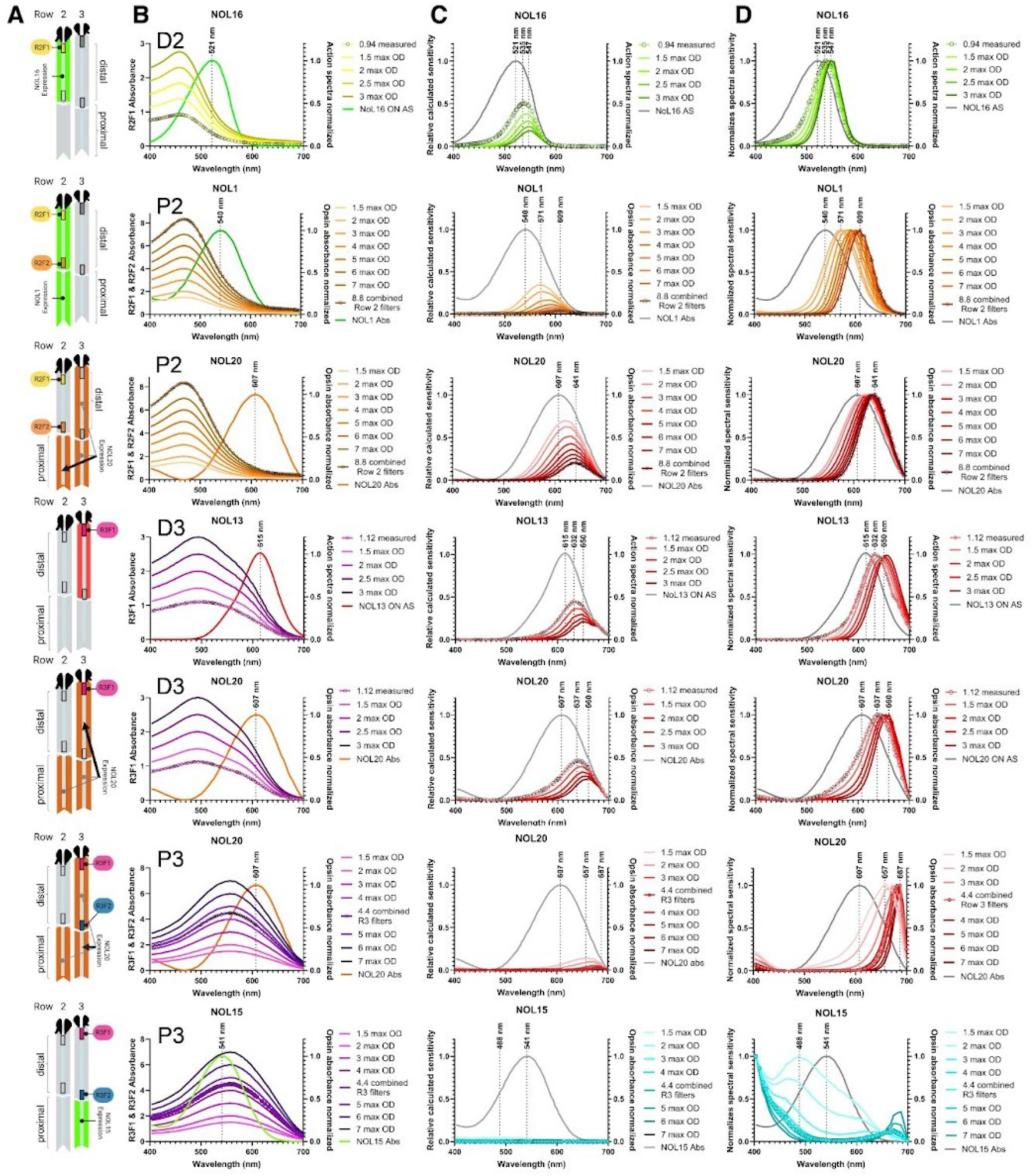
Effect of intra-rhabdomal filter properties on the rhodopsin spectral sensitivities in tiers D2, P2, D3 and P3. (**A**) Schematic representation of midband rows 2 and 3 including the rhodopsins and the filters relevant for the analysis presented in each row. (**B**) Normalized rhodopsin spectra and filter spectra of different optical density ranging from 1.5 to 3 for distal (D) filters, and ranging from 1.5 to 7 for combined filters in the proximal (P) tier. Digitized filter absorbances reported in the literature (*5*) are labelled by black circle symbols. (**C**) Calculated sensitivities of the indicated rhodopsins unfiltered (grey) and after filtering by screening pigments with maximal absorbance ranging from 1.5 to 3 for distal F1 filters, and ranging from 1.5 to 7 for proximal F2 filters. Numbers above vertical dashed/dotted lines represent, from left to right, the wavelength maxima before screening, after screening by filters of reported optical densities (OD), and after maximal screening when applicable (NOL15 is almost entirely shielded by intra-rhabdomal pigments and the low resulting sensitivity could not be fitted). (**D**) Normalized calculated sensitivity spectra plotted with original spectral sensitivity (grey). In all graphs, absorbances and sensitivities resulting from originally measured filters (*5*) are labeled by black open circle symbols.

**Table S1.**
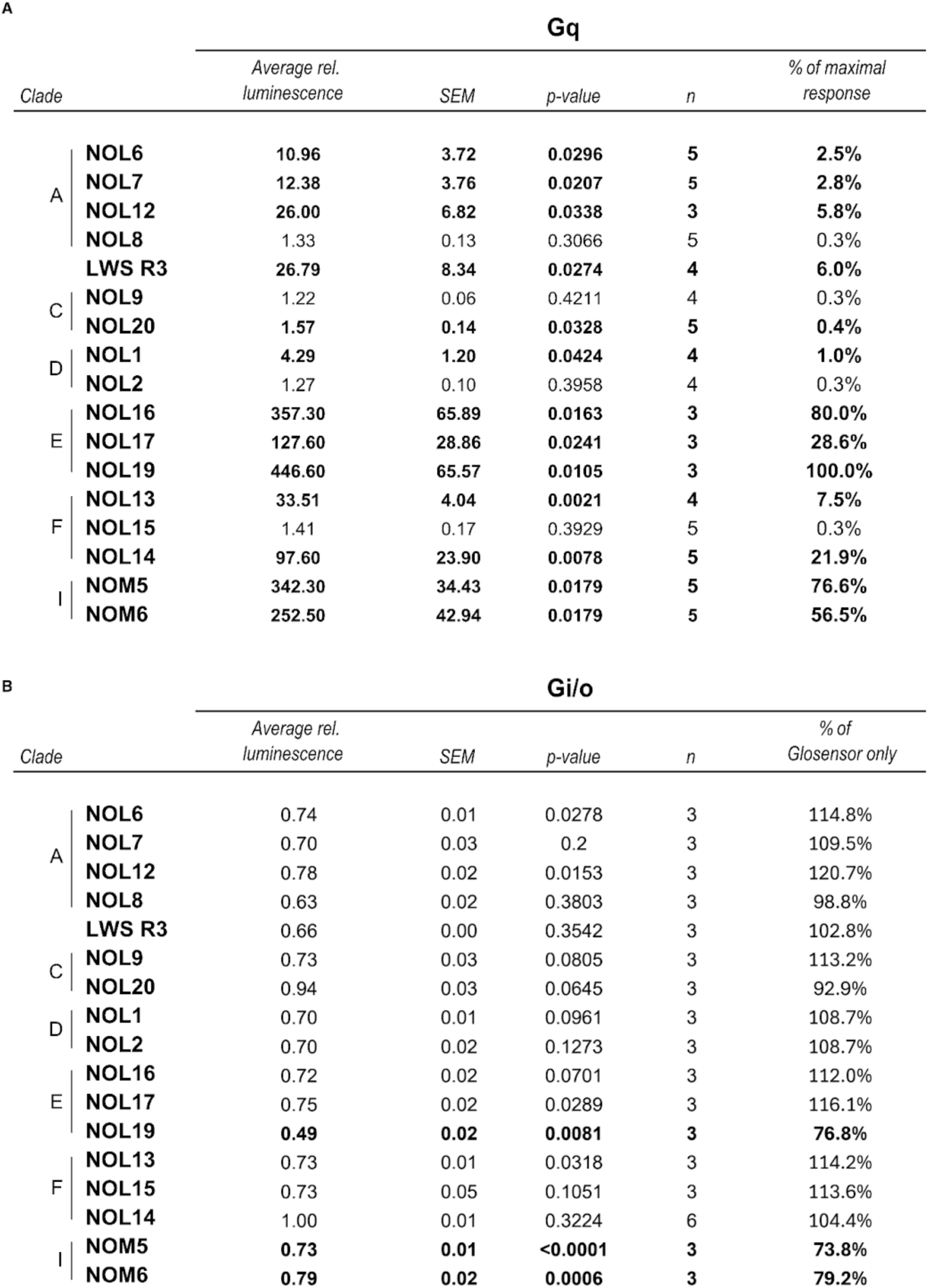
Gq- and Gi/o-coupled response characteristics measured in the aequorin and glosensor assay. Significant Gq and Gi/o-responses are reported in bold. (**A**). Maximal relative Gq-responses are expressed as a percentage of the highest Gq-signaling opsin (NOL19). (**B**). Maximal relative Gi/o responses are reported for each opsin, as the percentage of response of cells transfected only with glosensor (Glosensor only), at the peak decrease in luminescence, between the forskolin-induced plateau and the estimated response decay (see Methods). Table S2.

**Table S2.**
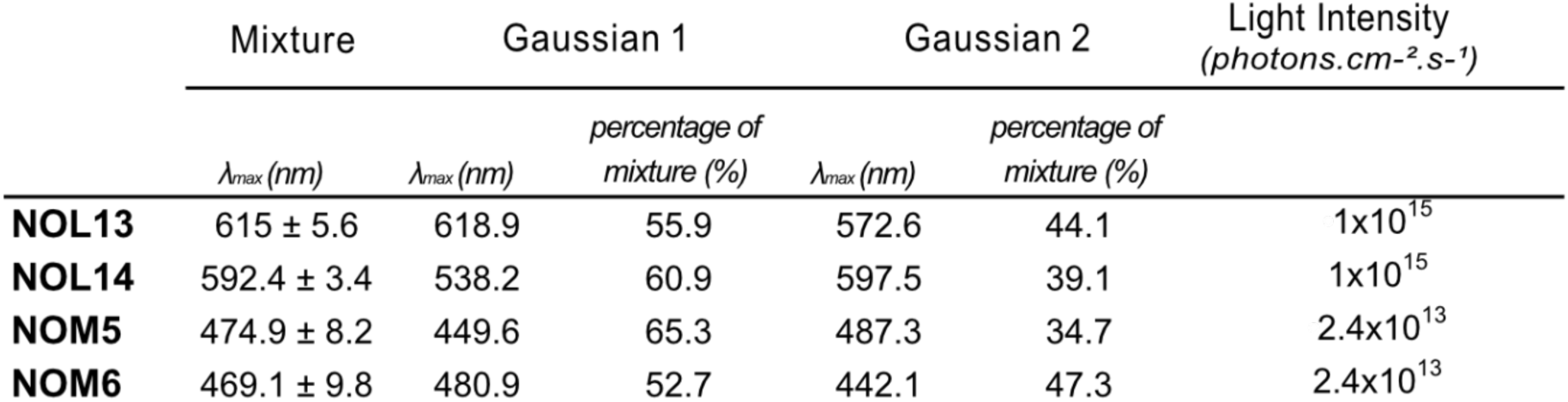
Dark state spectral characteristics of photoreversible opsins. The percentage of mixture for the individual gaussian is reported as the percentage of the area under the curve of the combined GMM model.

**Table S3.**
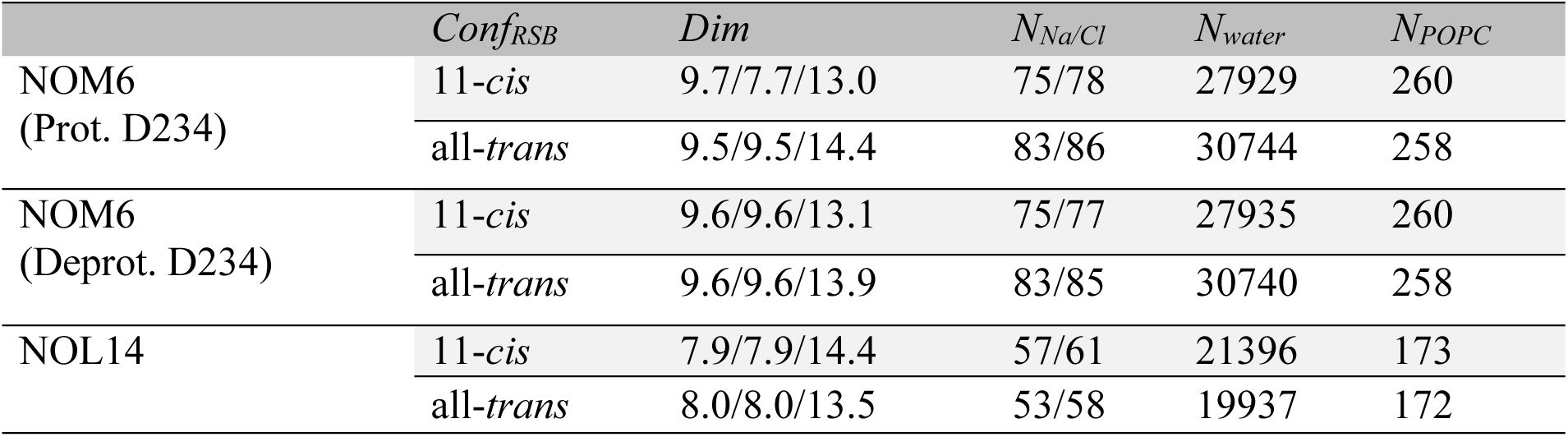
System setup for molecular dynamics simulations. *Conf_RSB_*: configuration of retinal Schiff base, *Dim*: box size (X/Y/Z nm), *N_Na/Cl_*: the number of Na^+^/Cl^-^, *N_water_*: the number of water molecules, *N_POPC_*: the number of POPC. All simulations were conducted at a NaCl concentration of 150 mM and at a temperature of 303 K and replicated three times.

**Table S4.**
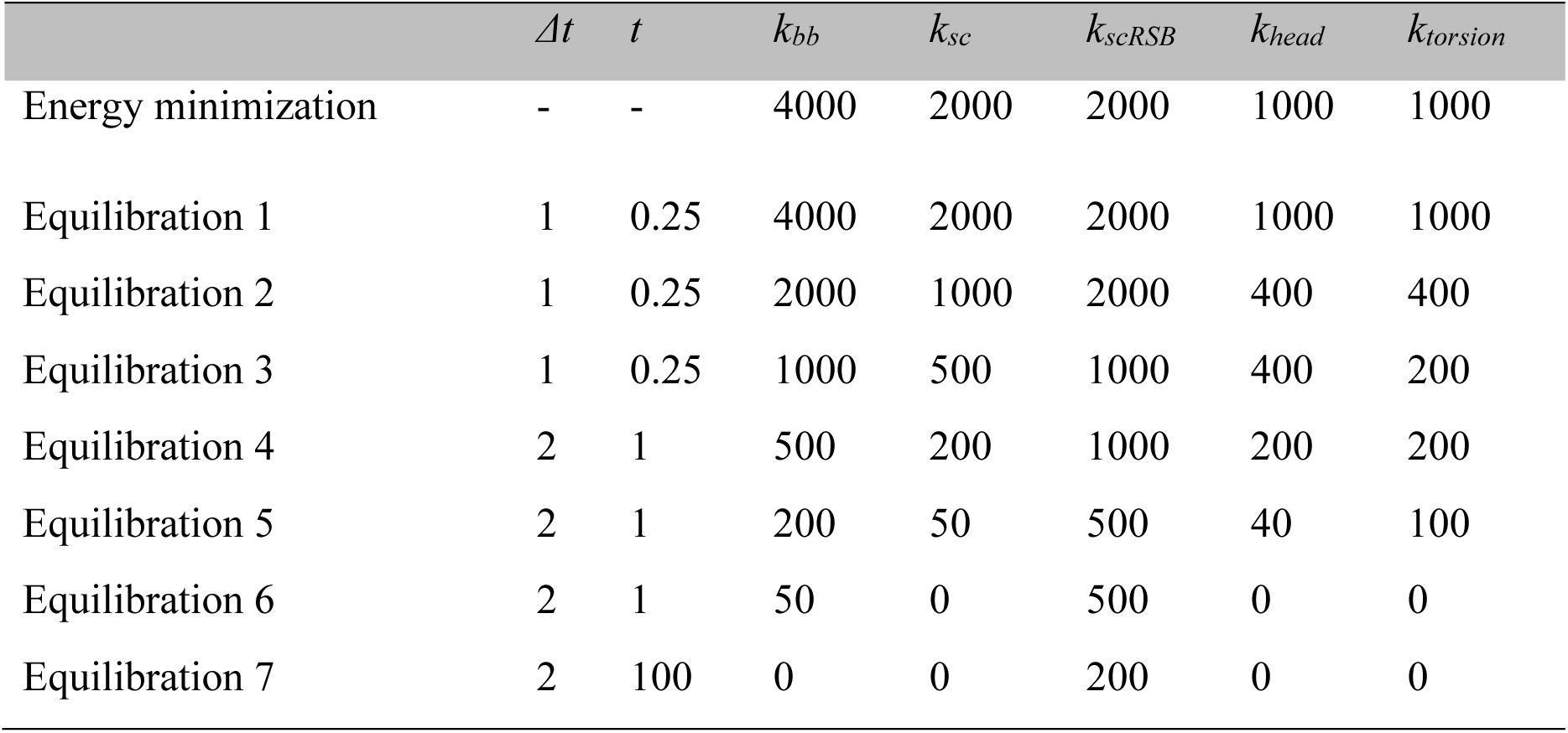
Step-wise release of restraint of applied positional and dihedral angular restraints for energy minimization and equilibration simulations. Time step: *Δt* (fs), individual simulation time: *t* (ns), force constant (kJ/mol/nm) for position/angle harmonic restraints of backbone heavy atoms (*k_bb_*), side-chain heavy atoms (*k_sc_*), side-chain heavy atoms of retinal Schiff base (*k_scRSB_*), lipid head group (*k_head_*), and lipid chirality and *cis* double bond (*k_torsion_*).

**Table S5.**
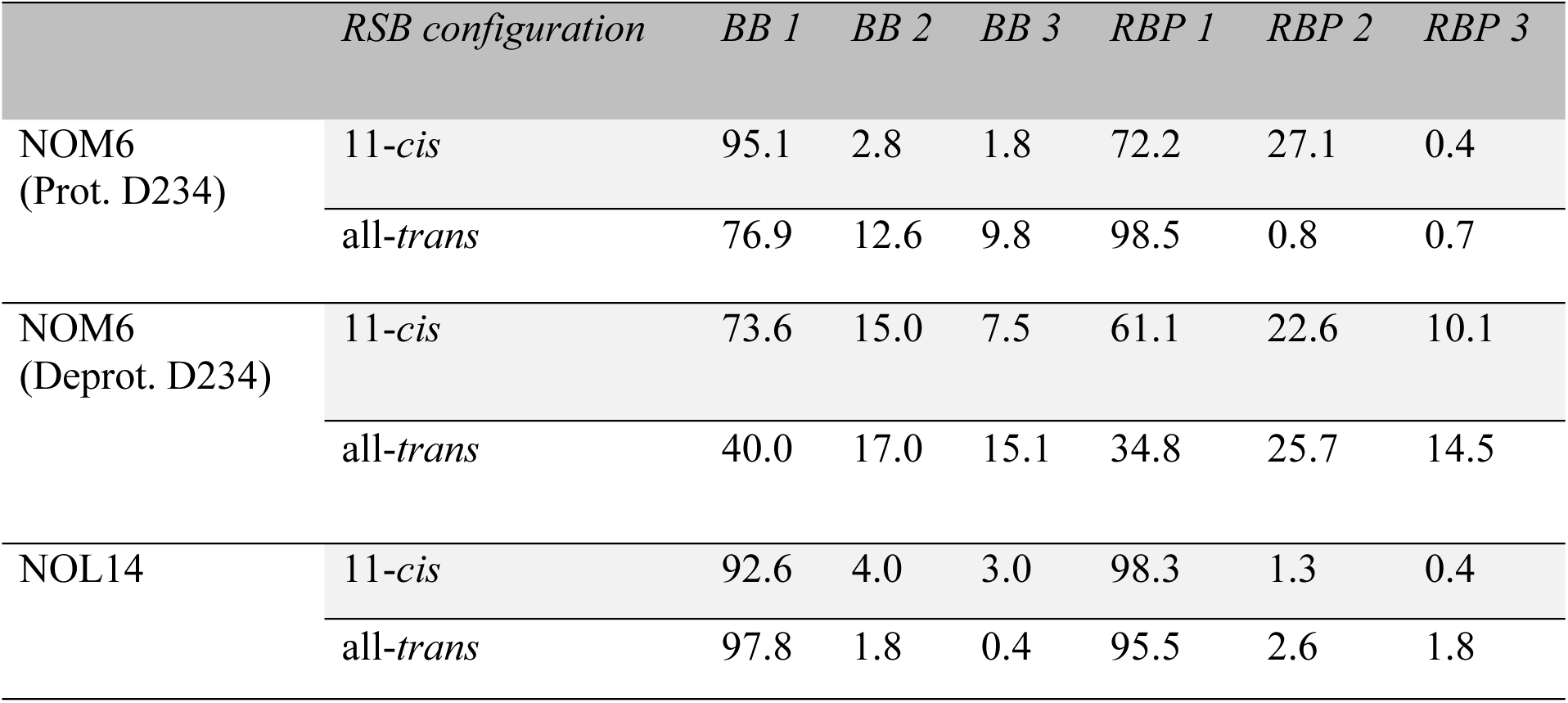
Cluster population (%) of ClusterBB and ClusterRBP. Cluster populations of ClusterBB (1-3) and ClusterRBP (1-3) were derived from 500-ns molecular dynamic simulations with three independent replicates. The percentage values represent the relative abundance of each conformational cluster obtained from trajectory-based structural clustering analysis.

## Data S1. (separate file)

Sequences used in the alignment of fig. S1, with bovine rhodopsin as reference (PDB: 1GZM_A). Sequences were obtained from previous publications (*29, 30*).

## References and Notes

1. T. W. Cronin, M. L. Porter, M. J. Bok, R. L. Caldwell, J. Marshall, Colour vision in stomatopod crustaceans. Philos. Trans. R. Soc. B Biol. Sci. 377, 20210278 (2022) 10.1098/rstb.2021.0278.

2. N. J. Marshall, M. F. Land, C. A. King, T. W. Cronin, The Compound Eyes of Mantis Shrimps (Crustacea, Hoplocarida, Stomatopoda). II. Colour Pigments in the Eyes of Stomatopod Crustaceans: Polychromatic Vision by Serial and Lateral Filtering. Philos. Trans. Biol. Sci. 334, 57–84 (1991) 10.1098/rstb.1991.0096.

3. J. Marshall, T. W. Cronin, S. Kleinlogel, Stomatopod eye structure and function: a review. Arthropod Struct. Dev. 36, 420–448 (2007) 10.1016/j.asd.2007.01.006.

4. T. W. Cronin, N. J. Marshall, C. A. Quinn, C. A. King, Ultraviolet photoreception in mantis shrimp. Vision Res. 34, 1443–1452 (1994) 10.1016/0042-6989(94)90145-7.

5. T. W. Cronin, N. J. Marshall, Multiple spectral classes of photoreceptors in the retinas of gonodactyloid stomatopod crustaceans. J. Comp. Physiol. A 166, 261–275 (1989) 10.1016/0042-6989(94)90145-7.

6. T. Cronin, N. J. Marshall, R. L. Caldwell, N. Shashar, Specialization of retinal function in the compound eyes of mantis shrimps. Vision Res. 34, 2639–2656 (1994) 10.1016/0042-6989(94)90221-6.

7. J. Marshall, J. Oberwinkler, The colourful world of the mantis shrimp. Nature 401, 873–874 (1999) 10.1038/44751.

8. H. H. Thoen, N. J. Strausfeld, J. Marshall, Neural organization of afferent pathways from the stomatopod compound eye. J. Comp. Neurol. 525, 3010–3030 (2017) 10.1002/cne.24256.

9. A. Streets, H. England, J. Marshall, Colour vision in stomatopod crustaceans: more questions than answers. J. Exp. Biol. 225, jeb243699 (2022) 10.1242/jeb.243699.

10. C.-W. J. Wang, N. J. Marshall, Behavioural evidence of spectral opponent processing in the visual system of stomatopod crustaceans. J. Exp. Biol. 228, jeb247952 (2025) 10.1242/jeb.247952.

11. T. W. Cronin, M. J. Bok, N. J. Marshall, R. L. Caldwell, Filtering and polychromatic vision in mantis shrimps: themes in visible and ultraviolet vision. Philos. Trans. R. Soc. B Biol. Sci. 369, 20130032 (2014) 10.1098/rstb.2013.0032.

12. M. J. Bok, M. L. Porter, T. W. Cronin, Ultraviolet filters in stomatopod crustaceans: diversity, ecology and evolution. J. Exp. Biol. 218, 2055–2066 (2015) 10.1242/jeb.122036.

13. N. S. Roberts, J. F. D. Hagen, R. J. Johnston, The diversity of invertebrate visual opsins spanning Protostomia, Deuterostomia, and Cnidaria. Dev. Biol. 492, 187–199 (2022) 10.1016/j.ydbio.2022.10.011.

14. T. W. Cronin, S. Johnsen, J. Marshall, E. J. Warrant, Visual Ecology (Princeton University Press (2014) ISBN 978-0-691-15184-7.

15. M. A. Liénard, G. D. Bernard, A. Allen, J.-M. Lassance, S. Song, R. R. Childers, N. Yu, D. Ye, A. Stephenson, W. A. Valencia-Montoya, S. Salzman, M. R. L. Whitaker, M. Calonje, F. Zhang, N. E. Pierce, The evolution of red color vision is linked to coordinated rhodopsin tuning in lycaenid butterflies. Proc. Natl. Acad. Sci. 118, e2008986118 (2021) 10.1073/pnas.2008986118.

16. S. L. Merbs, J. Nathans, Absorption spectra of human cone pigments. Nature 356, 433–435 (1992) 10.1038/356433a0.

17. J. W. L. Parry, K. L. Carleton, T. Spady, A. Carboo, D. M. Hunt, J. K. Bowmaker, Mix and Match Color Vision: Tuning Spectral Sensitivity by Differential Opsin Gene Expression in Lake Malawi Cichlids. Curr. Biol. 15, 1734–1739 (2005) 10.1016/j.cub.2005.08.010.

18. C. R. Sharkey, J. Blanco, N. P. Lord, T. J. Wardill, Jewel Beetle Opsin Duplication and Divergence Is the Mechanism for Diverse Spectral Sensitivities. Mol. Biol. Evol. 40, msad023 (2023) 10.1093/molbev/msad023.

19. T. C. Spady, J. W. L. Parry, P. R. Robinson, D. M. Hunt, J. K. Bowmaker, K. L. Carleton, Evolution of the Cichlid Visual Palette through Ontogenetic Subfunctionalization of the Opsin Gene Arrays. Mol. Biol. Evol. 23, 1538–1547 (2006) 10.1093/molbev/msl014.

20. S. Yokoyama, Molecular evolution of vertebrate visual pigments. Prog. Retin. Eye Res. 19, 385–419 (2000) 10.1016/S1350-9462(00)00002-1.

21. J. C. Corbo, Vitamin A1/A2 chromophore exchange: Its role in spectral tuning and visual plasticity. Dev. Biol. 475, 145–155 (2021) 10.1016/j.ydbio.2021.03.002.

22. V. H. Corredor, E. Hauzman, A. da S. Gonçalves, D. F. Ventura, Genetic characterization of the visual pigments of the red-eared turtle (Trachemys scripta elegans) and computational predictions of the spectral sensitivity. J. Photochem. Photobiol. 12, 100141 (2022) 10.1016/j.jpap.2022.100141.

23. J. M. Enright, M. B. Toomey, S. Sato, S. E. Temple, J. R. Allen, R. Fujiwara, V. M. Kramlinger, L. D. Nagy, K. M. Johnson, Y. Xiao, M. J. How, S. L. Johnson, N. W. Roberts, V. J. Kefalov, F. P. Guengerich, J. C. Corbo, Cyp27c1 Red-Shifts the Spectral Sensitivity of Photoreceptors by Converting Vitamin A1 into A2. Curr. Biol. 25, 3048–3057 (2015) 10.1016/j.cub.2015.10.018.

24. D. Escobar-Camacho, M. E. R. Pierotti, V. Ferenc, D. M. T. Sharpe, E. Ramos, C. Martins, K. L. Carleton, Variable vision in variable environments: the visual system of an invasive cichlid (Cichla monoculus) in Lake Gatun, Panama. J. Exp. Biol. 222, jeb188300 (2019) 10.1242/jeb.188300.

25. E. R. Loew, V. I. Govardovskii, Photoreceptors and visual pigments in the red-eared turtle, Trachemys scripta elegans. Vis. Neurosci. 18, 753–757 (2001) 10.1017/s0952523801185081.

26. J. C. Partridge, R. H. Douglas, Far-red sensitivity of dragon fish. Nature 375, 21–22 (1995) 10.1038/375021a0.

27. Y. Imamoto, Y. Shichida, Cone visual pigments. Biochim. Biophys. Acta BBA - Bioenerg. 1837, 664–673 (2014) 10.1016/j.bbabio.2013.08.009.

28. C. J. van der Kooi, D. G. Stavenga, K. Arikawa, G. Belušič, A. Kelber, Evolution of Insect Color Vision: From Spectral Sensitivity to Visual Ecology. Annu. Rev. Entomol. 66, 435–461 (2021) 10.1146/annurev-ento-061720-071644.

29. M. L. Porter, H. Awata, M. J. Bok, T. W. Cronin, Exceptional diversity of opsin expression patterns in Neogonodactylus oerstedii (Stomatopoda) retinas. Proc. Natl. Acad. Sci. 117, 8948–8957 (2020) 10.1073/pnas.1917303117.

30. M. W. Donohue, K. L. Carleton, T. W. Cronin, Opsin Expression in the Central Nervous System of the Mantis Shrimp Neogonodactylus oerstedii. Biol. Bull. 233, 58–69 (2017) 10.1086/694421.

31. I. G. Gleadall, T. Hariyama, Y. Tsukahara, The visual pigment chromophores in the retina of insect compound eyes, with special reference to the Coleoptera. J. Insect Physiol. 35, 787–795 (1989) 10.1016/0022-1910(89)90137-6.

32. T. H. Goldsmith, T. W. Cronin, The retinoids of seven species of mantis shrimp. Vis. Neurosci. 10, 915–920 (1993) 10.1017/s095252380000612x.

33. T. Suzuki, E. Eguchi, A survey of 3-dehydroretinal as a visual pigment chromophore in various species of crayfish and other freshwater crustaceans. Experientia 43, 1111–1113 (1987) 10.1007/BF01956053.

34. T. W. Cronin, N. J. Marshall, R. L. Caldwell, Visual pigment diversity in two genera of mantis shrimps implies rapid evolution (Crustacea; Stomatopoda). J. Comp. Physiol. A 179, 371–384 (1996) 10.1007/BF00194991.

35. R. J. L. Helena J. Bailes, Human melanopsin forms a pigment maximally sensitive to blue light (λmax ≈ 479 nm) supporting activation of Gq/11 and Gi/o signalling cascades | Proceedings of the Royal Society B: Biological Sciences. Proc. R. Soc. B., doi: 10.1098/rspb.2012.2987 (2013).

36. A. Inoue, F. Raimondi, F. M. N. Kadji, G. Singh, T. Kishi, A. Uwamizu, Y. Ono, Y. Shinjo, S. Ishida, N. Arang, K. Kawakami, J. S. Gutkind, J. Aoki, R. B. Russell, Illuminating G-Protein-Coupling Selectivity of GPCRs. Cell 177, 1933–1947.e25 (2019) 10.1016/j.cell.2019.04.044.

37. E. R. Ballister, J. Rodgers, F. Martial, R. J. Lucas, A live cell assay of GPCR coupling allows identification of optogenetic tools for controlling Go and Gi signaling. BMC Biol. 16, 10 (2018) 10.1186/s12915-017-0475-2.

38. A. Inoue, J. Ishiguro, H. Kitamura, N. Arima, M. Okutani, A. Shuto, S. Higashiyama, T. Ohwada, H. Arai, K. Makide, J. Aoki, TGFα shedding assay: an accurate and versatile method for detecting GPCR activation. Nat. Methods 9, 1021–1029 (2012) 10.1038/nmeth.2172.

39. R. J. McDowell, A. Didikoglu, T. Woelders, M. J. Gatt, F. Moffatt, S. Notash, R. A. Hut, T. M. Brown, R. J. Lucas, Beyond Lux: methods for species and photoreceptor-specific quantification of ambient light for mammals. BMC Biol. 22, 257 (2024) 10.1186/s12915-024-02038-1.

40. K. W. Foster, “Chapter 3 Action spectroscopy of photomovement” in Comprehensive Series in Photosciences, D.-P. Häder, A. M. Breure, Eds. Elsevier, vol. 1 of Photomovement, pp. 51–115 (2001) 10.1016/S1568-461X(01)80007-7.

41. R. G. Foster, M. W. Hankins, Circadian vision. Curr. Biol. CB 17, R746–751 (2007) 10.1016/j.cub.2007.07.007.

42. M. Koyanagi, B. Shen, T. Nagata, L. Sun, S. Wada, S. Kamimura, E. Kage-Nakadai, A. Terakita, High-performance optical control of GPCR signaling by bistable animal opsins MosOpn3 and LamPP in a molecular property–dependent manner. Proc. Natl. Acad. Sci. 119, e2204341119 (2022) 10.1073/pnas.2204341119.

43. A. Wagdi, D. Malan, U. Sathyanarayanan, J. S. Beauchamp, M. Vogt, D. Zipf, T. Beiert, B. Mansuroglu, V. Dusend, M. Meininghaus, L. Schneider, B. Kalthof, J. S. Wiegert, G. M. König, E. Kostenis, R. Patejdl, P. Sasse, T. Bruegmann, Selective optogenetic control of Gq signaling using human Neuropsin. Nat. Commun. 13, 1765 (2022) 10.1038/s41467-022-29265-w.

44. H. J. Bailes, L.-Y. Zhuang, R. J. Lucas, Reproducible and Sustained Regulation of Gαs Signalling Using a Metazoan Opsin as an Optogenetic Tool. PLOS ONE 7, e30774 (2012) 10.1371/journal.pone.0030774.

45. J. Abramson, J. Adler, J. Dunger, R. Evans, T. Green, A. Pritzel, O. Ronneberger, L. Willmore, A. J. Ballard, J. Bambrick, S. W. Bodenstein, D. A. Evans, C.-C. Hung, M. O’Neill, D. Reiman, K. Tunyasuvunakool, Z. Wu, A. Žemgulytė, E. Arvaniti, C. Beattie, O. Bertolli, A. Bridgland, A. Cherepanov, M. Congreve, A. I. Cowen-Rivers, A. Cowie, M. Figurnov, F. B. Fuchs, H. Gladman, R. Jain, Y. A. Khan, C. M. R. Low, K. Perlin, A. Potapenko, P. Savy, S. Singh, A. Stecula, A. Thillaisundaram, C. Tong, S. Yakneen, E. D. Zhong, M. Zielinski, A. Žídek, V. Bapst, P. Kohli, M. Jaderberg, D. Hassabis, J. M. Jumper, Accurate structure prediction of biomolecular interactions with AlphaFold 3. Nature 630, 493–500 (2024) 10.1038/s41586-024-07487-w.

46. R. A. Stein, H. S. Mchaourab, SPEACH_AF: Sampling protein ensembles and conformational heterogeneity with Alphafold2. PLOS Comput. Biol. 18, e1010483 (2022) 10.1371/journal.pcbi.1010483.

47. B. Honig, U. Dinur, K. Nakanishi, V. Balogh-Nair, M. A. Gawinowicz, M. Arnaboldi, M. G. Motto, An external point-charge model for wavelength regulation in visual pigments. J. Am. Chem. Soc. 101, 7084–7086 (1979) 10.1021/ja00517a060.

48. K. Nakanishi, V. Balogh-Nair, M. Arnaboldi, K. Tsujimoto, B. Honig, An external point-charge model for bacteriorhodopsin to account for its purple color. J. Am. Chem. Soc. 102, 7945–7947 (1980) 10.1021/ja00547a028.

49. E. J. Warrant, S. Johnsen, Vision and the light environment. Curr. Biol. 23, R990–R994 (2013) 10.1016/j.cub.2013.10.019.

50. G. D. Bernard, Red-Absorbing Visual Pigment of Butterflies. Science 203, 1125–1127 (1979) 10.1126/science.203.4385.1125.

51. T. D. Lamb, Photoreceptor spectral sensitivities: common shape in the long-wavelength region. Vision Res. 35, 3083–3091 (1995) 10.1016/0042-6989(95)00114-f.

52. T.-H. Chiou, A. R. Place, R. L. Caldwell, N. J. Marshall, T. W. Cronin, A novel function for a carotenoid: astaxanthin used as a polarizer for visual signalling in a mantis shrimp. J. Exp. Biol. 215, 584–589 (2012) 10.1242/jeb.066019.

53. C. H. Mazel, T. W. Cronin, R. L. Caldwell, N. J. Marshall, Fluorescent Enhancement of Signaling in a Mantis Shrimp. Science 303, 51–51 (2004) 10.1126/science.1089803.

54. A. M. Franklin, M. B. Applegate, S. M. Lewis, F. G. Omenetto, Stomatopods detect and assess achromatic cues in contests. Behav. Ecol. 28, 1329–1336 (2017) 10.1093/beheco/arx096.

55. A. M. Franklin, J. Marshall, A. D. Feinstein, M. J. Bok, A. D. Byrd, S. M. Lewis, Differences in signal contrast and camouflage among different colour variations of a stomatopod crustacean, Neogonodactylus oerstedii. Sci. Rep. 10, 1236 (2020) 10.1038/s41598-020-57990-z.

56. M. W. Donohue, J. H. Cohen, T. W. Cronin, Cerebral photoreception in mantis shrimp. Sci. Rep. 8, 9689 (2018) 10.1038/s41598-018-28004-w.

57. R. R. Rando, Molecular mechanisms in visual pigment regeneration. Photochem. Photobiol. 56, 1145–1156 (1992) 10.1111/j.1751-1097.1992.tb09739.x.

58. A. Morshedian, J. J. Kaylor, S. Y. Ng, A. Tsan, R. Frederiksen, T. Xu, L. Yuan, A. P. Sampath, R. A. Radu, G. L. Fain, G. H. Travis, Light-Driven Regeneration of Cone Visual Pigments through a Mechanism Involving RGR Opsin in Müller Glial Cells. Neuron 102, 1172–1183.e5 (2019) 10.1016/j.neuron.2019.04.004.

59. K. Sakai, Y. Shichida, Y. Imamoto, T. Yamashita, Creation of photocyclic vertebrate rhodopsin by single amino acid substitution. eLife 11, e75979 (2022) 10.7554/eLife.75979.

60. M. Koyanagi, S. Wada, E. Kawano-Yamashita, Y. Hara, S. Kuraku, S. Kosaka, K. Kawakami, S. Tamotsu, H. Tsukamoto, Y. Shichida, A. Terakita, Diversification of non-visual photopigment parapinopsin in spectral sensitivity for diverse pineal functions. BMC Biol. 13, 73 (2015) 10.1186/s12915-015-0174-9.

61. J. Wietek, A. Nozownik, M. Pulin, I. Saraf-Sinik, N. Matosevich, R. Gowrishankar, A. Gat, D. Malan, B. J. Brown, J. Dine, B. N. Imambocus, R. Levy, K. Sauter, A. Litvin, N. Regev, S. Subramaniam, K. Abrera, D. Summarli, E. M. Goren, G. Mizrachi, E. Bitton, A. Benjamin, B. A. Copits, P. Sasse, B. R. Rost, D. Schmitz, M. R. Bruchas, P. Soba, M. Oren-Suissa, Y. Nir, J. S. Wiegert, O. Yizhar, A bistable inhibitory optoGPCR for multiplexed optogenetic control of neural circuits. Nat. Methods 21, 1275–1287 (2024) 10.1038/s41592-024-02285-8.

62. T. Nagata, M. Koyanagi, H. Tsukamoto, E. Mutt, G. F. X. Schertler, X. Deupi, A. Terakita, The counterion–retinylidene Schiff base interaction of an invertebrate rhodopsin rearranges upon light activation. Commun. Biol. 2, 180 (2019) 10.1038/s42003-019-0409-3.

63. M. J. Rodrigues, O. Tejero, J. Mühle, F. Pamula, I. Das, C.-J. Tsai, A. Terakita, M. Sheves, G. F. X. Schertler, Activating an invertebrate bistable opsin with the all-trans 6.11 retinal analog. Proc. Natl. Acad. Sci. 121, e2406814121 (2024) 10.1073/pnas.2406814121.

64. A. Terakita, R. Hara, T. Hara, Retinal-binding protein as a shuttle for retinal in the rhodopsin-retinochrome system of the squid visual cells. Vision Res. 29, 639–652 (1989) 10.1016/0042-6989(89)90026-6.

65. M. A. Larkin, G. Blackshields, N. P. Brown, R. Chenna, P. A. McGettigan, H. McWilliam, F. Valentin, I. M. Wallace, A. Wilm, R. Lopez, J. D. Thompson, T. J. Gibson, D. G. Higgins, Clustal W and Clustal X version 2.0. Bioinformatics 23, 2947–2948 (2007) 10.1093/bioinformatics/btm404.

66. D. G. Gibson, L. Young, R.-Y. Chuang, J. C. Venter, C. A. Hutchison, H. O. Smith, Enzymatic assembly of DNA molecules up to several hundred kilobases. Nat. Methods 6, 343–345 (2009) 10.1038/nmeth.1318.

67. C.-F. Bowin, A. Inoue, G. Schulte, WNT-3A–induced β-catenin signaling does not require signaling through heterotrimeric G proteins. J. Biol. Chem. 294, 11677–11684 (2019) 10.1074/jbc.AC119.009412.

68. V. I. Govardovskii, N. Fyhrquist, T. Reuter, D. G. Kuzmin, K. Donner, In search of the visual pigment template. Vis. Neurosci. 17, 509–528 (2000) 10.1017/s0952523800174036.

69. P. Virtanen, R. Gommers, T. E. Oliphant, M. Haberland, T. Reddy, D. Cournapeau, E. Burovski, P. Peterson, W. Weckesser, J. Bright, S. J. van der Walt, M. Brett, J. Wilson, K. J. Millman, N. Mayorov, A. R. J. Nelson, E. Jones, R. Kern, E. Larson, C. J. Carey, İ. Polat, Y. Feng, E. W. Moore, J. VanderPlas, D. Laxalde, J. Perktold, R. Cimrman, I. Henriksen, E. A. Quintero, C. R. Harris, A. M. Archibald, A. H. Ribeiro, F. Pedregosa, P. van Mulbregt, SciPy 1.0: fundamental algorithms for scientific computing in Python. Nat. Methods 17, 261–272 (2020) 10.1038/s41592-019-0686-2.

70. F. Pedregosa, G. Varoquaux, A. Gramfort, V. Michel, B. Thirion, O. Grisel, M. Blondel, A. Müller, J. Nothman, G. Louppe, P. Prettenhofer, R. Weiss, V. Dubourg, J. Vanderplas, A. Passos, D. Cournapeau, M. Brucher, M. Perrot, É. Duchesnay, Scikit-learn: Machine Learning in Python. arXiv arXiv:1201.0490 [Preprint] (2018). 10.48550/arXiv.1201.0490.

71. L. S. Brown, Fungal rhodopsins and opsin-related proteins: eukaryotic homologues of bacteriorhodopsin with unknown functions. Photochem. Photobiol. Sci. 3, 555 (2004) 10.1039/b315527g.

72. J. Schindelin, I. Arganda-Carreras, E. Frise, V. Kaynig, M. Longair, T. Pietzsch, S. Preibisch, C. Rueden, S. Saalfeld, B. Schmid, J.-Y. Tinevez, D. J. White, V. Hartenstein, K. Eliceiri, P. Tomancak, A. Cardona, Fiji: an open-source platform for biological-image analysis. Nat. Methods 9, 676–682 (2012) 10.1038/nmeth.2019.

73. M. Mirdita, K. Schütze, Y. Moriwaki, L. Heo, S. Ovchinnikov, M. Steinegger, ColabFold: making protein folding accessible to all. Nat. Methods 19, 679–682 (2022) 10.1038/s41592-022-01488-1.

74. C. J. Williams, J. J. Headd, N. W. Moriarty, M. G. Prisant, L. L. Videau, L. N. Deis, V. Verma, D. A. Keedy, B. J. Hintze, V. B. Chen, S. Jain, S. M. Lewis, W. B. Arendall III, J. Snoeyink, P. D. Adams, S. C. Lovell, J. S. Richardson, D. C. Richardson, MolProbity: More and better reference data for improved all-atom structure validation. Protein Sci. 27, 293–315 (2018) 10.1002/pro.3330.

75. L. S. Johnson, S. R. Eddy, E. Portugaly, Hidden Markov model speed heuristic and iterative HMM search procedure. BMC Bioinformatics 11, 431 (2010) 10.1186/1471-2105-11-431.

76. M. Steinegger, J. Söding, MMseqs2 enables sensitive protein sequence searching for the analysis of massive data sets. Nat. Biotechnol. 35, 1026–1028 (2017) 10.1038/nbt.3988.

77. D. Liebschner, P. V. Afonine, M. L. Baker, G. Bunkóczi, V. B. Chen, T. I. Croll, B. Hintze, L.-W. Hung, S. Jain, A. J. McCoy, N. W. Moriarty, R. D. Oeffner, B. K. Poon, M. G. Prisant, R. J. Read, J. S. Richardson, D. C. Richardson, M. D. Sammito, O. V. Sobolev, D. H. Stockwell, T. C. Terwilliger, A. G. Urzhumtsev, L. L. Videau, C. J. Williams, P. D. Adams, Macromolecular structure determination using X-rays, neutrons and electrons: recent developments in Phenix. Acta Crystallogr. Sect. Struct. Biol. 75, 861–877 (2019) 10.1107/S2059798319011471.

78. A. Šali, T. L. Blundell, Comparative Protein Modelling by Satisfaction of Spatial Restraints. J. Mol. Biol. 234, 779–815 (1993) 10.1006/jmbi.1993.1626.

79. G. Jones, P. Willett, R. C. Glen, A. R. Leach, R. Taylor, Development and validation of a genetic algorithm for flexible docking1. J. Mol. Biol. 267, 727–748 (1997) 10.1006/jmbi.1996.0897.

80. M. Murakami, T. Kouyama, Crystal structure of squid rhodopsin. Nature 453, 363–367 (2008) 10.1038/nature06925.

81. T. Gruhl, T. Weinert, M. J. Rodrigues, C. J. Milne, G. Ortolani, K. Nass, E. Nango, S. Sen, P. J. M. Johnson, C. Cirelli, A. Furrer, S. Mous, P. Skopintsev, D. James, F. Dworkowski, P. Båth, D. Kekilli, D. Ozerov, R. Tanaka, H. Glover, C. Bacellar, S. Brünle, C. M. Casadei, A. D. Diethelm, D. Gashi, G. Gotthard, R. Guixà-González, Y. Joti, V. Kabanova, G. Knopp, E. Lesca, P. Ma, I. Martiel, J. Mühle, S. Owada, F. Pamula, D. Sarabi, O. Tejero, C.-J. Tsai, N. Varma, A. Wach, S. Boutet, K. Tono, P. Nogly, X. Deupi, S. Iwata, R. Neutze, J. Standfuss, G. Schertler, V. Panneels, Ultrafast structural changes direct the first molecular events of vision. Nature 615, 939–944 (2023) 10.1038/s41586-023-05863-6.

82. S. Zhang, J. M. Krieger, Y. Zhang, C. Kaya, B. Kaynak, K. Mikulska-Ruminska, P. Doruker, H. Li, I. Bahar, ProDy 2.0: increased scale and scope after 10 years of protein dynamics modelling with Python. Bioinformatics 37, 3657–3659 (2021) 10.1093/bioinformatics/btab187.

83. M. J. Abraham, T. Murtola, R. Schulz, S. Páll, J. C. Smith, B. Hess, E. Lindahl, GROMACS: High performance molecular simulations through multi-level parallelism from laptops to supercomputers. SoftwareX 1–2, 19–25 (2015) 10.1016/j.softx.2015.06.001.

84. J. Huang, S. Rauscher, G. Nawrocki, T. Ran, M. Feig, B. L. de Groot, H. Grubmüller, A. D. MacKerell, CHARMM36m: an improved force field for folded and intrinsically disordered proteins. Nat. Methods 14, 71–73 (2017) 10.1038/nmeth.4067.

85. W. L. Jorgensen, J. Chandrasekhar, J. D. Madura, R. W. Impey, M. L. Klein, Comparison of simple potential functions for simulating liquid water. J. Chem. Phys. 79, 926–935 (1983) 10.1063/1.445869.

86. S. Jo, T. Kim, V. G. Iyer, W. Im, CHARMM-GUI: A web-based graphical user interface for CHARMM. J. Comput. Chem. 29, 1859–1865 (2008) 10.1002/jcc.20945.

87. E. Tajkhorshid, B. Paizs, S. Suhai, Conformational Effects on the Proton Affinity of the Schiff Base in Bacteriorhodopsin: A Density Functional Study. J. Phys. Chem. B 101, 8021–8028 (1997) 10.1021/jp971283t.

88. E. Tajkhorshid, J. Baudry, K. Schulten, S. Suhai, Molecular Dynamics Study of the Nature and Origin of Retinal’s Twisted Structure in Bacteriorhodopsin. Biophys. J. 78, 683–693 (2000) 10.1016/S0006-3495(00)76626-4.

89. E. Tajkhorshid, S. Suhai, Influence of the Methyl Groups on the Structure, Charge Distribution, and Proton Affinity of the Retinal Schiff Base. J. Phys. Chem. B 103, 5581–5590 (1999) 10.1021/jp983742b.

90. T. Darden, D. York, L. Pedersen, Particle mesh Ewald: An N⋅log(N) method for Ewald sums in large systems. J. Chem. Phys. 98, 10089–10092 (1993) 10.1063/1.464397.

91. G. Bussi, D. Donadio, M. Parrinello, Canonical sampling through velocity rescaling. J. Chem. Phys. 126, 014101 (2007) 10.1063/1.2408420.

92. H. J. C. Berendsen, J. P. M. Postma, W. F. van Gunsteren, A. DiNola, J. R. Haak, Molecular dynamics with coupling to an external bath. J. Chem. Phys. 81, 3684–3690 (1984) 10.1063/1.448118.

93. M. Parrinello, Crystal Structure and Pair Potentials: A Molecular-Dynamics Study. Phys. Rev. Lett. 45, 1196–1199 (1980) 10.1103/PhysRevLett.45.1196.

94. X. Daura, K. Gademann, B. Jaun, D. Seebach, W. F. van Gunsteren, A. E. Mark, Peptide Folding: When Simulation Meets Experiment. Angew. Chem. Int. Ed. 38, 236–240 (1999) 10.1002/(SICI)1521-3773(19990115)38:1/2<236::AID-ANIE236>3.0.CO;2-M.

95. R. Salomon-Ferrer, D. A. Case, R. C. Walker, An overview of the Amber biomolecular simulation package. WIREs Comput. Mol. Sci. 3, 198–210 (2013) 10.1002/wcms.1121.

96. M. Elstner, Self-consistent-charge density-functional tight-binding method for simulations of complex materials properties. Phys. Rev. B 58, 7260–7268 (1998) 10.1103/PhysRevB.58.7260.

97. M. Elstner, P. Hobza, T. Frauenheim, S. Suhai, E. Kaxiras, Hydrogen bonding and stacking interactions of nucleic acid base pairs: A density-functional-theory based treatment. J. Chem. Phys. 114, 5149–5155 (2001) 10.1063/1.1329889.

98. J. Schirmer, Beyond the random-phase approximation: A new approximation scheme for the polarization propagator. Phys. Rev. A 26, 2395–2416 (1982) 10.1103/PhysRevA.26.2395.

99. S. G. Balasubramani, G. P. Chen, S. Coriani, M. Diedenhofen, M. S. Frank, Y. J. Franzke, F. Furche, R. Grotjahn, M. E. Harding, C. Hättig, A. Hellweg, B. Helmich-Paris, C. Holzer, U. Huniar, M. Kaupp, A. Marefat Khah, S. Karbalaei Khani, T. Müller, F. Mack, B. D. Nguyen, S. M. Parker, E. Perlt, D. Rappoport, K. Reiter, S. Roy, M. Rückert, G. Schmitz, M. Sierka, E. Tapavicza, D. P. Tew, C. van Wüllen, V. K. Voora, F. Weigend, A. Wodyński, J. M. Yu, TURBOMOLE: Modular program suite for ab initio quantum-chemical and condensed-matter simulations. J. Chem. Phys. 152, 184107 (2020) 10.1063/5.0004635.

100. A. M. Waterhouse, J. B. Procter, D. M. A. Martin, M. Clamp, G. J. Barton, Jalview Version 2—a multiple sequence alignment editor and analysis workbench. Bioinformatics 25, 1189–1191 (2009) 10.1093/bioinformatics/btp033.

101. M. A. Liénard, W. A. Valencia-Montoya, N. E. Pierce, Molecular advances to study the function, evolution and spectral tuning of arthropod visual opsins. Philos. Trans. R. Soc. B Biol. Sci., doi: 10.1098/rstb.2021.0279 (2022) 10.1098/rstb.2021.0279.

102. Q. Zhou, D. Yang, M. Wu, Y. Guo, W. Guo, L. Zhong, X. Cai, A. Dai, W. Jang, E. I. Shakhnovich, Z.-J. Liu, R. C. Stevens, N. A. Lambert, M. M. Babu, M.-W. Wang, S. Zhao, Common activation mechanism of class A GPCRs. eLife 8, e50279 (2019) 10.7554/eLife.50279.

