## Supplementary Methods and Figures for "Spectral tuning and signaling of diverse and most red sensitive animal opsins in Mantis Shrimp eyes"

**The PDF file includes:**

Materials and Methods
Figs. S1 to S13
Tables S1 to S5

**Other Supplementary Materials for this manuscript include the following:**

Data S1

### Materials and Methods

Phylogenetic analyses

*N. oerstedii* opsin DNA sequences were obtained from NCBI and reanalysed in Geneious to exclude partial sequences (Supp. Dataset S1). Full-length opsins were selected if they met further criteria: belonging to the proposed long-wave (NOL) or middle-wave (NOM) clades and being expressed either in the colour absorbing midband rows, the lateral hemisphere of the mantis shrimp eye (NOL14), as well as in the eye stalk (LWS R3). Deduced protein sequences for the 17 selected opsins (14 NOLs, 2 NOMs and LWS R3) were aligned in ClustalX (*65*), and the phylogeny was reconstructed as reported earlier (*29*).

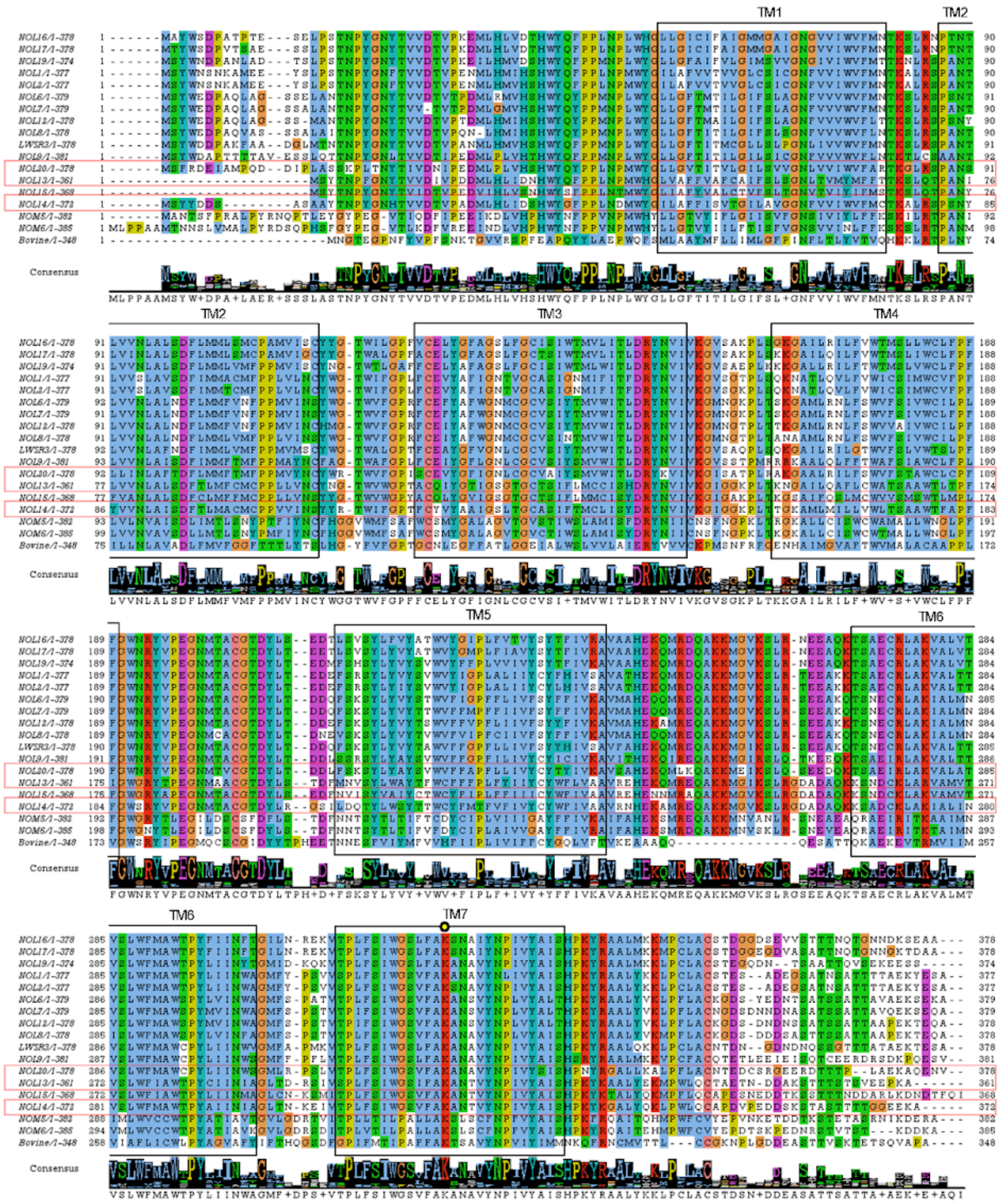


**Fig. S1.**

**Comparison of the full length NOL and NOM sequences of *N. oerstedii* with bovine rhodopsin as reference** (PDB: 1GZM_A) (*100*). Predicted transmembrane helices are marked with black rectangles based on the bovine rhodopsin model, and the retinal binding lysine is marked with a yellow circle. The most red sensitive rhodopsins are marked with red rectangles.

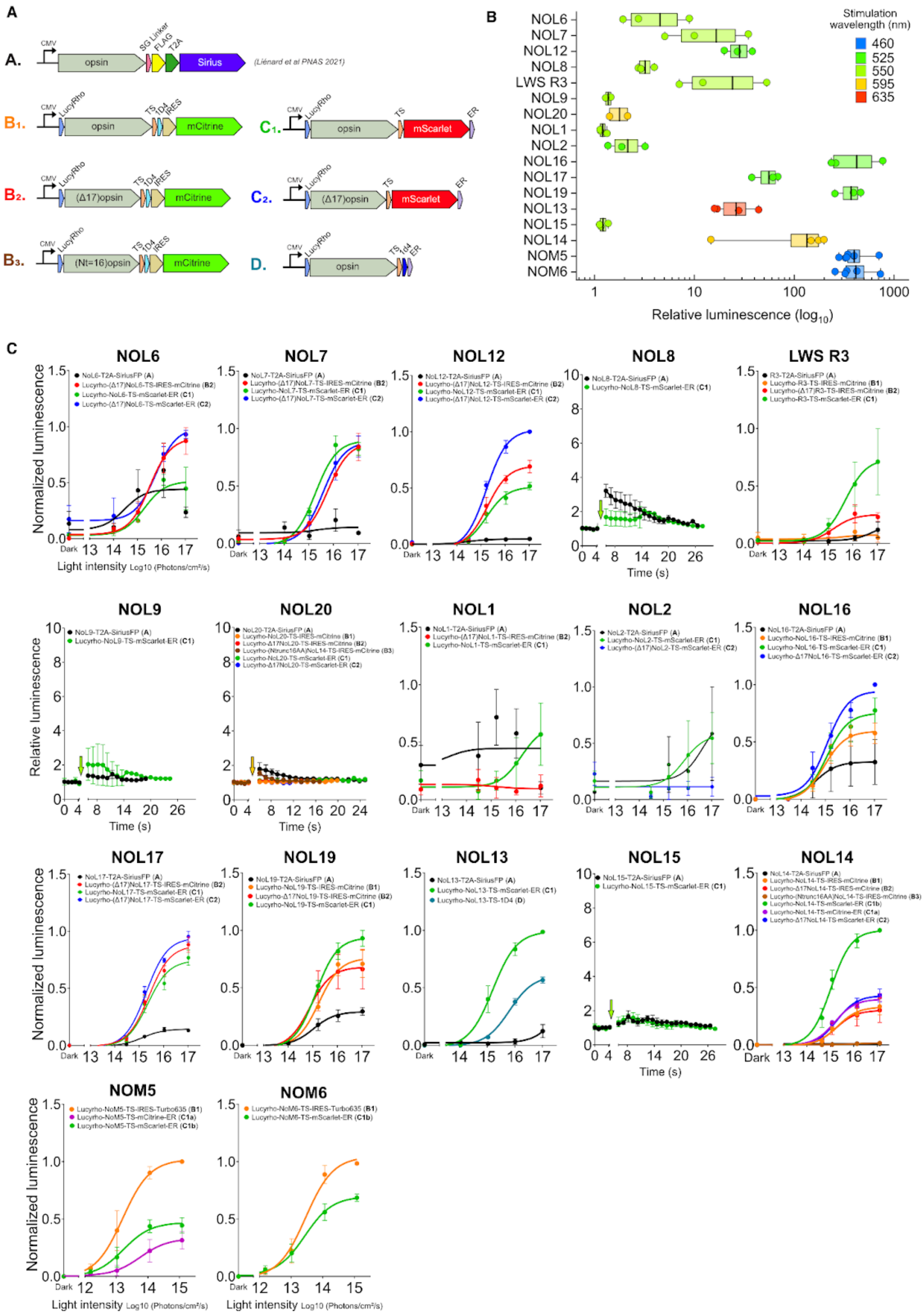

A
B
C
opsin
Sirius
mCitrine
mScarlet
(Liénard et al PNAS 2021)
Stimulation wavelength (nm)
460
525
550
595
635
Relative luminescence (log10)
NOL6
NOL7
NOL12
NOL8
LWS R3
NOL9
NOL20
NOL1
NOL2
NOL16
NOL17
NOL19
NOL13
NOL15
NOL14
NOM5
NOM6
Normalized luminescence
Relative luminescence
Light intensity Log10 (Photons/cm²/s)
Time (s)

**Fig. S2.**

**Optimisation of constructs.** (**A**) Design of DNA constructs used for optimisation of the luminescent responses (SG linker 2aa, 1D4 affinity tag TETSQVAPA 9aa, FLAG affinity tag DYKDDDDK 8aa, LucyRho self cleavable leucine-rich signal peptide, T2A Self cleaving peptide from *Thosea asigna* virus 2A GSGEGRGSLLTCGDVEENPGP 21aa, TS Golgi export trafficking signal sequence KSRITSEGEYIPLDQIDINV 20aa, ER endoplasmic reticulum export sequence FCYENEV 7aa, IRES internal ribosome entry site, Δ17 deletion of t17aa of an opsin N-terminus, Nt=16 only 16aa remaining upstream of TM1 of the opsin). Construct A. has been previously described (*101*). (**B**) Comparison of relative aequorin luminescent levels of the most effective (highest signal) constructs for each opsin following a 1s activation (wavelength indicated in the legend). (**C**) Light-titration aequorin responses of various DNA constructs used in this study. Unresponsive opsins (NOL8, NOL9, NOL15, NOL20) signals are shown as relative luminescence over time, at the highest light intensity ($1x10^{17}$ photons.cm$^{-2}$.s$^{-1}$).

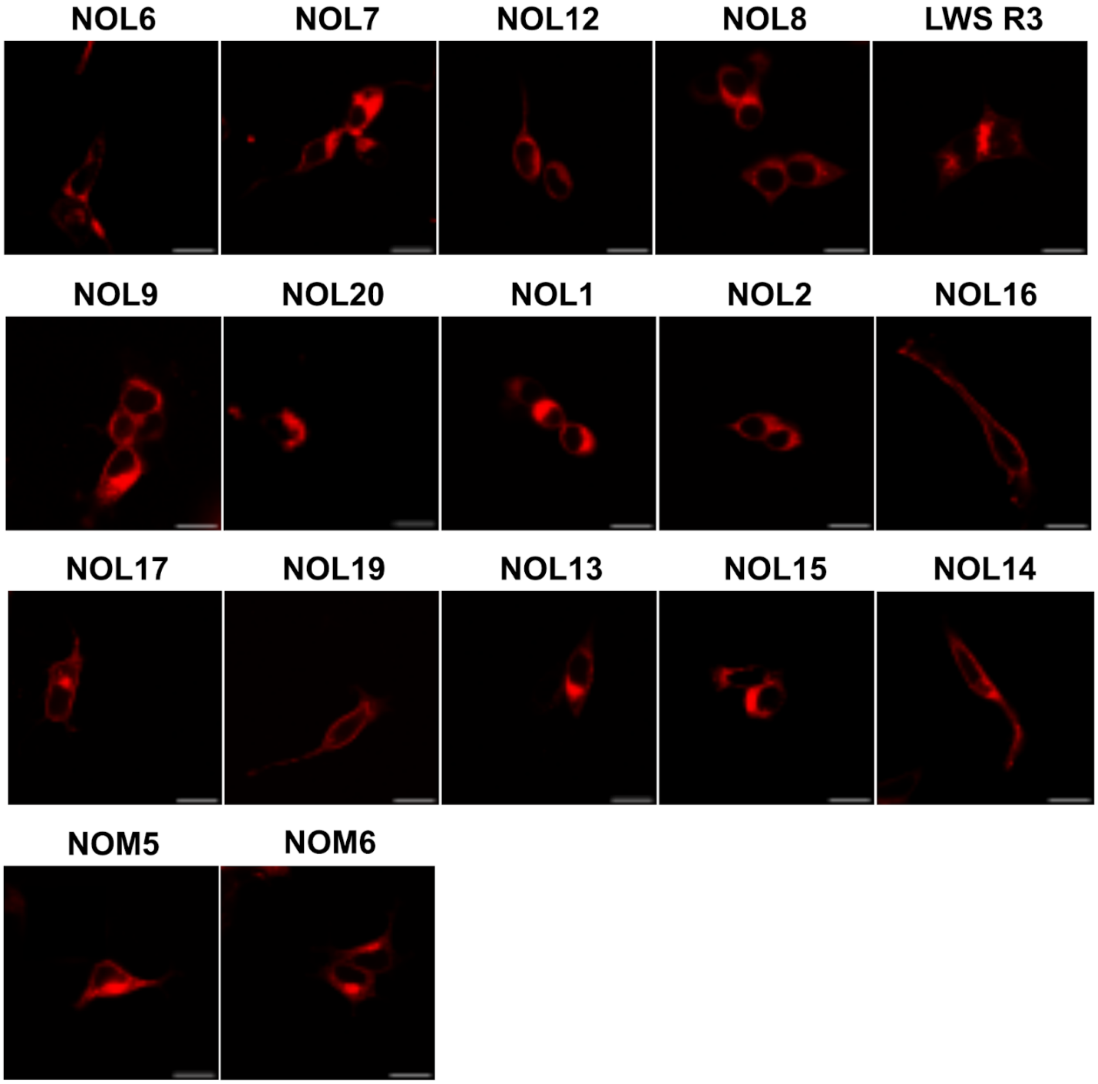


**Fig. S3.**

**Expression profile of NOM and NOLs opsins.** HEK293T cells expressing NOM or NOL opsins fused to mScarlet at the C-termini (construct C1 in fig. S2). Scale bar = 20µm.

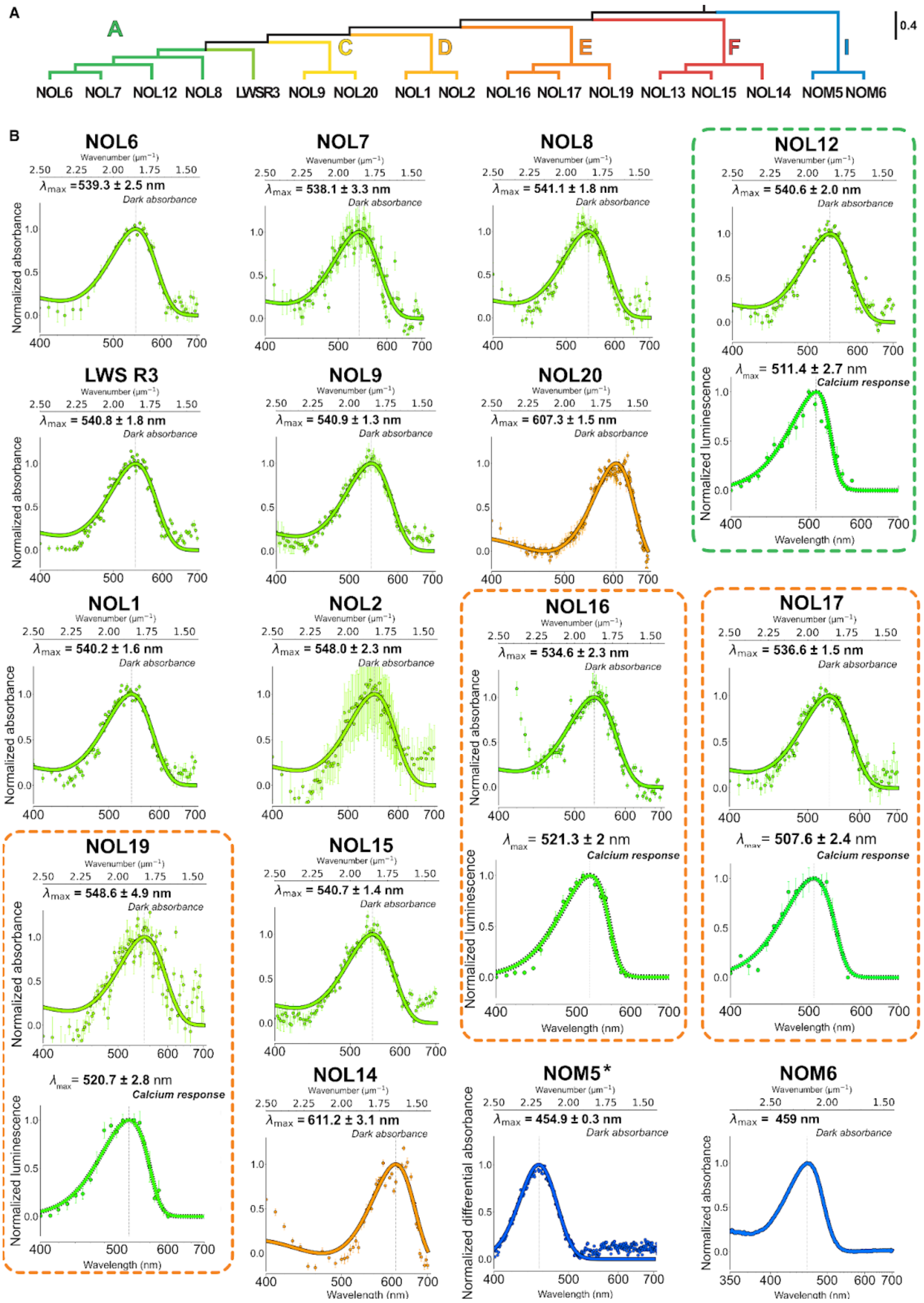
A
A
C
D
E
F
I
0.4
NOL6 NOL7 NOL12 NOL8 LWSR3 NOL9 NOL20 NOL1 NOL2 NOL16 NOL17 NOL19 NOL13 NOL15 NOL14 NOM5 NOM6
B
NOL6
$\lambda_{max}$ =539.3 ± 2.5 nm
Dark absorbance
NOL7
$\lambda_{max}$ = 538.1 ± 3.3 nm
Dark absorbance
NOL8
$\lambda_{max}$ = 541.1 ± 1.8 nm
Dark absorbance
NOL12
$\lambda_{max}$ = 540.6 ± 2.0 nm
Dark absorbance
$\lambda_{max}$ = 511.4 ± 2.7 nm
Calcium response
LWS R3
$\lambda_{max}$ = 540.8 ± 1.8 nm
Dark absorbance
NOL9
$\lambda_{max}$ = 540.9 ± 1.3 nm
Dark absorbance
NOL20
$\lambda_{max}$ = 607.3 ± 1.5 nm
Dark absorbance
NOL1
$\lambda_{max}$ = 540.2 ± 1.6 nm
Dark absorbance
NOL2
$\lambda_{max}$ = 548.0 ± 2.3 nm
Dark absorbance
NOL16
$\lambda_{max}$ = 534.6 ± 2.3 nm
Dark absorbance
$\lambda_{max}$ = 521.3 ± 2 nm
Calcium response
NOL17
$\lambda_{max}$ = 536.6 ± 1.5 nm
Dark absorbance
$\lambda_{max}$ = 507.6 ± 2.4 nm
Calcium response
NOL19
$\lambda_{max}$ = 548.6 ± 4.9 nm
Dark absorbance
$\lambda_{max}$ = 520.7 ± 2.8 nm
Calcium response
NOL15
$\lambda_{max}$ = 540.7 ± 1.4 nm
Dark absorbance
NOL14
$\lambda_{max}$ = 611.2 ± 3.1 nm
Dark absorbance
NOM5*
$\lambda_{max}$ = 454.9 ± 0.3 nm
Dark absorbance
NOM6
$\lambda_{max}$ = 459 nm
Dark absorbance
Wavenumber ($\mu m^{-1}$)
Normalized absorbance
Normalized luminescence
Normalized differential absorbance
Wavelength (nm)

**Fig. S4.**

**Dark absorption and action spectra of NOL and NOM opsins.**

(**A**) Phylogenetic relations of NOLs and NOMs from clade A, C, D, E, F and I (adapted from Porter et al. 2020). (**B**) Dark absorbance spectra measured in an UV-vis Nano-spectrometer after reconstitution with *11-cis* retinal, with corresponding action spectra below when possible, established from maximal aequorin responses, as indicated by the dashed frames, color coded according to phylogenetic branch. Light intensity of the 1 s pulse used for the action spectra : NOL12 : $8x10^{14}$ photons $cm^{-2}$ $s^{-1}$ (n=4); NOL16 (n=3) and NOL17 (n=3) : $4x10^{14}$ photons $cm^{-2}$ $s^{-1}$ ; NOL19 : $8x10^{13}$ photons $cm^{-2}$ $s^{-1}$ (n=4).Absorbance and action spectra of all NOLs were fitted with a non-linear least squares optimization of the Govardovskii visual template to estimate the best-fit $\lambda_{max}$. Data is presented as mean values and with ± SEM for NOLs. NOL6 and NOL14 spectra were previously reported (*101*). Dark absorption for NOM5 was measured with *9-cis* retinal (*) and plotted as the differential absorbance between the dark absorption and the absorption after a 60s pulse of light (455 nm), fitted with a simple gaussian. For NOM6 a continuous spectra without any fitting but corrected for Mie scattering is shown. Bimodal action spectra for NOL13, NOL14, NOM5 and NOM6 are presented in Fig. 2.

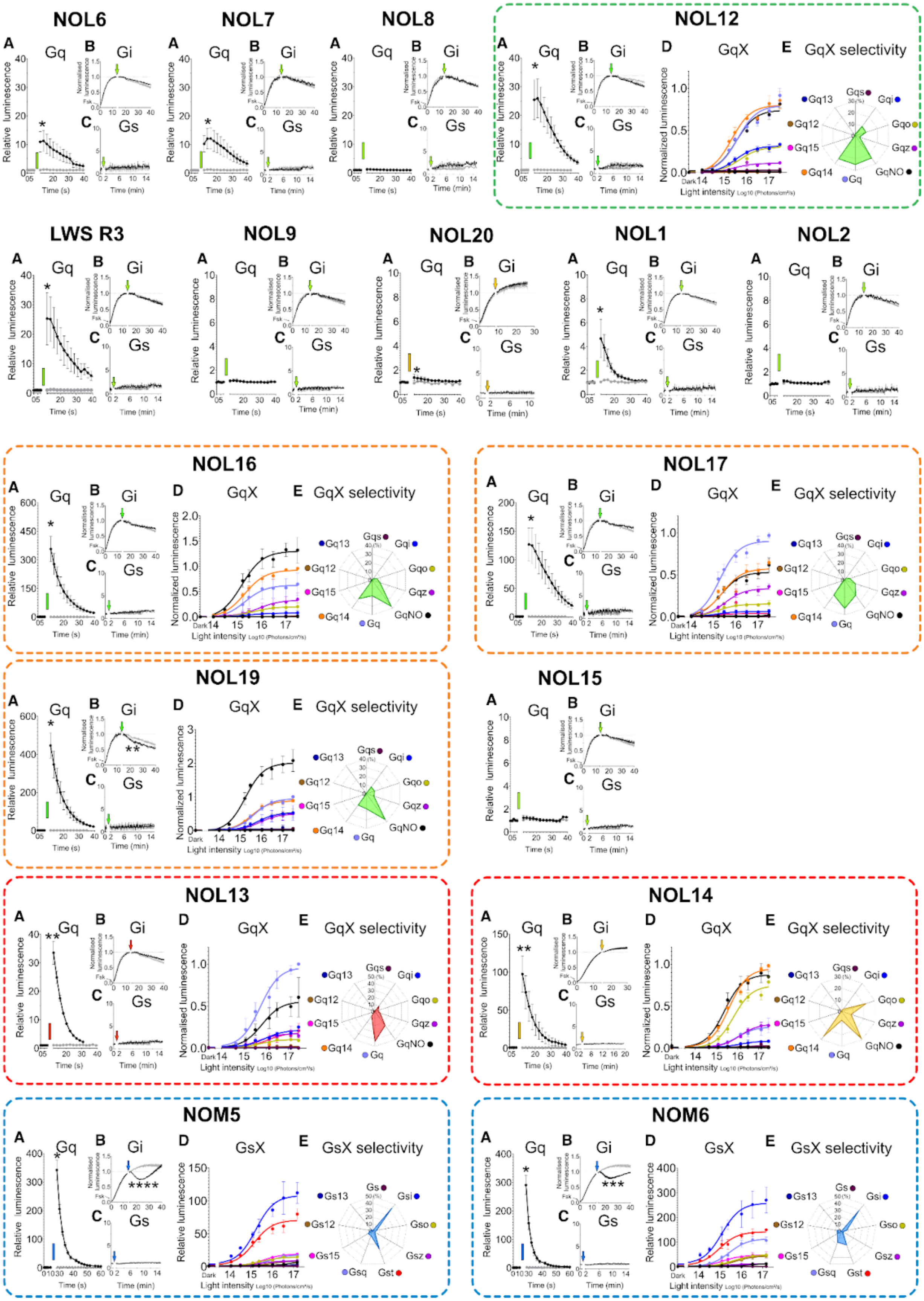
NOL6
NOL7
NOL8
NOL12
LWS R3
NOL9
NOL20
NOL1
NOL2
NOL16
NOL17
NOL19
NOL15
NOL13
NOL14
NOM5
NOM6
A Gq
B Gi
C Gs
D GqX
E GqX selectivity
D GsX
E GsX selectivity
Relative luminescence
Normalized luminescence
Time (s)
Time (min)
Light intensity Log10 (Photons/cm²/s)
Gq13
Gq12
Gq15
Gq14
Gq
Gqs
Gqi
Gqo
Gqz
GqNO
Gs13
Gs12
Gs15
Gsq
Gst
Gs
Gsi
Gso
Gsz

**Fig. S5.**

**G-protein selectivity profiles of NOL and NOM opsins.**

(A) Responses to the Gq-protein family were measured by using the aequorin assay (see Table S1 for detailed values). (B-C) Coupling efficiency to the (B) Gi- and (C) Gs-protein families in the glosensor assay. Relative luminescence values are shown as the raw luminescence divided by the averaged pre-illumination baseline. For Gi, cAMP luminescent levels were elevated using 2µM forskolin, 15 min before illumination. (D) Light-dependent G-protein signaling was further investigated for NOMs using the GsX assay and for NOLs using the GqX assay when possible (indicated by dashed frames, colourized based on their phylogenetic clade (see fig. S4). Light-dependent maximal post-stimuli responses are normalized between the no GqX/GsX control and the maximal relative response. (E) Selectivity profiles were established from maximal responses to GsX or GqX chimeras and expressed as percentages of the sum of GsX or GqX protein activity. NOL12, NOL16, NOL17, NOL19 were stimulated with 525 nm light, NOL6, NOL7, NOL8, NOL9, NOL1, NOL2, NOL15, LWS R3 with 550 nm, NOL14, NOL20 with 595 nm, and NOL13 with 635 nm. All responses were measured after a dark pre-illumination baseline followed by a 1s light pulse, indicated by colored arrows and rectangles. All data is presented as mean ± SEM. Different levels of significance at the peak are indicated by * for $p < 0.05$, ** for $p < 0.01$, *** for $p < 0.001$, and **** for $p < 0.0001$, compared to cells transfected only with luminescent reporters (grey traces).

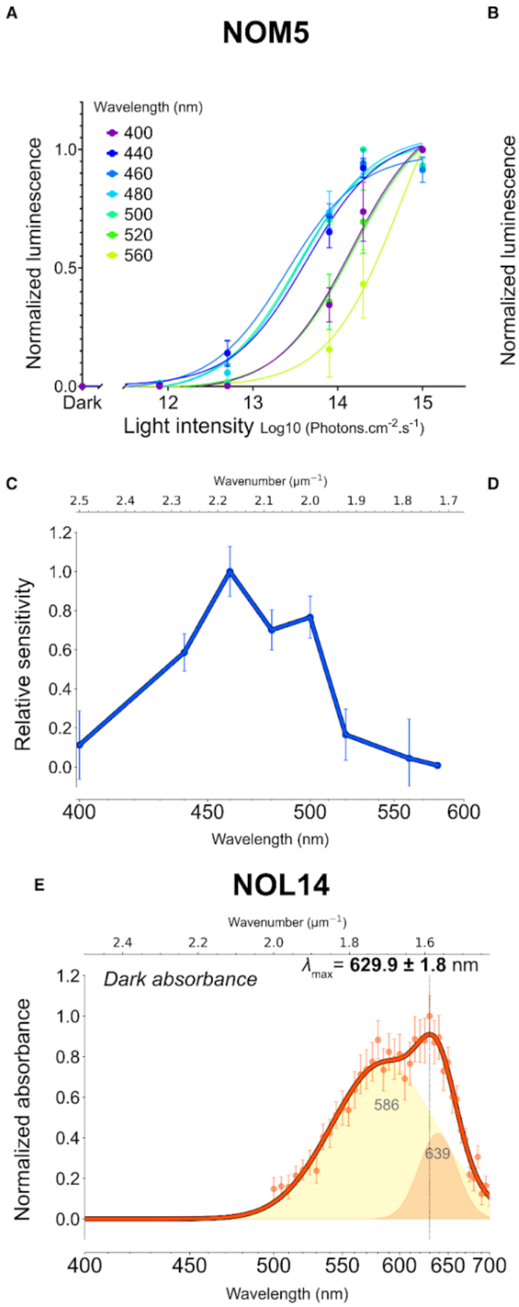


**Fig. S6.**

**Dark state spectral composition of NOM5, NOM6 and NOL14.**

Light-dependent aequorin responses following a 1s light pulse of (**A**) NOM5 and (**B**) NOM6 normalized to the maximal luminescent response per wavelength (n=3). Spectral relative sensitivity of (**C**) NOM5 and (**D**) NOM6 calculated from EC50 values extracted from a. and b. at each wavelength. (**E**) Dark absorbance spectrum of NOL14 (data from (*101*)). NOM6 and NOL14 spectra are fitted with a two component Gaussian Mixture Model (GMM) suggesting the existence of two probably pH-dependent isoforms. Data is presented as mean ± SEM.

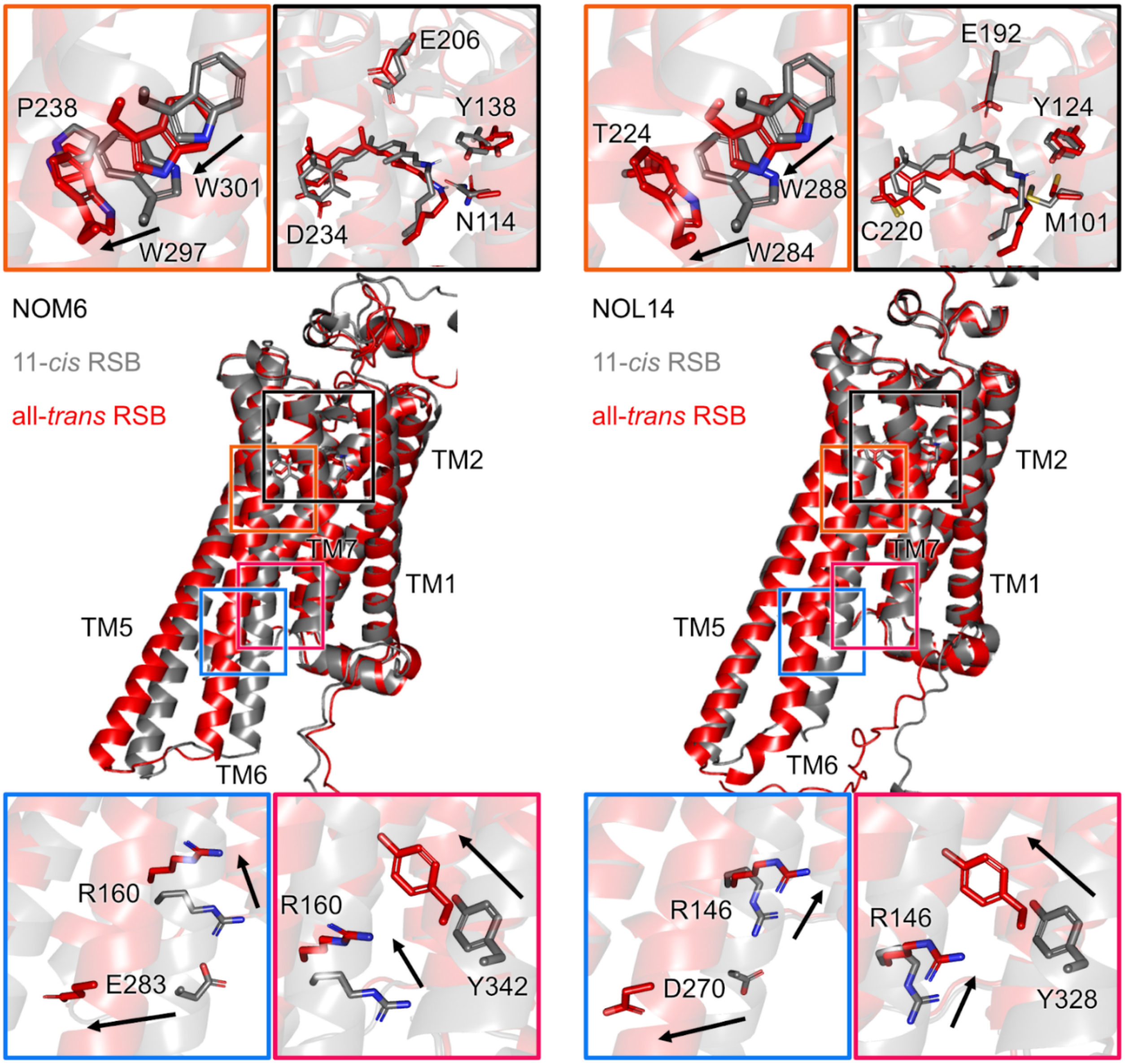


**Fig. S7.**

**Comparison of computed structural models between NOM6 and NOL14.** Inactive (11,*cis*-retinal Schiff base bound) and active (all-*trans* retinal Schiff base bound) states are shown as gray and red cartoons, respectively. In the active states, conserved motifs undergo similar rearrangement that distinguish them from the inactive states (*102*). The PIF motif contains a tryptophan substitution at position 6.44 adjacent to the CWxP tryptophan. Activation rearranges these residues (W297/W301 in NOM6 and W284/W288 in NOL14), reshaping the hydrophobic core, disrupting the DRY ionic lock (R160-E283 and R146-D270), and reorienting the NPxxY tyrosine (Y342 and Y328) toward the intracellular cavity.

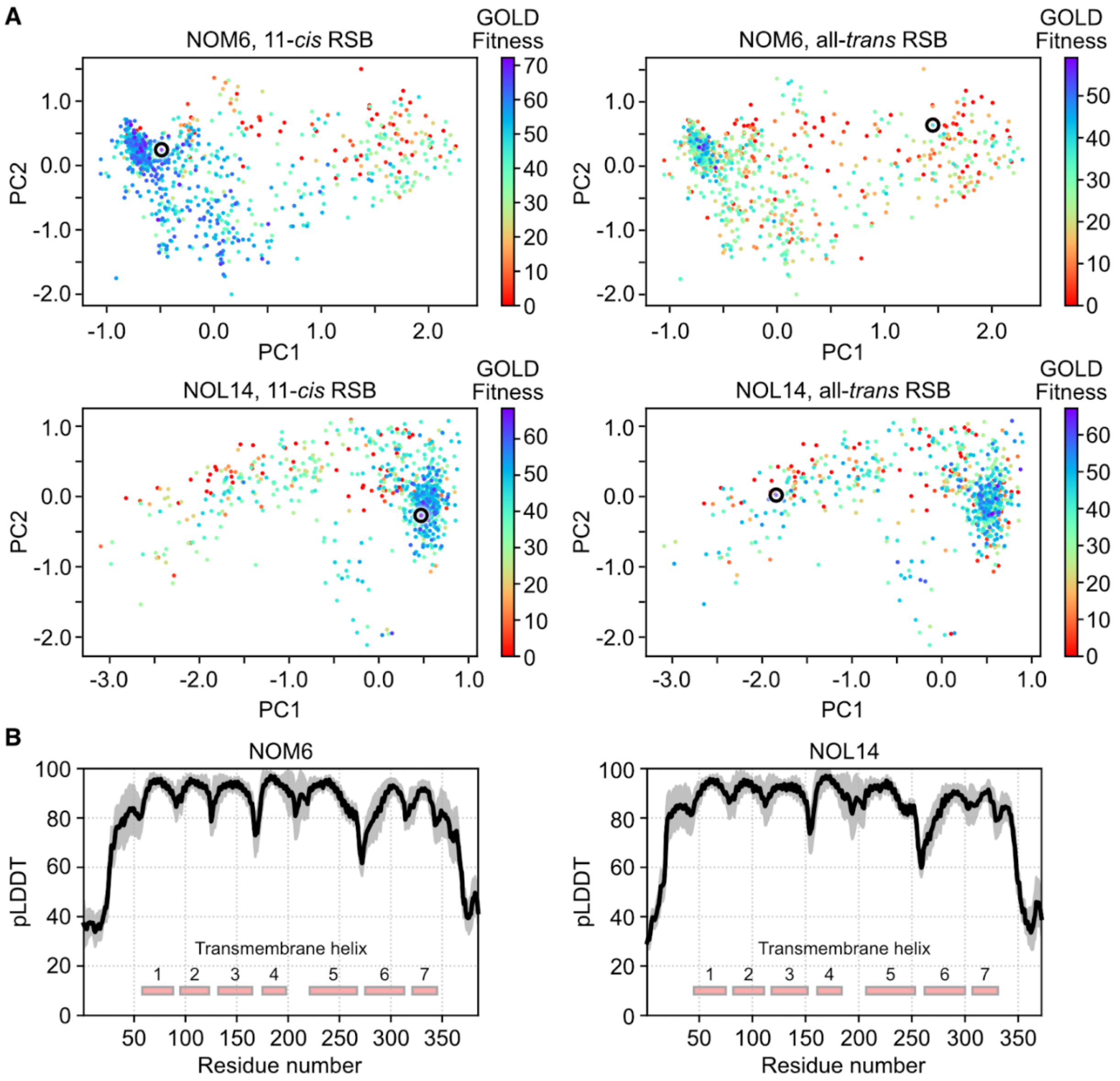


**Fig. S8.**

**Predicted conformational ensembles and evaluation of model accuracy.** (**A**) Principal component analysis (PCA) of NOM6 and NOL14 conformations, excluding the N- and C-terminal regions. Docking scores for the 11-*cis* and all-*trans* retinal Schiff bases are shown by color coding, and the conformations with negative scores were discarded. Selected inactive-like (left) and active-like (right) states are indicated by black circles. (**B**) Predicted local distance difference test (pLDDT) scores of the conformations. Transmembrane helices are shown as red rectangles. The black line represents the mean pLDDT of Cα atoms, and the gray shading indicates the standard deviation. The number of conformations analyzed was 885 for NOM6 and 810 for NOL14.

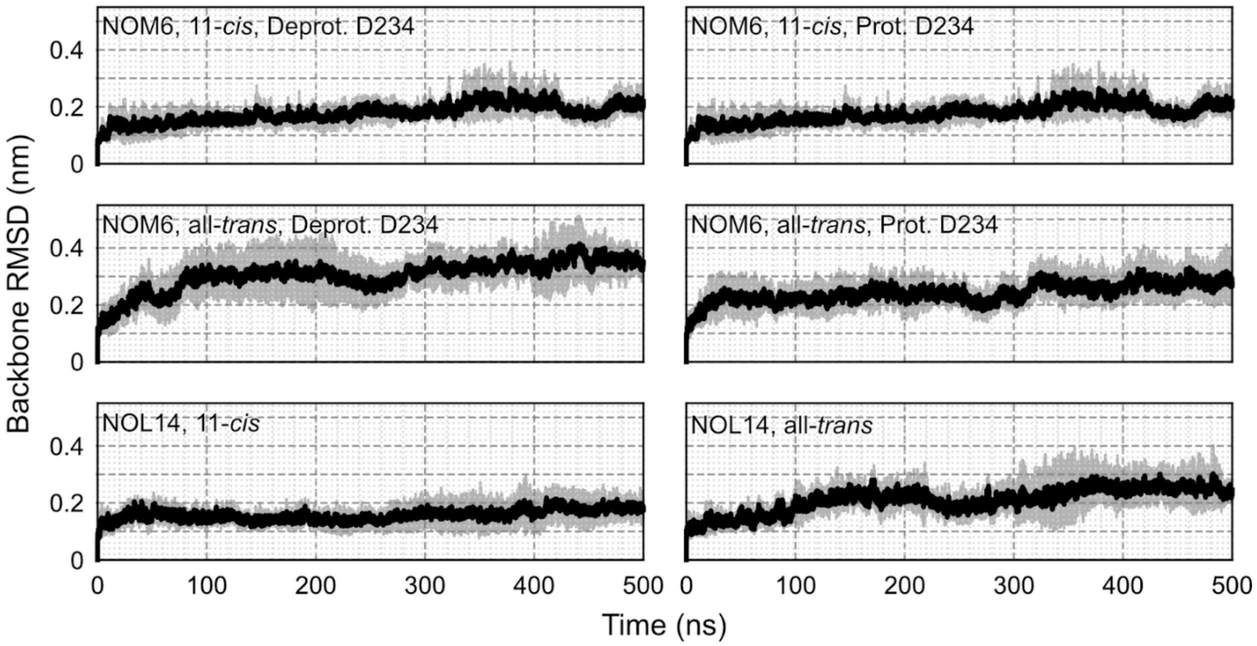


**Fig. S9.**

**Root mean square deviations (RMSDs) of NOM6 and NOL14.** Backbone RMSDs were calculated excluding the N- and C-terminal regions. The solid line represents the mean RMSD, and the shaded area indicates the standard deviation. Molecular dynamics simulations were performed for 500 ns and repeated three times.

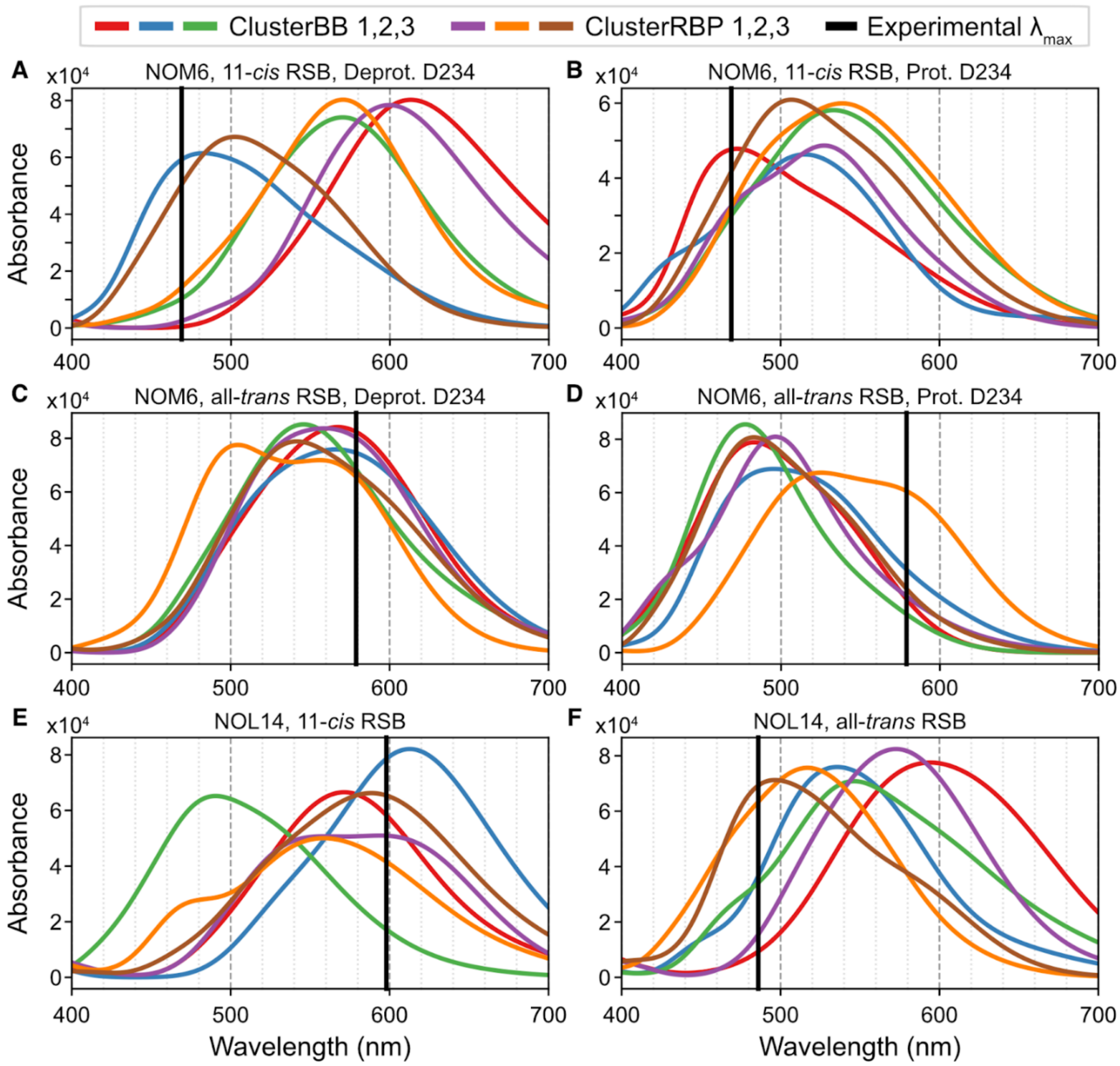


**Fig. S10.**

**Averaged UV-Vis spectra of NOM6 and NOL14.** Spectra calculated at ADC(2) / cc-pVDZ level from snapshots obtained using DFTB2 with dispersion correction. (**A-F**) The experimental mean absorption maximum is shown as a black solid line. Selected clusters are as follows: NOM6, 11-*cis* RSB with protonated D234 ($\lambda_{max}$ = 473 nm, ClusterBB 1 in (**B**); NOM6, all-*trans* RSB with deprotonated D234 ($\lambda_{max}$ = 566 nm, ClusterBB 2 in (**C**)); NOL14, 11-*cis* RSB ($\lambda_{max}$ = 594 nm, ClusterRBP 1 in (**E**)); and NOL14, all-*trans* RSB ($\lambda_{max}$ = 497 nm, ClusterRBP 3 in (**F**)). Note that NOM6 with 11-*cis* RSB and deprotonated D234 (**A**) and NOM6 with all-*trans* RSB and protonated D234 (**D**) were not selected, as their calculated spectral trends were inconsistent with the experimental observations.

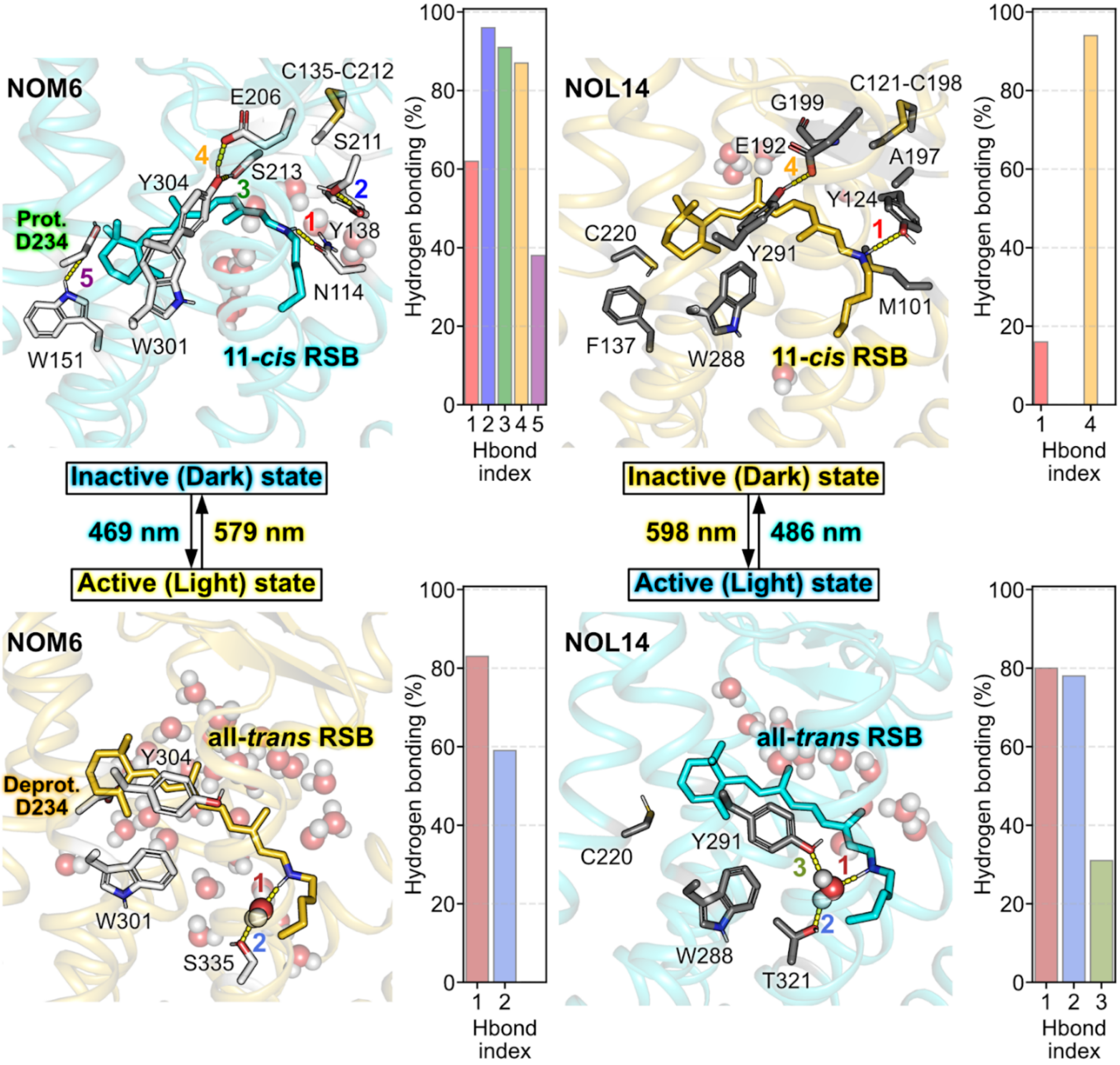


**Fig. S11.**

**Hydrogen bonding analysis of NOM6 and NOL14.** Percentage of hydrogen-bonding interactions were analyzed over 1-ns hybrid quantum mechanics / molecular mechanics (QM/MM) simulations using a distance cutoff of 3.5 Å between donor (D) and acceptor (A) atoms and an angle cutoff of 150° for the D-H-A geometry. Water molecules are shown as spheres.

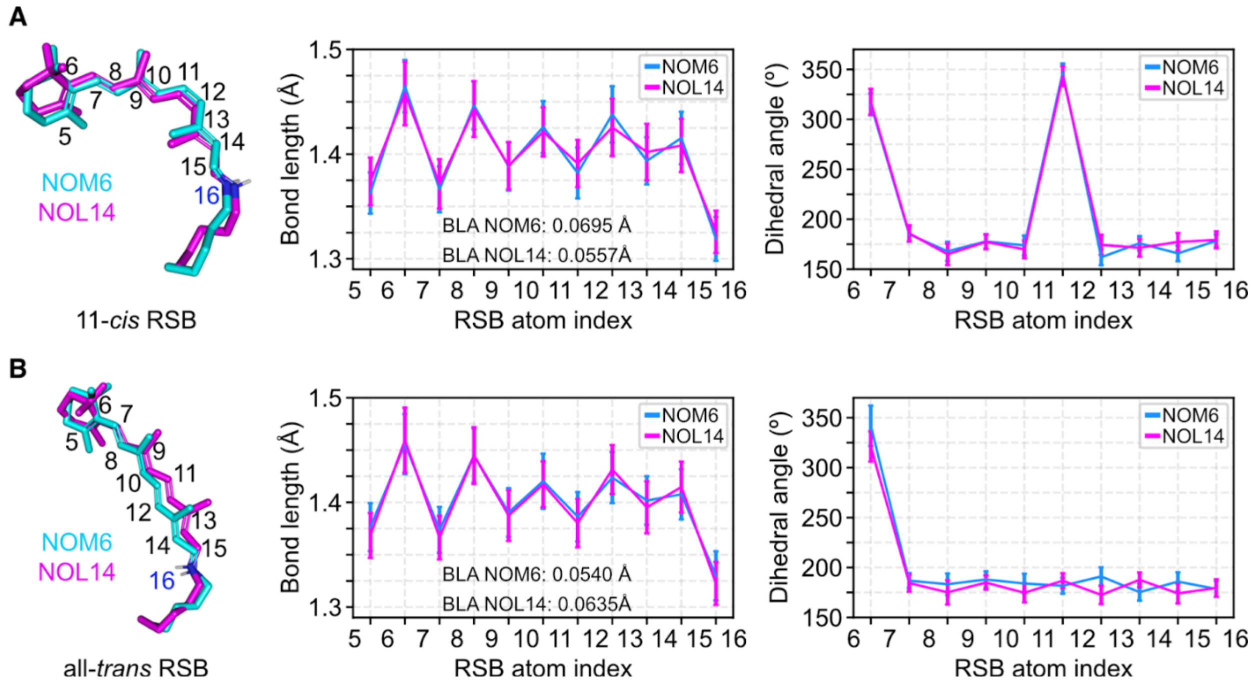


**Fig. S12.**

**Bond length and dihedral angle analysis.** Bond lengths and dihedral angles of (**A**) 11-*cis* and (**B**) all-*trans* retinal Schiff base bound in NOM6 (cyan) and NOL14 (magenta) were obtained from 1-ns hybrid quantum mechanics / molecular mechanics (QM/MM) simulations. The bond length alternation (BLA) was calculated as the difference between the average lengths of single and double bonds.

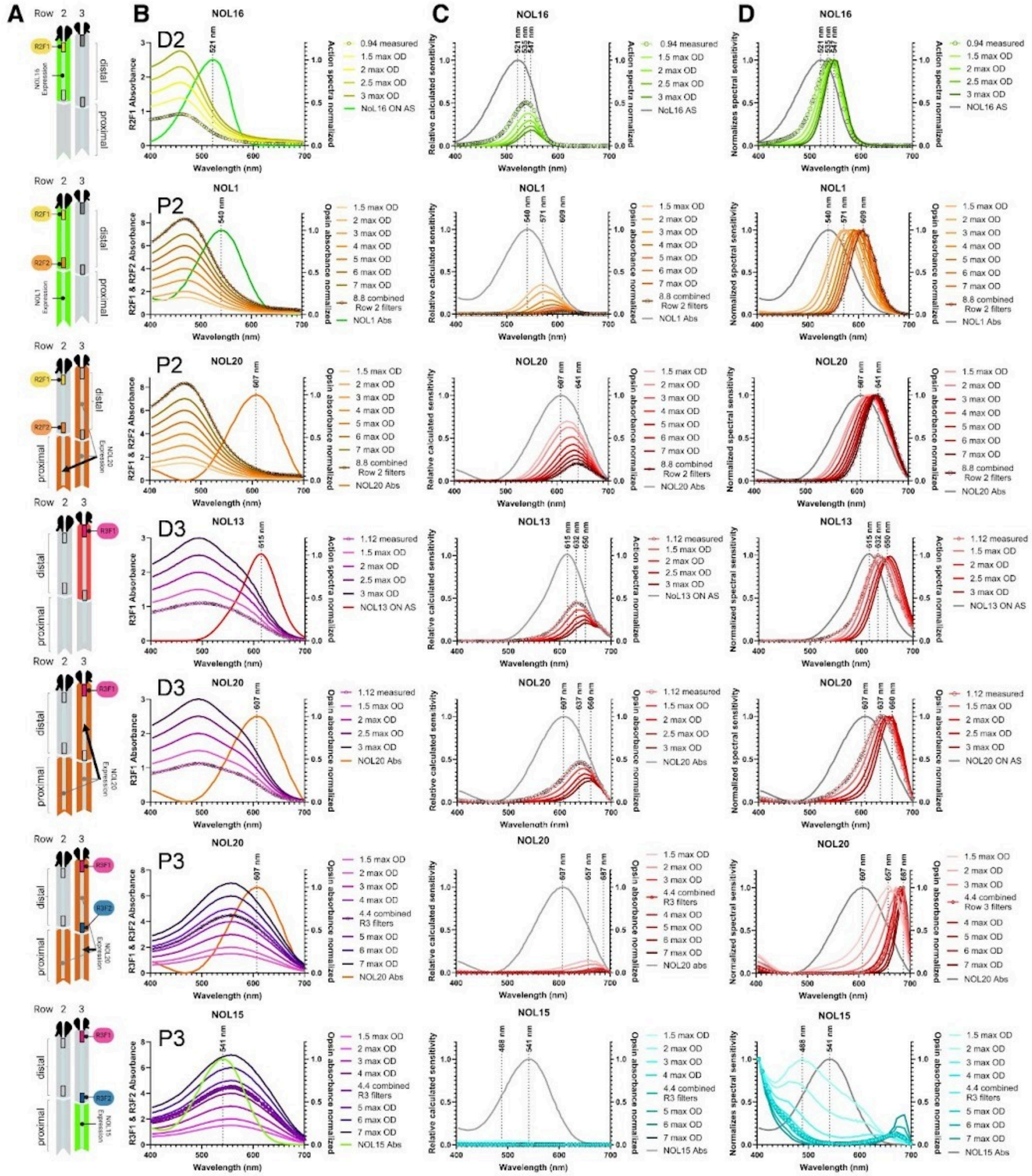


**Fig. S13.**

**Effect of intra-rhabdomal filter properties on the rhodopsin spectral sensitivities in tiers D2, P2, D3 and P3.** (**A**) Schematic representation of midband rows 2 and 3 including the rhodopsins and the filters relevant for the analysis presented in each row. (**B**) Normalized rhodopsin spectra and filter spectra of different optical density ranging from 1.5 to 3 for distal (D) filters, and ranging from 1.5 to 7 for combined filters in the proximal (P) tier. Digitized filter absorbances reported in the literature (*5*) are labelled by black circle symbols. (**C**) Calculated sensitivities of the indicated rhodopsins unfiltered (grey) and after filtering by screening pigments with maximal absorbance ranging from 1.5 to 3 for distal F1 filters, and ranging from 1.5 to 7 for proximal F2 filters. Numbers above vertical dashed/dotted lines represent, from left to right, the wavelength maxima before screening, after screening by filters of reported optical densities (OD), and after maximal screening when applicable (NOL15 is almost entirely shielded by intra-rhabdomal pigments and the low resulting sensitivity could not be fitted). (**D**) Normalized calculated sensitivity spectra plotted with original spectral sensitivity (grey). In all graphs, absorbances and sensitivities resulting from originally measured filters (*5*) are labeled by black open circle symbols.

A

| Clade | | Gq | | | | |
|---|---|---|---|---|---|---|
| | | *Average rel. luminescence* | *SEM* | *p-value* | *n* | *% of maximal response* |
| A | **NOL6** | **10.96** | **3.72** | **0.0296** | **5** | **2.5%** |
| A | **NOL7** | **12.38** | **3.76** | **0.0207** | **5** | **2.8%** |
| A | **NOL12** | **26.00** | **6.82** | **0.0338** | **3** | **5.8%** |
| A | **NOL8** | 1.33 | 0.13 | 0.3066 | 5 | 0.3% |
| | **LWS R3** | **26.79** | **8.34** | **0.0274** | **4** | **6.0%** |
| C | **NOL9** | 1.22 | 0.06 | 0.4211 | 4 | 0.3% |
| C | **NOL20** | **1.57** | **0.14** | **0.0328** | **5** | **0.4%** |
| D | **NOL1** | **4.29** | **1.20** | **0.0424** | **4** | **1.0%** |
| D | **NOL2** | 1.27 | 0.10 | 0.3958 | 4 | 0.3% |
| E | **NOL16** | **357.30** | **65.89** | **0.0163** | **3** | **80.0%** |
| E | **NOL17** | **127.60** | **28.86** | **0.0241** | **3** | **28.6%** |
| E | **NOL19** | **446.60** | **65.57** | **0.0105** | **3** | **100.0%** |
| F | **NOL13** | **33.51** | **4.04** | **0.0021** | **4** | **7.5%** |
| F | **NOL15** | 1.41 | 0.17 | 0.3929 | 5 | 0.3% |
| F | **NOL14** | **97.60** | **23.90** | **0.0078** | **5** | **21.9%** |
| I | **NOM5** | **342.30** | **34.43** | **0.0179** | **5** | **76.6%** |
| I | **NOM6** | **252.50** | **42.94** | **0.0179** | **5** | **56.5%** |

B

| Clade | | Gi/o | | | | |
|---|---|---|---|---|---|---|
| | | *Average rel. luminescence* | *SEM* | *p-value* | *n* | *% of Glosensor only* |
| A | **NOL6** | 0.74 | 0.01 | 0.0278 | 3 | 114.8% |
| A | **NOL7** | 0.70 | 0.03 | 0.2 | 3 | 109.5% |
| A | **NOL12** | 0.78 | 0.02 | 0.0153 | 3 | 120.7% |
| A | **NOL8** | 0.63 | 0.02 | 0.3803 | 3 | 98.8% |
| | **LWS R3** | 0.66 | 0.00 | 0.3542 | 3 | 102.8% |
| C | **NOL9** | 0.73 | 0.03 | 0.0805 | 3 | 113.2% |
| C | **NOL20** | 0.94 | 0.03 | 0.0645 | 3 | 92.9% |
| D | **NOL1** | 0.70 | 0.01 | 0.0961 | 3 | 108.7% |
| D | **NOL2** | 0.70 | 0.02 | 0.1273 | 3 | 108.7% |
| E | **NOL16** | 0.72 | 0.02 | 0.0701 | 3 | 112.0% |
| E | **NOL17** | 0.75 | 0.02 | 0.0289 | 3 | 116.1% |
| E | **NOL19** | **0.49** | **0.02** | **0.0081** | **3** | **76.8%** |
| F | **NOL13** | 0.73 | 0.01 | 0.0318 | 3 | 114.2% |
| F | **NOL15** | 0.73 | 0.05 | 0.1051 | 3 | 113.6% |
| F | **NOL14** | 1.00 | 0.01 | 0.3224 | 6 | 104.4% |
| I | **NOM5** | **0.73** | **0.01** | **<0.0001** | **3** | **73.8%** |
| I | **NOM6** | **0.79** | **0.02** | **0.0006** | **3** | **79.2%** |

**Table S1.**

**Gq- and Gi/o-coupled response characteristics measured in the aequorin and glosensor assay.** Significant Gq and Gi/o-responses are reported in bold. (**A**)**.** Maximal relative Gq-responses are expressed as a percentage of the highest Gq-signaling opsin (NOL19). (**B**)**.** Maximal relative Gi/o responses are reported for each opsin, as the percentage of response of cells transfected only with glosensor (Glosensor only), at the peak decrease in luminescence, between the forskolin-induced plateau and the estimated response decay (see Methods). Table S2.

| | Mixture | Gaussian 1 | | Gaussian 2 | | Light Intensity (photons.cm-².s-¹) |
|---|---|---|---|---|---|---|
| | $\lambda_{max}$ (nm) | $\lambda_{max}$ (nm) | percentage of mixture (%) | $\lambda_{max}$ (nm) | percentage of mixture (%) | |
| **NOL13** | 615 ± 5.6 | 618.9 | 55.9 | 572.6 | 44.1 | $1x10^{15}$ |
| **NOL14** | 592.4 ± 3.4 | 538.2 | 60.9 | 597.5 | 39.1 | $1x10^{15}$ |
| **NOM5** | 474.9 ± 8.2 | 449.6 | 65.3 | 487.3 | 34.7 | $2.4x10^{13}$ |
| **NOM6** | 469.1 ± 9.8 | 480.9 | 52.7 | 442.1 | 47.3 | $2.4x10^{13}$ |

**Table S2.**

**Dark state spectral characteristics of photoreversible opsins.** The percentage of mixture for the individual gaussian is reported as the percentage of the area under the curve of the combined GMM model.

| | $Conf_{RSB}$ | *Dim* | $N_{Na/Cl}$ | $N_{water}$ | $N_{POPC}$ |
|---|---|---|---|---|---|
| NOM6 (Prot. D234) | 11-*cis* | 9.7/7.7/13.0 | 75/78 | 27929 | 260 |
| | all-*trans* | 9.5/9.5/14.4 | 83/86 | 30744 | 258 |
| NOM6 (Deprot. D234) | 11-*cis* | 9.6/9.6/13.1 | 75/77 | 27935 | 260 |
| | all-*trans* | 9.6/9.6/13.9 | 83/85 | 30740 | 258 |
| NOL14 | 11-*cis* | 7.9/7.9/14.4 | 57/61 | 21396 | 173 |
| | all-*trans* | 8.0/8.0/13.5 | 53/58 | 19937 | 172 |

**Table S3.**

**System setup for molecular dynamics simulations.** $Conf_{RSB}$: configuration of retinal Schiff base, *Dim*: box size (X/Y/Z nm), $N_{Na/Cl}$: the number of $Na^+/Cl^-$, $N_{water}$: the number of water molecules, $N_{POPC}$: the number of POPC. All simulations were conducted at a NaCl concentration of 150 mM and at a temperature of 303 K and replicated three times.

| | $\Delta t$ | $t$ | $k_{bb}$ | $k_{sc}$ | $k_{scRSB}$ | $k_{head}$ | $k_{torsion}$ |
|---|---|---|---|---|---|---|---|
| Energy minimization | - | - | 4000 | 2000 | 2000 | 1000 | 1000 |
| Equilibration 1 | 1 | 0.25 | 4000 | 2000 | 2000 | 1000 | 1000 |
| Equilibration 2 | 1 | 0.25 | 2000 | 1000 | 2000 | 400 | 400 |
| Equilibration 3 | 1 | 0.25 | 1000 | 500 | 1000 | 400 | 200 |
| Equilibration 4 | 2 | 1 | 500 | 200 | 1000 | 200 | 200 |
| Equilibration 5 | 2 | 1 | 200 | 50 | 500 | 40 | 100 |
| Equilibration 6 | 2 | 1 | 50 | 0 | 500 | 0 | 0 |
| Equilibration 7 | 2 | 100 | 0 | 0 | 200 | 0 | 0 |

**Table S4.**

**Step-wise release of restraint of applied positional and dihedral angular restraints for energy minimization and equilibration simulations.** Time step: $\Delta t$ (fs), individual simulation time: $t$ (ns), force constant (kJ/mol/nm) for position/angle harmonic restraints of backbone heavy atoms ($k_{bb}$), side-chain heavy atoms ($k_{sc}$), side-chain heavy atoms of retinal Schiff base ($k_{scRSB}$), lipid head group ($k_{head}$), and lipid chirality and *cis* double bond ($k_{torsion}$).

| | *RSB configuration* | *BB 1* | *BB 2* | *BB 3* | *RBP 1* | *RBP 2* | *RBP 3* |
|---|---|---|---|---|---|---|---|
| NOM6 (Prot. D234) | 11-*cis* | 95.1 | 2.8 | 1.8 | 72.2 | 27.1 | 0.4 |
| | all-*trans* | 76.9 | 12.6 | 9.8 | 98.5 | 0.8 | 0.7 |
| NOM6 (Deprot. D234) | 11-*cis* | 73.6 | 15.0 | 7.5 | 61.1 | 22.6 | 10.1 |
| | all-*trans* | 40.0 | 17.0 | 15.1 | 34.8 | 25.7 | 14.5 |
| NOL14 | 11-*cis* | 92.6 | 4.0 | 3.0 | 98.3 | 1.3 | 0.4 |
| | all-*trans* | 97.8 | 1.8 | 0.4 | 95.5 | 2.6 | 1.8 |

**Table S5.**

**Cluster population (%) of ClusterBB and ClusterRBP.** Cluster populations of ClusterBB (1-3) and ClusterRBP (1-3) were derived from 500-ns molecular dynamic simulations with three independent replicates. The percentage values represent the relative abundance of each conformational cluster obtained from trajectory-based structural clustering analysis.

**Data S1. (separate file)**

Sequences used in the alignment of fig. S1, with bovine rhodopsin as reference (PDB: 1GZM_A). Sequences were obtained from previous publications *(29, 30).*
